# Local fitness and epistatic effects lead to distinct patterns of linkage disequilibrium in protein-coding genes

**DOI:** 10.1101/2021.03.25.437004

**Authors:** Aaron P. Ragsdale

## Abstract

Selected mutations interfere and interact with evolutionary processes at nearby loci, distorting allele frequency trajectories and creating correlations between pairs of mutations. A number of recent studies have used patterns of linkage disequilibrium (LD) between selected variants to test for selective interference and epistatic interactions, with some disagreement over interpreting observations from data. Interpretation is hindered by a lack of analytic or even numerical expectations for patterns of variation between pairs of loci under the combined effects of selection, dominance, epistasis, and demography. Here, I develop a numerical approach to compute the expected two-locus sampling distribution under diploid selection with arbitrary epistasis and dominance, recombination, and variable population size. I use this to explore how epistasis and dominance affect expected signed LD, including for non-steadystate demography relevant to human populations. Using whole-genome sequencing data from humans, I explore genome-wide patterns of LD within protein-coding genes. I show that positive LD between missense mutations within genes is driven by strong positive allele-frequency correlations between pairs of mutations that fall within the same annotated conserved domain, pointing to compensatory mutations or antagonistic epistasis as the prevailing mode of interaction within conserved genic elements. LD between missense mutations is reduced outside of conserved domains, as would expected under Hill-Robertson interference. This variation in both mutational fitness effects and selective interactions within proteincoding genes calls for more refined inferences of the joint distribution of fitness and interactive effects, and the methods presented here should prove useful in that pursuit.

## 1 Introduction

Most new mutations that affect fitness are deleterious and tend to be eliminated from a population. The amount of time that a deleterious mutation segregates depends on the strength of selection against genomes that carry it, with very damaging mutations kept at low frequencies and purged relatively rapidly. In the time between mutation and fixation or loss, selected variants, both beneficial and damaging, can dramatically impact patterns of variation in nearby linked regions (e.g., Smith and Haigh, 1974; Charlesworth *et al*., 1995; Kim and Stephan, 2000). This distortion away from neutral expectations has been empirically documented using sequencing data from an ever-growing set of study systems (Novembre and Di Rienzo, 2009; Cutter and Payseur, 2013; Comeron, 2014), but questions remain about the primary mode of interactions between multiple linked variants and their joint effects on genome-wide patterns of diversity.

In their foundational paper, Hill and Robertson (1966) recognized that linked selected variants reciprocally impede the efficacy of selection at each locus, a process known as selective interference. Linked selection reduces the fixation probability of advantageous mutations and increases that of deleterious mutations compared to expectations under single-locus models (Birky and Walsh, 1988). Allele frequency dynamics and correlations of linked selected variants are also predicted to deviate from single-locus expectations. Under a multiplicative fitness model, where the fitness reduction of a genome carrying multiple deleterious variants is equal to the product of the fitness reduction of each independent mutation, we expect to see net linkage disequilibrium (LD) equal to zero for unlinked sites (Kondrashov, 1995). But for linked loci, those mutations are expected to segregate on different haplotypes more often than together, leading to negative, or repulsion, LD, although the extent of LD depends non-trivially on the strength of selection and the probability of recombination separating loci (Hill and Robertson, 1966; McVean and Charlesworth, 2000).

Non-additive effects, including dominance (i.e., interactions *within* a locus) and epistasis (interactions *between* loci), further complicate our evolutionary models. A large fraction of nonsynonymous coding mutations are thought to be at least partially recessive (Agrawal and Whitlock, 2011; Huber *et al*., 2018), with average levels of dominance correlating with strength of selection (Kacser and Burns, 1981), and dominance plays an important role in shaping expected equilibrium allele frequencies and the mutation load of strongly damaging disease mutations (Clark, 1998). On the other hand, epistasis differentially impacts the deleterious load in asexually and sexually reproducing organisms (Kimura and Maruyama, 1966; Kondrashov, 1995), has been invoked as an explanation for the evolutionary advantage of sex (Kondrashov, 1982; Charlesworth, 1990; Barton and Charlesworth, 1998), and can drive incompatibilities that lead to postzygotic isolation during the process of speciation (Turelli and Orr, 2000). Within populations, epistasis is known to cause signed LD to deviate dramatically from zero (Charlesworth, 1990; Kondrashov, 1995). However, despite appreciation of the effect of dominance on linked variation (Turelli and Orr, 2000; Zhao and Charlesworth, 2016) and the evolutionary importance of epistatic interactions, we currently lack models for predicting patterns of correlations between linked mutations under general selection models.

In this paper, I develop a numerical approach to solve for the two-locus sampling distribution under a general diploid selection model with variable recombination and single-population size history. I use this model to describe how epistasis and dominance shape expected patterns of signed LD, under both steady-state and non-equilibrium demography, that have been used to test for interference and epistasis in population genomic data. I then turn to human sequencing data and compare patterns of LD for synonymous, missense, and loss-of-function mutations in protein-coding genes and annotated conserved domains. I show that while synonymous and missense variants display similar slightly positive average LD within genes, for missense mutations this signal is driven by correlations between pairs of mutations within, but not between or outside of, protein-coding domains. This suggests an importance for antagonistic epistasis or a prevalence of compensatory nonsynonymous mutations within conserved elements.

### 1.1 Empirical observations

The most direct way to test for interactions between linked selected variants is through deep mutation scanning experiments, in which many distinct mutations are introduced within a target gene and then organismal fitness or some protein function is experimentally measured (Romero and Arnold, 2009; Bank *et al*., 2015; Puchta *et al*., 2016; Steinberg and Ostermeier, 2016). For example, using the model system of the TEM-1 *β*-lactamase gene in *E. coli*, Bershtein *et al*. (2006) found evidence for synergistic epistasis, in which the combined effect of multiple deleterious mutations on individual fitness was larger than would be expected from multiplying the independent observed effects of each individual mutation. The scale of mutation scanning experiments continues to increase, promising greater resolution of the fitness landscape in such model systems that can be compared to evolutionary theory (Otwinowski *et al*., 2018).

Directed mutational studies are not possible in most natural populations, and we must turn to population genetic approaches to infer selective interactions between observed segregating polymorphisms. Motivated by theoretical predictions that linked negatively selected mutations will display negative LD due to interference (Hill and Robertson, 1966) and that epistasis will drive expected LD away from zero, a number of recent studies have used patterns of LD within classes of putatively selected variants to infer modes of selective interactions. Callahan *et al*. (2011) observed that pairs of tightly linked nonsynonymous mutations cluster together more than expected along the same lineages in the *Drosophilid* species complex, and that those clustered mutations tend to preserve the charge of the protein and were in positive LD compared to pairs of synonymous mutations at the same distance. From this, they proposed that compensatory nonsynonymous variants are regularly tolerated and maintained. More recently, Taverner *et al*. (2020) replicated this finding across a diverse set of genera, showing that such epistatic interactions are important for protein evolution.

Sohall *et al*. (2017) observed negative LD between loss-of-function variants in protein-coding genes (loss-of-function mutations include stop gains and losses, frameshifts, and other nonsense mutations) in both human and fruit fly populations. This was interpreted as evidence for widespread synergistic epistasis between these mutations, in which the fitness reduction of multiple mutations is greater than the product of that of each individual mutation independently. Both Sandler *et al*. (2021) and Garcia and Lohmüeller (2021) have recently reevaluated patterns of LD between coding variants in humans, fruit flies, and *Capsella grandiflora*, and suggested interference and dominance may instead be driving patterns of LD (Garcia and Lohmüeller, 2021) or questioned whether LD between loss-of-function variants is significantly different from zero (Sandler *et al*., 2021).

A number of factors impede our interpretation of patterns of signed LD between coding variants. First, for strongly deleterious or loss-of-function mutations, their low allele frequencies mean that measurements of LD and other diversity statistics are very noisy. Second, comparisons are based on theory with limiting assumptions, such as steady-state demography, simple selection and interaction models, or unlinked loci. To generate predictions under more complex models, we rely on expensive forward simulations. Such simulations can help build intuition and be used to test inference methods, but they do not efficiently provide expectations for quantities of interest across the range of relevant parameters. Analytical and numerical methods for expected haplotype frequencies and LD under general selective interaction models are thus crucial for interpreting patterns of variation observed in data.

## 2 Results

### 2.1 Expected signed LD under steady-state demography

In the Methods, I expand on the moments system developed in Ragsdale and Gravel (2019) to compute the expected sampling distribution of two-locus haplotypes (ψ_*n*_, Figure 1) under a general model of selective interactions. This sampling distribution stores the expected density or observed counts of pairs of biallelic loci with each possible haplotype frequency configuration in a sample of size *n*. Below, we compute expectations for ψ_*n*_, under varying scenarios of selection and interaction between pairs of loci. It is not possible in this framework to include the effects of additional linked selected mutations, such as background selection due to many linked variants, and individual-based forward simulations are still needed for such scenarios (e.g., Figures S7–S10).

**Figure 1:**
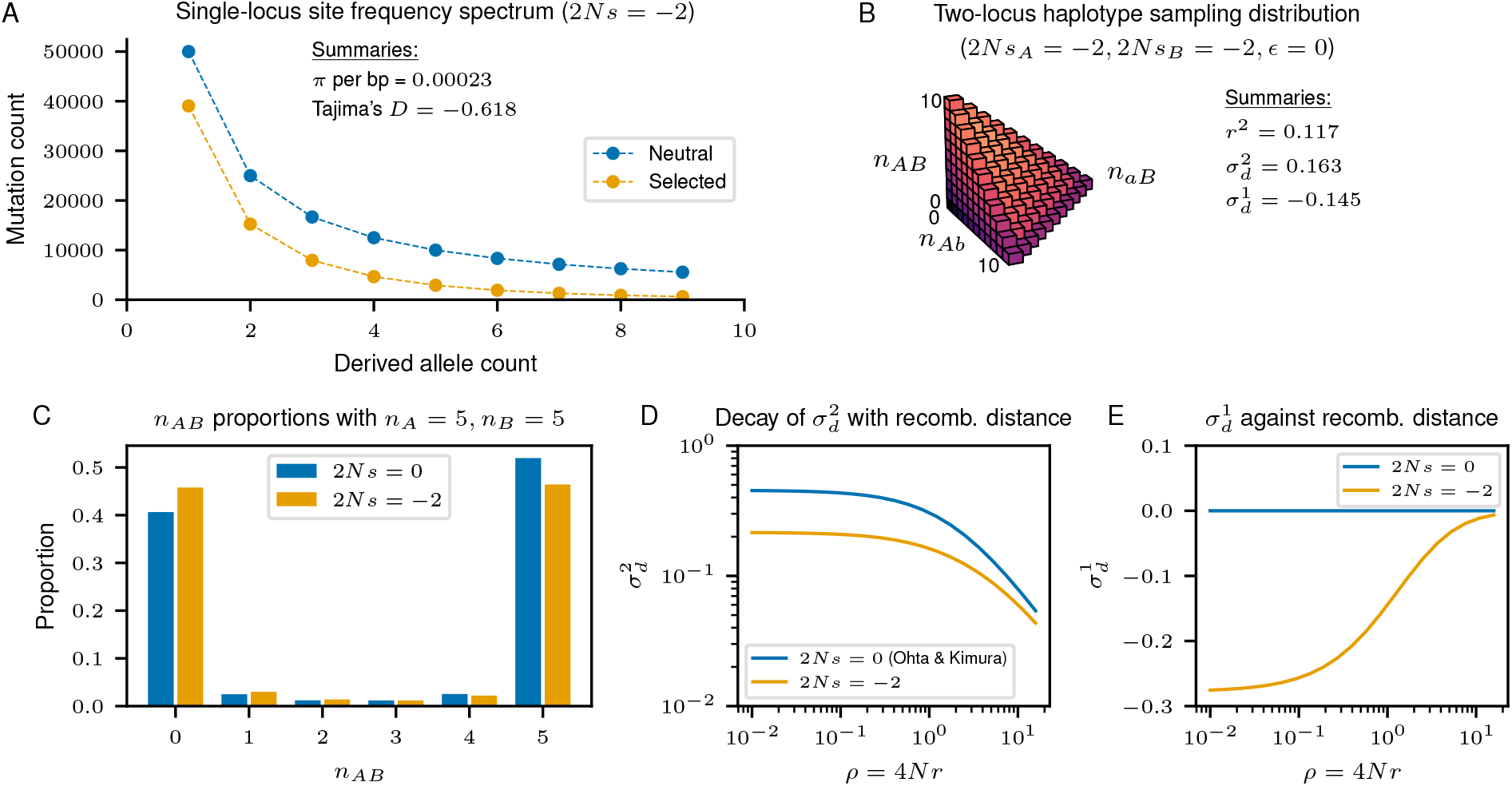
Sampling distributions and their summaries. Low-order summaries of sampling distributions are commonly computed for allele frequencies (A, the site-frequency spectrum, SFS) and two-locus haplotype distributions (B, linkage disequilibrium, LD). Demographic and selective processes affect both the SFS and LD, and observations of nonzero values of Tajima’s *D* or signed LD 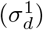 are often taken as evidence for selection or interactions between loci, respectively. (C) The full two-locus haplotype sampling distribution is a three-dimensional object, making it difficult to visualize. We can instead visualize conditional distributions of the full sampling distribution, e.g., conditioned on observing *n_A_* copies of the *A* allele at the left locus and *n_B_* copies of *B* at the right locus (e.g., Hudson, 2001). (D) 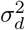, which is closely related to *r*^2^, is expected to decay with increasing recombination distance between loci (Ohta and Kimura, 1969). Selection distorts squared LD away from neutral expectations. (E) 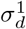, refered to as signed LD here, is expected to be zero for pairs of neutral mutations (Hill and Robertson, 1968). Interference between linked selected mutation causes negative signed LD (Hill and Robertson, 1966), and other forms of interactions between selected mutations can cause large negative or positive signed LD.

In many cases it is simpler to visualize summaries such as the expectation or variance of *D* (Figure 1D, E) or conditional slices of the distribution (Figure 1C) instead of the full three-dimensional distribution ψ_*n*_. Here I focus on low-order LD statistics including 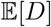 and 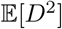 and their decay with recombination distance, as these are statistics that are commonly used to test for interactions between loci. For pairs of biallelic loci, with alleles labeled *A*/*a* at the left locus and *B*/*b* at the right locus, *D* = *f_AB_* – *f_A_f_B_* is the standard covariance measure of LD, where *f_AB_* is the frequency of haplotypes carrying both *A* and *B*, and *f_A_* and *f_B_* are the marginal allele frequencies of A and B.

Instead of unnormalized 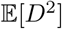 and 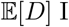 consider expectations for 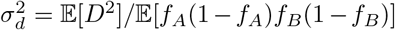 and 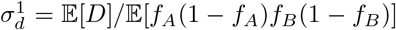. Such normalized statistics have two benefits: first, the mutation rate cancels so that expectations are robust to assumptions about the per-base mutation rate, and second, we can compare to analytic expectations for these quantities under neutrality and constant population size (Ohta and Kimura, 1971).

Below, I first consider the case of additive selection, Hill-Robertson interference, and epistasis. I then explore the effect of dominance acting within loci but without epistatis, and then describe a general diploid selection model and consider gene-based dominance effects. In what follows I focus on parameters where the strengths of selection and dominance at each locus are equal, but note that the methods presented here allow for arbitrary and unequal selection and dominance at the two loci. I also focus primarily on weak to moderate negative selection (|2*Ns*| ≈ 1 – 20), since this range of selection leads to the strongest signals of interference (Figure 2) and is the parameter regime for which the numerical appraoch is most accurate.

**Figure 2:**
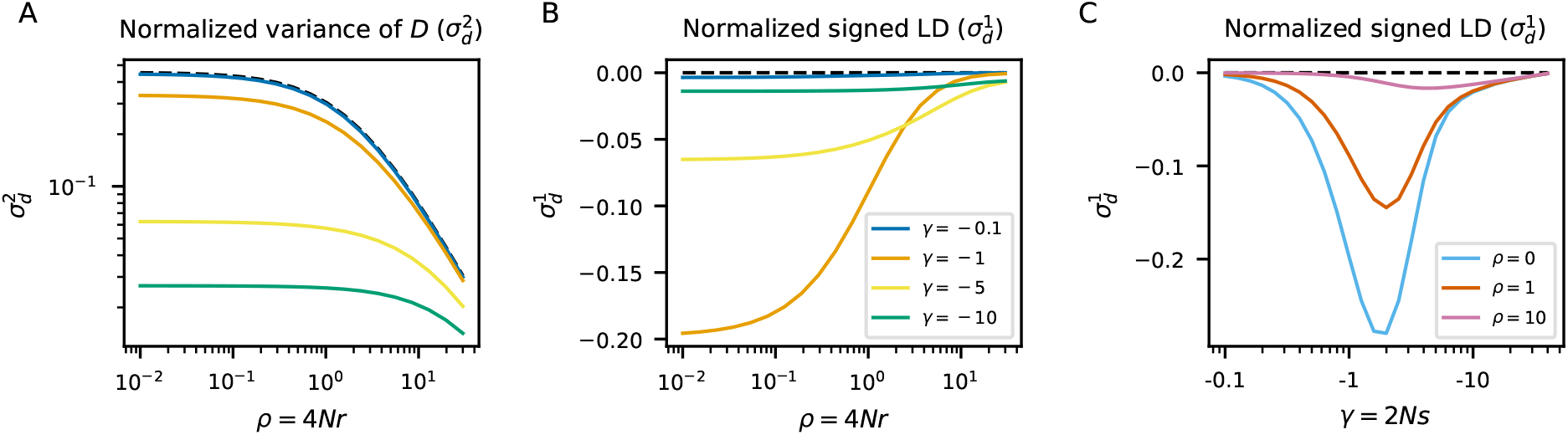
Hill-Robertson interference. Interference between pairs of selected mutations can cause negative signed LD in the absence of dominance and epistatic effects. (A) The expected normalized variance of 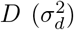 decreases below neutral expectations as the strength of negative selection increases. (B, C) For tightly linked loci (4*Nr* ≲ 1), interference is most noticeable for pairs of mutations with *s* ≈ 1/*N*. At larger recombination distances (4*Nr* > 1), signed LD is most negative for somewhat stronger selection coefficients. Dashed lines show neutral expectations.

#### 2.1.1 Additive selection and epistasis

For mutations under additive selection (*h* = 1/2) and no epistasis, we recover the well known Hill and Robertson (1966) interference result of negative LD between selected mutations, which is strongest for pairs of mutations that have selection coefficients *γ* = *O*(1), or *s* ≈ 1/2*N*_e_(Figure 2). For strongly deleterious mutations, LD is close to zero even with tight linkage, as they almost always segregate at low enough frequencies that they are unlikely to interfere with each other (McVean and Charlesworth, 2000).

With epistasis, mean signed LD is large for both weakly and strongly selected variants, with sign depending on the direction of epistatic interactions (Figure 3). Synergistic epistasis (in which the effect of two mutations together is larger than the product of each individual mutation’s effect) results in negative LD while antagonistic epistasis (in which the combined effect is less than the product of independent effects) results in positive LD, and large nonzero LD can occur even when epistasis is relatively weak. Epistasis-induced LD can extend over long distances, especially for strongly deleterious mutations. For example, in Figure 3F even moderately deleterious mutations with population-size-scaled selection coefficients of γ = −10 show large mean LD that extends to values of *ρ* much greater than 1 (for humans, assuming roughly 1 cM/Mb, this is on the order of 100 kilobases or more). More strongly deleterious interacting mutations are expected to show large signed LD over much larger recombination distances.

**Figure 3:**
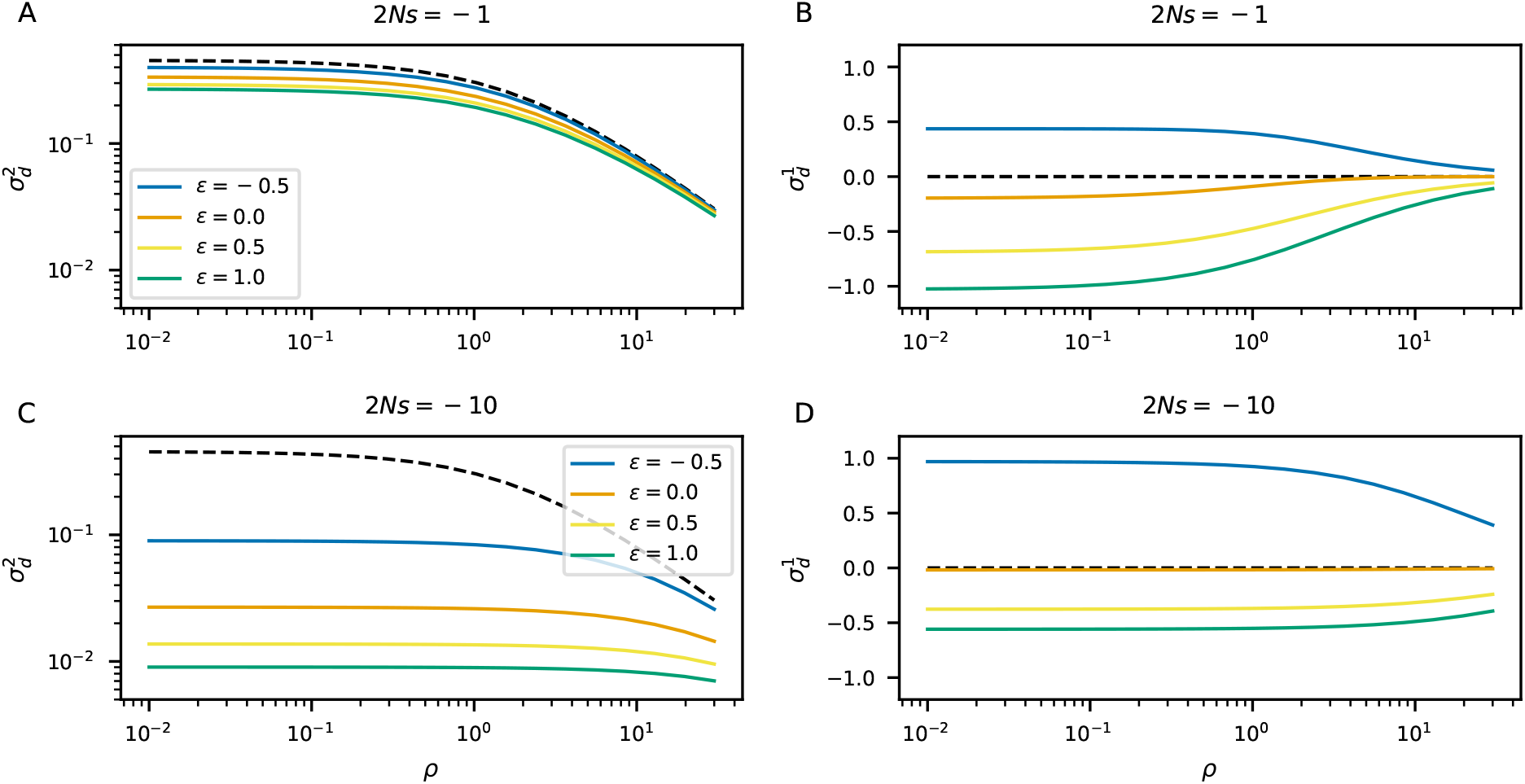
Additive selection and epistasis. Left panels (A and C) show expectations for the decay of 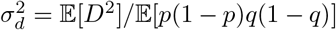 with recombination distance, and right panels (B and D) show expectations for the decay of 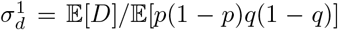. Dashed lines show neutral expectations. For both weak (*s* = –1/2*N*, A and B) and moderate (*s* = – 10/2*N*, C and D) selection, antagonistic epistasis (*ϵ* < 0) gives rise to positive signed LD and increased 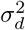 over a multiplicative model (*ϵ* = 0), and synergistic epistasis (*ϵ* > 0) results in negative signed LD beyond that of Hill-Robertson interference and decreased 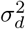.

#### 2.1.2 Dominance

The effect of non-additive selection on correlations between mutations has received increased attention recently. For example, Garcia and Lohmueller (2021) used large-scale forward simulations to explore how dominance impacts patterns of LD, showing that LD depends on the magnitudes of both the selection and dominance coefficients in a nonlinear way. Roze (2021) found an analytic expression for LD between pairs of strongly deleterious mutations under steady-state demography, showing that LD can be either positive or negative depending on the strength of dominance.

The combined effect of the strength of selection and dominance on interference is indeed nontrivial (Figure 4A), as observed by Garcia and Lohmüeller (2021). Some parameter regimes can cause strong negative LD between pairs of negatively selected variants, with moderately selected recessive variants having stronger signals of interference than additive selection (Figure 4B). Unlike epistatic interactions, signed LD decays rapidly with increasing distance between loci and is roughly zero for *ρ* ≫ 1. For weakly selected mutations with γ = –1, there is no monotonic effect of the level of dominance on negative LD, with both recessive and dominant pairs of mutations having more negative LD than the additive case.

**Figure 4:**
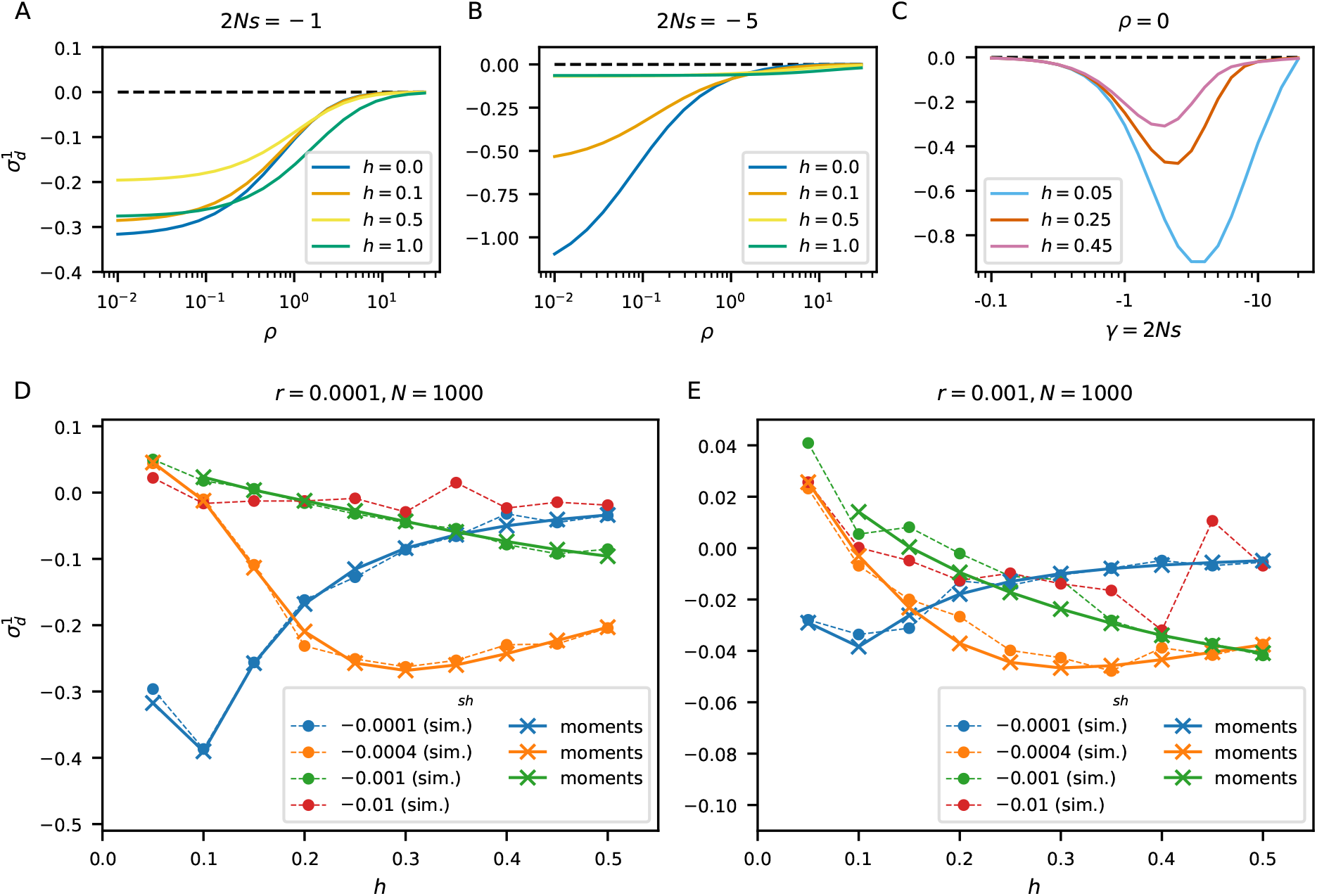
The effect of dominance on LD. (A, B) The strengths of selection and dominance interact in a nonlinear way to shape expected signed LD. For weakly to moderately selected mutations, as shown here, signed LD can be large and negative for tightly linked loci (e.g., *γ* = –5, *h* = 0, and *ρ* < 1). However, this large signed LD decays with recombination distance faster under a model of recessivity than does signed LD under a model of additive selection and epistasis (Figure 3). (C) Interference effects are most pronounced for recessive deleterious variants. (D, E) Recessive strongly deleterious mutations can have positive signed LD, as recently shown by Roze (2021). However the dominance threshold at which 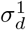 switches from positive to negative depends on the strength of selection, and weakly selected mutations can show non-monotonic behavior as *h* varies. Selection parameters of *sh* = –0.1 imply extremely strong selection (*h* = 0.05 results in *s* = –0.2 and *γ* = –400 for homozygous diploids at a single locus). The numerical appraoch for ψ_*n*_ cannot handle such strong selection.

Discrete simulations and the moments approach confirm the result from Roze (2021), that positive LD can occur for strong negative selection and small values of *h* (Figure 4D, E). However, while the analytic formula in Roze (2021) predicts that LD should be positive for *h* < 0.25 and negative for *h* > 0.25, this appears to only hold in the limit of strong selection (compare to Figure 1A in Roze (2021)). For moderate to moderately strong selection, this threshold of *h* can be less than 0.25, and LD is negative for all 0 ≤ *h* ≤ 1 for weakly deleterious mutations with interference strongest between pairs of partially recessive mutations (Figure 4C).

#### 2.1.3 General selection and gene-based dominance

Beyond the standard models of epistasis and dominance shown above, a large family of selection models can be specified by assigning unique fitness effects to each possible diploid pair of haplotypes. If we assume the diploid genotype that is homozygous for the ancestral alleles (*ab*/*ab*) has fitness 1, then there are nine other possible diploid two-locus genotypes that could be given unique fitnesses (Table S1), noting that *AB*/*ab* and *Ab*/*aB* genotypes can have differing selection coefficients.

The case with *s_AB/ab_* ≠ *s_Ab/aB_* arises in a scenario where a mutation at either locus within a haplotype impacts some functional region or element, but a diploid individual carrying at least one copy that is free of mutations has minimal fitness loss. In this “gene-based dominance” scenario (e.g., Sanjak *et al*., 2017), an *AB*/*ab* genotype has higher fitness than an *Ab*/*aB* type (a simple implementation of this model is given in the final column in Table S1). Such a gene-based fitness model gives similar expected positive signed LD to the model of antagonistic epistasis (Figure 5), although the interpretation of those two models can differ. With a highly parameterized space of possible general diploid selection models, multiple models with different biological interpretations can give similar patterns of expected signed LD.

**Figure 5:**
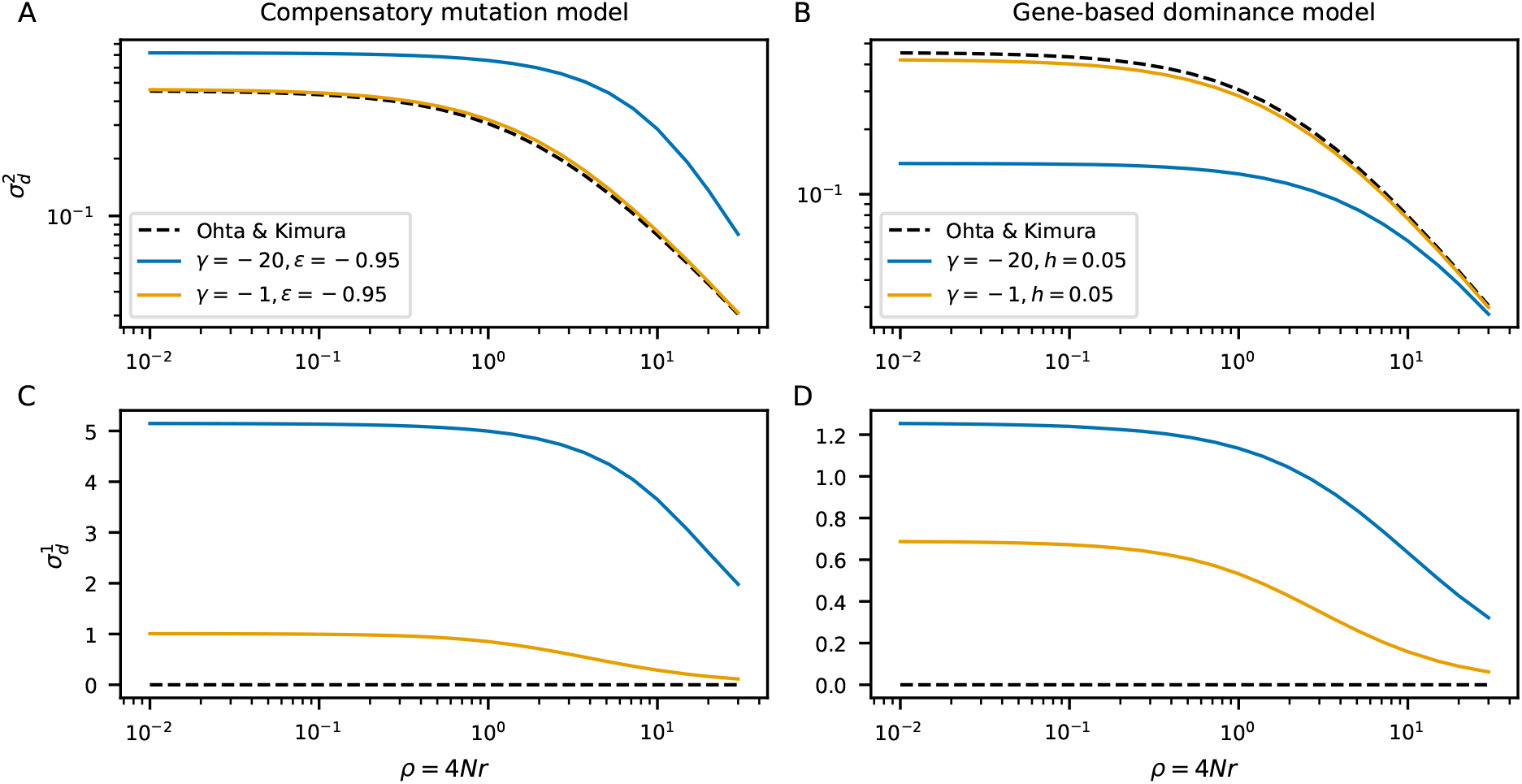
Multiple modes of interactions can lead to large positive signed LD. Both antagonistic epistasis (A and C, and which includes compensatory mutation models) and gene-based dominance (B and D) lead to large positive signed LD. Compensatory mutations (*ϵ* ≈ −1) also cause increased 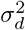 compared to neutral expectations (dashed black lines), while weaker antagonistic epistasis does not increase 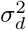 above neutral expectations (compare to Figure 3C). Gene-based dominance instead causes lower 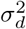 than neutral expectations. While signed LD may be similar between different interaction models, other two-locus summaries of the data may help to distinguish between such models.

### 2.2 The effect of population size changes on signed LD

The moment system for ψ_*n*_ readily incorporates variable population size as well as mutation and recombination rates and selection parameters that change over time. Here I focus on non-equilibrium population size history and consider scenarios relevant to human demographic history, including bottlenecks and expansions. I explore two simple models (Figure 6A), one with an instantaneous expansion and another with a bottleneck followed by recovery. I also consider two demographic histories inferred using genome-wide gene genealogy reconstruction (Speidel *et al*., 2019) applied to the 1000 Genomes Project Consortium *et al*. (2015) dataset, and focus on size histories for the Yoruba from Ibidan, Nigeria (YRI) and Utahns of North and West European ancestry (CEU) (Figure 7A).

**Figure 6:**
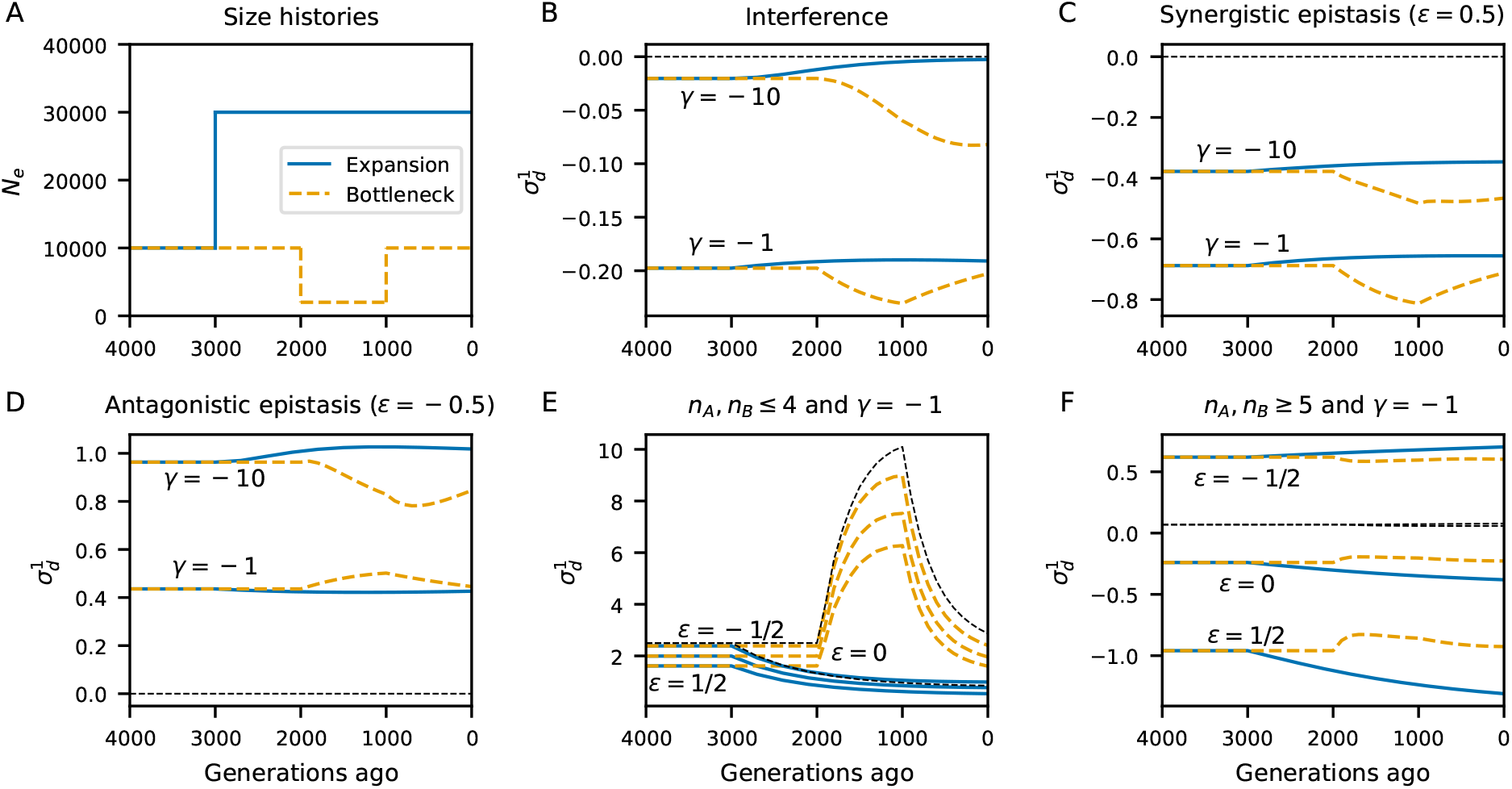
The effects of demography on signed LD. Simulations under two toy models, one with an instantaneous expansion in the past and the other with a five-fold reduction and recovery illustrated in (A), show the effects of population size changes on expected patterns of LD. For two selection strengths (γ = –1 and γ = –10) and three cases of interactions (non-epistatic interference (*ϵ* = 0, B), synergistic epistasis (*ϵ* = 1/2, C), and antagonistic epistasis (*ϵ* = –1/2, D), each with ρ = 0), sudden decreases in population size can cause large changes in signed LD, often in the opposite direction than more subtle shifts due to instantaneous expansion events. (E) Signed LD between pairs of rare alleles (both *n_A_*, *n_B_* ≤ 4, with sample size *n* = 50) is more sensitive to population size changes. (F) Signed LD between pairs of common alleles (both *n_A_*, *n_B_* ≥ 5/50) is comparatively more stable over time. Additional comparisons, including for dominance models and showing 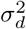, are shown in Figures S11–S13. Dashed lines indicate neutral expectations.

**Figure 7:**
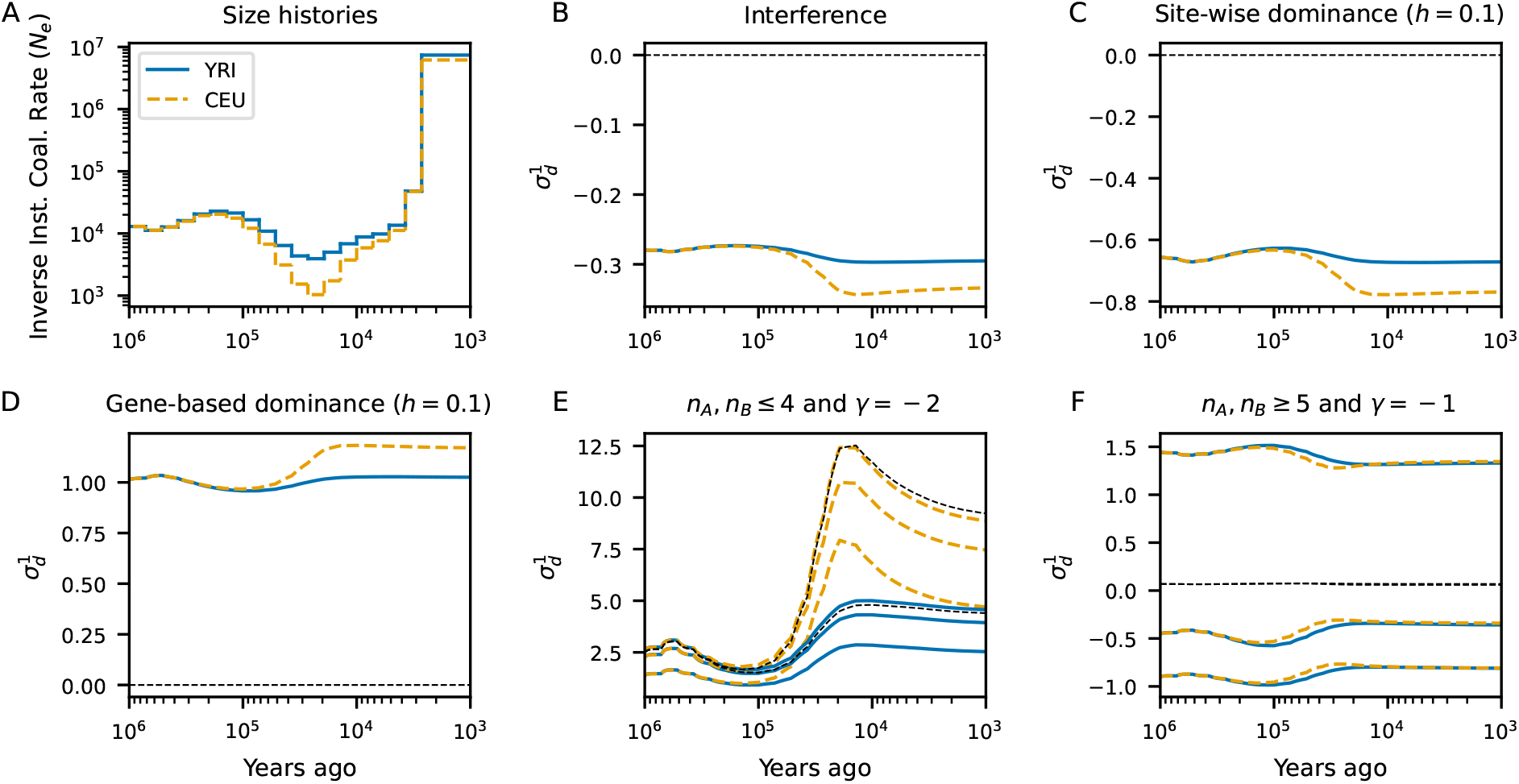
Signed LD under inferred models of human population-size history. Piecewise-constant population size histories inferred by `Relate` applied to 1000 Genomes Project Consortium *et al*. (2015) phase 3 data were used to simulate time series of two-locus statistics, as in Figure 6. (A) The CEU (Utah residents with Northern and Western European ancestry) are inferred to have a stronger bottleneck than the YRI (Yoruba in Ibadan, Nigeria) 10–100ka, reflecting the out-of-Africa event. Here, we highlight the effect of size changes on 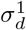 under non-additive selection models with *ρ* = 0 and γ = –2 at both loci, comparing standard interference (B) to site-wise dominance (C) and gene-based dominance (C). As with epistasis, more severe bottlenecks have larger effects on signed LD, and LD among common variants is more stable than among pairs of rare variants. Additional comparisons with epistasis and showing 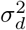 are in Figures S14–S16. Dashed lines indicate neutral expectations.

For each of the four histories, Figures 6, 7, and S11–S16 show the dynamics of 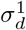 and 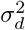) for a given parameterization of two-locus selection, including synergistic and antagonistic epistasis, dominance within loci, and gene-based dominance. In general across each selection model, population size expansions are not expected to strongly affect 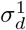, whether that expansion occurs deeper in the past as in the simple expansion model or rapid expansion more recently, as in the YRI. On the other hand, population size reductions tend to push signed LD to more extreme values and subsequent recoveries or expansion again reduce the magnitude of LD. Under no selection condition tested here do population size changes cause expected LD to change sign, showing that while the magnitude of deviation of LD from zero is sensitive to population size history, interpreting general patterns of the observed sign of LD in data should not be strongly affected by population size history.

### 2.3 Signed LD within protein-coding genes

Here, I examine patterns of signed LD between mutations in human protein-coding genes partitioned by functional annotations. Synonymous and missense mutations show similar levels of slighly positive signed LD when considering pairs of mutations within the same gene averaged over all autosomal chromosomes. Loss-of-function mutations have more negative LD, possibly due to differing modes of selective interactions for loss-of-function and missense mutations (Figure 8A–B). Within each population, measurement noise gives 95% confidence intervals that overlap with zero in each mutation class, although the observed patterns are remarkably consistent across African, European, and East Asian populations in the Thousand Genomes dataset. Comparing mean LD across populations, LD in Eurasian populations is somewhat larger on average, that is, more positive for missense mutations and more negative for loss-of-function mutations. This is in agreement with differences in expectations between populations that have or have not gone through a bottleneck in their recent history (Figures 6 and 7).

**Figure 8:**
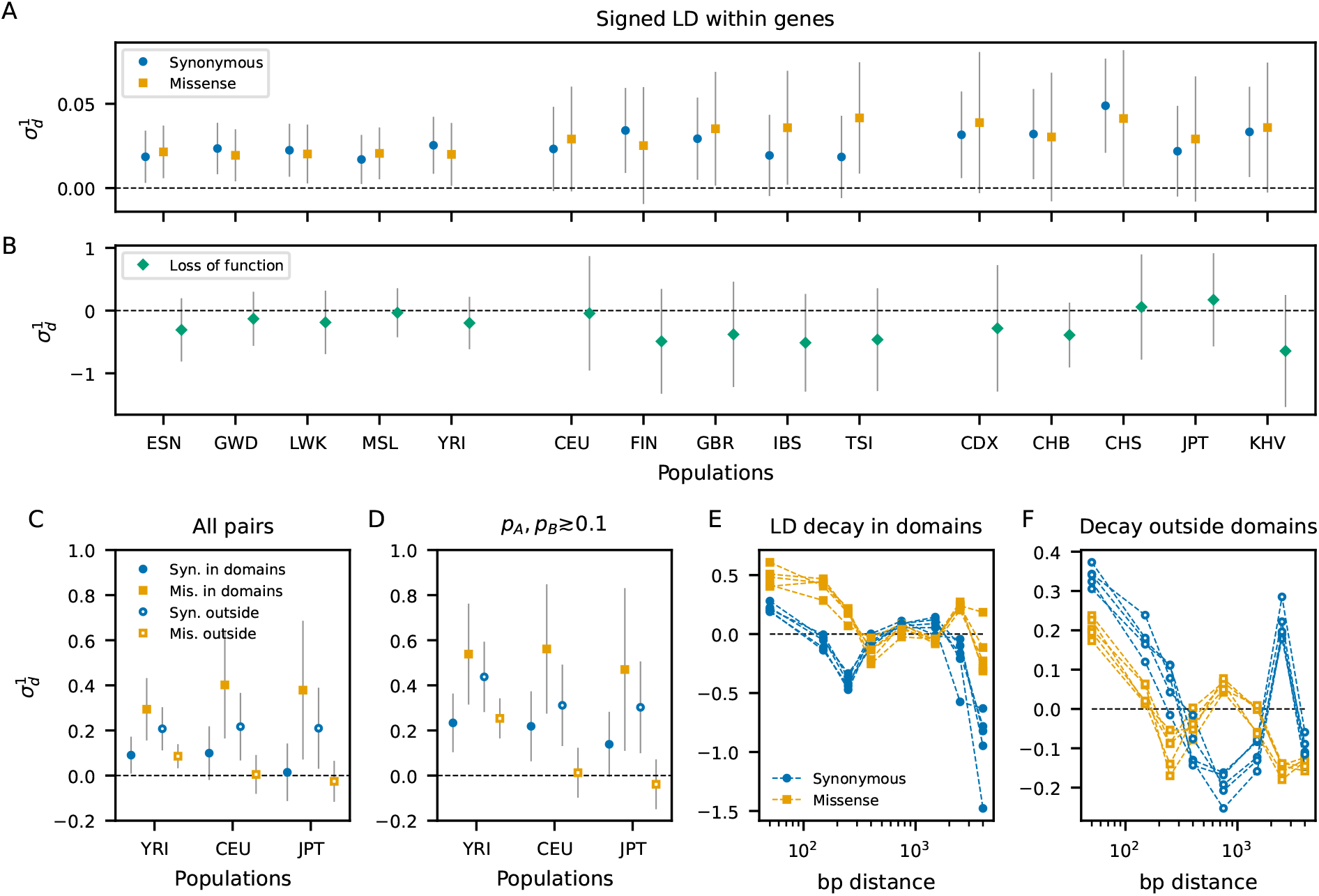
LD in human protein-coding genes and annotated domains. (A) Gene-wide averages of signed LD are slightly positive between pairs of both missense and synonymous mutations, considering pairs of mutations at matching distances. This positive, equal 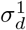 is also observed when conditioning on allele frequencies or considering only common variants with minor allele frequencies ≳ 0.1 (Tables S6–S11). (B) While there are relatively fewer pairs of loss-of-function mutations within genes, causing larger measurement uncertainty, such pairs tend to have negative average LD. Note that measurement noises for each class of mutations overlap with zero and with each other, making it difficult to draw firm conclusions on the patterns of interactions occurring gene-wide. (C) Partitioning pairs of mutations as falling within or outside of conserved domains reveals opposing patterns of signed LD, with 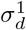 between missense mutations larger than that of synonymous mutations within domains. Outside of conserved domains, missense mutations have reduced LD compared to synonymous mutations. Distances of pairs outside of domains were matched to within-domain mutation pair distances. (D) This pattern again holds for common variants. (E, F) The signal of increased LD between missense variants within domains and decreased LD outside of domains is driven largely by tightly linked mutations (distances ≲ 400 bp), here showing African populations in the 1000 Genomes Project Consortium *et al*. (2015). Additional comparisons are shown in the Supporing Information, and Thousand Genomes population labels are described in Table S3.

#### 2.3.1 Positive LD between pairs of missense mutations in conserved domains

The similarity in signed LD between missense and synonymous mutations would suggest that interference between missense mutations is minimal, or at least no stronger than interference between synonymous mutations. However, interactive effects differ dramatically between pairs of mutations found in different intra-genic regions, with opposing effects canceling out when taking gene-wide averages. Due to the rarity of loss-of-function mutations, I only compare synonymous and missense mutations when looking at finer partitions of mutations within genes.

Annotated conserved domains in protein-coding genes play a significant role in driving signals of positive LD within genes. Such protein-coding domains are conserved elements of genes, often associated with some known functional or structural feature of a protein (Stanek *et al*., 2020). Purifying selection is expected to be stronger within conserved domains than within the same gene but outside of those domains. Indeed, the site-frequency spectrum (SFS) is skewed to lower frequencies for both missense and loss-of-function mutations within domains when compared to the same classes of mutations outside of domains, with much more negative values of Tajima’s D within domains (Table S4). On the other hand, no difference is observed for synonymous mutations whether within or outside domains, suggesting roughly equivalent effects of selection (either direct or linked) on synonymous variation.

Pairs of missense mutations that both fall within the same functional domain have large positive LD that is elevated above that of pairs of synonymous mutations that both fall within the same domain (Figures 8C–F and S18–S22). This difference between pairs of missense and synonymous variants within domains is especially pronounced for linked pairs within a few hundred base pairs of each other (Figures 8E and S18). Assuming most synonymous mutations are neutral, we would expect signed LD between missense mutations to be less than that between synonymous mutations under predictions from models of either Hill-Robertson interference, synergistic epistasis, and some models of dominance (Figures 3 and 4). This observation is opposite to those expectations, suggesting a different prevailing interactive effect between nonsynonymous mutations within domains.

The strength of selection on missense mutations within and outside of domains is observed to differ, leading to an excess of rare missense mutations within conserved domains (Table S4). LD is known to be sensitive to allele frequencies, with rare mutations showing large positive signed LD (Good, 2022). To test whether the signal of increased LD between missense mutations within domains is driven by rare variants, I considered subsets of pairs of mutations based on their derived allele frequences (Figures 8D and S20–S22). Rare and uncommon variants show large average LD for each class of mutations, but common variants recapitulate the opposing patterns of LD that is seen when averaging over pairs at all frequencies, as unconditioned statistics in the form of 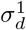 and 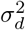 are dominated by common variants.

#### 2.3.2 Reduced LD between pairs of missense mutations outside of conserved domains

The large positive signal of LD for missense mutations within the same domain does not extend to pairs of missense mutations that are in different domains. Pairs of missense and synonymous mutations show nearly equal levels of LD close to zero across domains, with missense mutations slightly more negative than synonymous mutations (Figure S23). The interactive effect driving large LD in domains is therefore likely domain-specific. However, the average distance between pairs of mutations within domains is much smaller than between domains, and this observation may be primarily driven by the higher recombination distances between pairs of mutations across distinct domains.

Pairs of mutations that both fall outside of annotated domains have the opposite pattern of signed LD to pairs of mutations falling within the same domain. For pairs of mutations outside of domains but with distances matched to those within domains, pairs of synonymous mutations have larger positive LD than missense mutations. More distant pairs of mutations outside of domains, matched to the same distances as the between-domain comparison, each have LD roughly equal to zero (Figure S23).

The role that tightly linked variants have in driving these opposing signals can be seen in the decay of signed LD with distance between mutations (Figures S17–S19). Both synonymous and missense mutation pairs at distances greater than a few hundred base pairs have average LD that fluctuates around zero. However, for mutations outside domains, synonymous variants separated by short distances have large positive LD, while missense mutations have lower LD (Figure S19). In contrast, for mutations within the same domains, missense mutations have more positive LD at short distances than synonymous mutations (Figure S18).

## 3 Discussion

Previous theoretical and simulation studies have shown that interference and interactions between selected mutations reduce the efficacy of selection at linked loci, impacting substitution rates, the deleterious mutation load, and dynamics of segregating mutations (Hill and Robertson, 1968; Birky and Walsh, 1988; Barton, 1995; McVean and Charlesworth, 2000). Interference and synergistic epistasis between moderately deleterious mutations are both expected to cause negative LD between selected mutations, which can be readily tested using population genetic data (Sohail et al., 2017; Sandler et al., 2021; Garcia and Lohmueller, 2021). Here, I used a closed numerical approach to generate expectations for LD under a wide range of selective scenarios, and then compared patterns of LD in human populations between classes of coding mutations using unbiased estimators for LD from unphased genotypes.

By taking broad genome- and gene-wide surveys of LD across functional classes of mutations, heterogeneous patterns of interactive effects that occur within genes can be missed. From gene-wide averages, LD between pairs of missense mutations does not appear to be different from pairs of synonymous variants, although LD between loss-of-function variants is more negative. Hill-Robertson effects are expected to be strongest for slightly to moderately deleterious variants (with |*s*| ~ 1/*N*_e_), as strongly deleterious mutations are not expected to interfere with one another (McVean and Charlesworth, 2000). However, loss-of-function mutations do not typically fall within this regime, and inference of the distribution of fitness effects (DFE) for new loss-of-function variants shows that a large majority are strongly deleterious (Supporting Information).

Instead, negative synergistic epistasis between strongly deleterious mutations does produce large negative deviations of mean LD. Weakly deleterious recessive mutations can also produce this pattern, but strongly deleterious recessive mutations lead to slightly positive LD (Roze, 2021). While most loss-of-function mutations are strongly deleterious, those that rise to appreciable frequency are likely more benign and 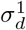 may be driven by patterns of weakly deleterious loss-of-function mutations. The difficulty in distinguishing these effects is compounded by the large measurement noise for 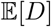, especially for loss-of-function variants for which only a few hundred within-gene pairs exist in the human population data analyzed here and which are separated by larger distances on average than neighboring missense and synonymous mutations.

In addition to LD varying by distance, LD can also vary due to differences in allele frequencies among classes of mutations (as selection drives mutations to higher or lower frequencies). Matching distances between pairs as well as allele frequecies between classes of mutations helps to reduce these concerns. It is possible that recombination rates can vary between annotated regions, resulting in differing patterns of background selection, which can affect allele frequencies and LD (e.g., Figures S7–S10). I did not condition on local recombination rates or inferred levels of background selection here, which is left to future work.

### 3.1 Non-uniform interactions between selected mutations within genes

Positive average LD between both missense and synonymous mutations has been reported in humans, *Drosophila*, and other species (Sohail *et al*., 2017; Sandler *et al*., 2021), while others have found that nonsynonymous mutations show lower LD than synonymous mutations (Garcia and Lohmüeller, 2021). The similarity of their gene-wide LD observed in this study (Figure 8) might suggest that interference or interactions between missense mutations are minimal, or at least no stronger than those between synonymous mutations. However, averaging over all observed pairs of mutations within a gene masks element-specific interactive effects that drive LD in opposite directions. Nonsynonymous mutations found within conserved protein domains, identified as conserved subunits of a gene with some structural or functional role (Stanek *et al*., 2020), are more strongly selected against on average, but also have increased signed LD over synonymous mutations at the same distances within domains. Missense mutation outside of domains but at the same distances as those within domains have more negative LD, both compared to distance-matched synonymous mutations outside of domains and to mutations within domains. Neither dominance effects (aside from very strongly selected recessive mutations (Roze, 2021)), synergistic epistasis, nor Hill-Robertson interference are expected to result in positive LD, so some other interactive effect should be driving this signal of positive LD within conserved domains.

There are a number of possible interaction scenarios that can result in positive signed LD between tightly linked loci. One explanation is a prevalence of pairs of compensatory mutations that are tolerated to cosegregate at high frequencies within conserved domains (Yeang and Haussler, 2007; Ivankov *et al*., 2014). Callahan *et al*. (2011) and Taverner *et al*. (2020) have proposed such a mechanism to explain observed clusters of nonsynonymous substitutions in *Drosophila* and other species. Another possibility is a model of antagonistic, or diminishing returns epistasis, in which a single amino acid-changing mutation within a domain damages the functionality of that subunit, but additional mutations within that same domain reduce fitness by a factor less than the first mutation. A third possibility, related to antagonistic epistasis, is that selection acts on the functional domain as a unit instead of on mutations within the domain individually (such as under a model of gene-based dominance). In this scenario, double heterozygotes have different fitnesses depending on whether the mutations are found on same haplotype or on different haplotypes.

We do not see an increase in LD between mutations that are found in different annotated domains within the same gene, which are on average at much larger recombination distances. Nor do we find increased LD outside of domains. Rather, for pairs of mutations outside of domains but matched to the same distance as those within the same domain, missense variants have considerably lower LD than synonymous variants. This difference between synonymous and missense pairs of variants largely disappears for SNPs separated by more than a few hundred base pairs (Figure 8E, F). This suggests that Hill-Robertson interference is the primary mode of interaction between missense mutations falling outside of domains, in agreement with Garcia and Lohmueller (2021), as epistasis is expected to impact LD over larger distances than what is observed. Importantly, however, the strength of epistasis is also likely to be a function of distance between mutations, complicating this interpretation.

Taken together, selection on segregating variants within protein-coding genes is nonuniform, with both the overall strength of selection and interactions between variants differing between annotated elements. Missense and loss-of-function mutations within conserved domains are subject to stronger selection, skewing the SFS to lower frequencies. Typical approaches for inferring the distribution of fitness effects from population genomic data average over these differences by considering all nonsynonymous mutations together (Boyko et al., 2008; Kim et al., 2017). If patterns of selection differ by both mutational class and location or if loss-of-function variants are in fact recessive, this can impact inferences of the DFE (Supporting Information). It would be straightforward to adapt DFE-inference methods to infer more detailed representation of the heterogeneous effects of new mutations within genes by partitioning by missense and loss-of-function classes as well as by annotated domains. Similarly, the different modes of interaction between mutations across annotated domains mean that our standard models and simulation approaches may be too simple to capture the evolutionary trajectories and patterns of diversity that differ at finer scales.

### 3.2 Causes of positive LD between synonymous mutations

In a non-structured randomly mating population, neutral mutations are expected to have average signed LD of zero, but across all populations analyzed here, LD between synonymous mutations is positive. While selection on some subset of synonymous variants is possible, it should be much weaker on average than between missense mutations, and any interference between selected synonymous variants should lead to negative LD.

Spatial population structure may be responsible for the increase of LD observed between synonymous variants. Neutral evolution in structured populations alone cannot create this effect. Under standard population genetic assumptions of constant mutation and recombination rates and independent mutation events, migration, admixture, and spatial structure do not result in nonzero 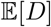. Rather, for a set of mutations with *given* differences in allele frequencies between source populations, *D* can be nonzero in an admixed population between those particular mutations even when *D* is zero in the source populations (Cavalli-Sforza and Bodmer, 1971). This result relies on conditioning on directional differences in allele frequencies between populations. Without conditioning, the expected difference in mean allele frequency between source populations is zero 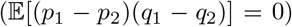, as differences are equally likely to be positive or negative, and thus resulting 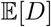 is still zero when taking genome-wide averages.

Instead, Sohail *et al*. (2017) and Sandler *et al*. (2021) used forward simulations under multi-population or explicit spatial models and a multiplicative fitness function to explore the joint effects of structure, assortative mating, and interference, finding that together these can lead to nonzero LD. For example, Sandler *et al*. (2021) found positive LD among neutral mutations in a simulation with admixture and linked selected mutations, showing that the combined effects of spatial structure and background selection can lead to positive average LD between neutral mutations. Positive LD between synonymous mutations is only observed at short distances in human data, which could be used to constrain such models that predict positive LD over varying distances.

Nonrandom mutational processes provide an alternative explanation for positive LD between neutral variants separated by short distances, in which clusters of mutations occur simultaneously in the same mutational event. Such multinucleotide mutations have been shown to affect patterns of LD over short distances, on the order of 10s to 100s of base pairs (Harris and Nielsen, 2014), and clustered mutational events may be common in humans (Besenbacher et al., 2016). Indeed, a simple exponential model in which the fraction of mutations causing a multinucleotide mutation event decays with distance fits the observed patterns of 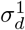 between synonymous variants (Supporting Information, Figure S28), with the best-fit model requiring only a small fraction of mutations to cause multinucleotide mutation events. We may therefore treat positive LD as the baseline expectation for tightly linked variants primarily due to clustered mutations, so that subsequent selective processes and interactions cause LD to deviate from that expectation, as seen for missense mutations across different sub-elements of genes. It may also be more appropriate to compare the negative LD observed between linked loss-of-function variants to that positive baseline expectation instead of zero, which would imply that they are more recessive or have stronger synergistic epistasis than from inferences assuming a neutral expectation of zero. Additional analyses and extensive simulations will be required to tease apart the effects of population structure, selective interference at linked sites, and clustered mutational events on patterns of LD.

### 3.3 Challenges to distinguishing highly parameterized models of selective interactions

When partitioning measurements of LD by mutation classes or regions within genes, the decreasing number of pairwise comparisons leads to large estimated measurement noise. Within each population, confidence intervals of observed 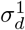 often overlap with zero or overlap with that of other classes of mutations. While observed patterns are remarkably consistent across the 15 populations considered here, their joint evolutionary histories make formal testing of significance difficult due to shared variation, as they cannot be treated as independent measurements. Extensive forward simulations will likely be needed to more thoroughly assess significance.

From a modeling perspective, in exploring or inferring multi-locus selective models, the number of plausible selection scenarios becomes quite large as we relax the strict assumptions of additivity and multiplicative interactions. The inclusion of dominance effects, epistasis, or other interactions leads to a rapid increase in the number of parameters to consider. This makes performing forward simulations that span the range of all such selective interaction scenarios burdensome.

Instead, closed, numerical approaches to compute expectations for two-locus statistics under a wide range of selection models, as presented here and in Friedlander and Steinrücken (2022), allow us to explore this highly parameterized space of models far more efficiently, and it opens the possibility for performing likelihood-based inference using signed LD or other two-locus summaries of the data. For example, inferring the joint distribution of dominance and selection is underpowered using the SFS alone, but because signed LD is sensitive to the levels of dominance (Figure 4), inferring the DFE with dominance may be feasible using the joint distribution of allele frequencies and LD. The results presented in this paper likely do not cover the space of all possible two-locus models, and other unexplored models may result in similar patterns of signed LD. Comparisons to empirical data should therefore be treated with caution, and additional careful modeling will allow a more firm determination of the interactive effects causing observed patterns of variation.

Using a single low-order summary of signed LD, such as 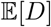 or 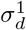, is likely insufficient to confidently discriminate modes of selective interactions between linked mutations. Both dominance and epistasis can result in strongly negative LD, although the expected decay of LD under those scenarios can differ. Similarly, multiple interaction models can lead to positive signed LD. Among all interaction models, the extent of LD and rate of its decay also depends on the underlying distribution of selection coefficients among a class of mutations, which are unknown for a given pair of mutations, so that we must integrate over a distribution of fitness effects. This DFE, however, will have been inferred under a simple set of assumptions, such as additivity and interchangeability between sites within a gene, potentially biasing any inference using previously inferred DFEs to learn about patterns of interactions. Again, this underlines the need to jointly infer strengths and interactions of selected variants and to consider patterns of variation at finer genomic scales.

Finally, exploring additional summaries of the two-locus sampling distribution should increase power to distinguish between interaction models. The numerical methods developed here provide the full two-locus sampling distribution, and expectations for a large family of informative two-locus statistics can be computed directly from this distribution. These expectations can then be compared directly to empirical observations, which can be taken from either phased or unphased data (Ragsdale and Gravel, 2020). Additional work performing such an analysis is therefore warranted, which should provide a path forward for distinguishing between modes of selective interactions.

## 4 Methods

### 4.1 Existing theory and numerical methods

Many well-known properties of two-locus dynamics and equilibrium LD come from early work on the multilocus diffusion approximation (Kimura, 1955; Hill and Robertson, 1968; Ohta and Kimura, 1969, 1971). This includes the result that genome-wide averages of signed LD are expected to be zero under neutrality. Under a two-locus biallelic model, where the left locus allows alleles *A* and *a* and the right locus allows alleles *B* and *b*, the standard covariance measure of LD is defined as *D* = *f_AB_* – *f_A_f_B_*, where *f_AB_* is the haplotype frequency of double-derived types carrying both *A* and *B*, and *f_A_* and *f_B_* are the marginal frequencies of the derived alleles at each locus. This covariance decays due to both drift and recombination at a rate proportional to the inverse of the effective population size and the distance separating loci (Hilland Robertson, 1968):

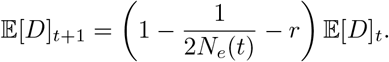

While 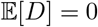, the variance of *D* is non-zero, and Ohta and Kimura (1971) presented their groundbreaking result that the variance of *D* under neutrality and steady-state demography, normalized by the joint heterozygosity of the two loci, is

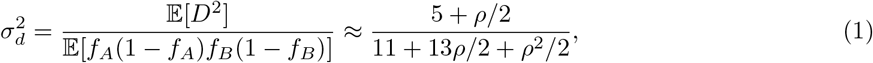

where *α* = 4*N*_e_*r*.

Analytic progress beyond these results has come haltingly. In the 1980s, recursions were developed to compute the two-locus sampling distribution under neutrality, that is, the probability of observing given counts of two-locus haplotypes in a sample of size n (Golding, 1984). This approach would continue to be developed and later form the foundation for the inference of local recombination rates from population genetic data (Hudson, 2001; McVean *et al*., 2004). More recently, Song and Song (2007) computed 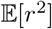 using a diffusion approximation approach, although their solution involves the summation of infinitely many terms and is restricted to neutrality and steady-state demography.

To include selection, however, there have been relatively few advances beyond the Monte Carlo simulation approach taken by Hill and Robertson (1966), albeit now with more powerful computational resources and sophisticated software for performing flexible forward simulation (e.g., Thornton, 2019; Haller and Messer, 2019). Analytic results for two-locus distributions under selection are notoriously difficult, with a few notable flashes of progress. For example, McVean (2007) considered the effect of a recent sweep on patterns of LD between neutral loci near the locus under selection, and in a recent paper, Good (2022) presented analytic solutions for patterns of LD between rare mutations under additive selection with epistasis. Nonetheless, such approaches are typically confined to steady-state demography and constrained selection models.

Numerical methods inhabit the space in between expensive discrete simulations and limited analytic solutions, providing a more efficient and practical method to compute expectations of two-locus diversity measures under a wider range of parameters and demographic scenarios. Ragsdale and Gutenkunst (2017) used a finite differencing approach to numerically solve the two-locus diffusion equation with additive selection at either locus, although they focused on the applicability of two-locus statistics to demographic inference. More recently, Ragsdale and Gravel (2019) extended the Hill and Robertson (1968) system for 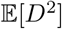 to compute arbitrary moments of the distribution of *D* for any number of populations connected by migration and admixture. They also showed that such a moments-construction can be used to solve for the two-locus sampling distribution within a single population, though it requires a moment-closure approximation for nonzero recombination and selection. Below, I extend this approach to model arbitrary diploid selection, which encompasses dominance, epistasis, and other forms of selective interactions between two loci. In a concurrent study to this paper, Friedlander and Steinrücken (2022) developed numerical solutions to the same moments system, which they used to describe selected haplotype trajectories and the distortion of neutral diversity at loci variably linked to beneficial alleles that sweep to high frequencies under non-equilibrium demography.

### 4.2 The two-locus sampling distribution with arbitrary selection

The two-locus sampling distribution is the direct analog to the single-locus site frequency spectrum (SFS) of a given sample size (Figure 1). Instead of describing the density or number of mutations with a given allele count out of *n* samples, the two-locus distribution ψ_*n*_ stores the density or number of pairs of loci with observed haplotype counts, so that ψ_*n*_(*i*, *j*, *k*) is the number of pairs for which we observe *i* copies of the *AB* haplotype, *j* of type *Ab*, *k* of type *aB*, and *n* – *i* – *j* – *k* of type *ab*. The size of ψ_*n*_ grows rapidly, with *O*(*n*^3^) entries, which practically limits computational approaches to moderate sample sizes *n* ≲ 100 and a single population.

Under neutrality, a number of approaches exist to compute ψ_*n*_, including the recursion due to Golding(1984) and Ethier and Griffiths (1990), or more recent numerical approaches to the two-locus coalescent (Kamm *et al*., 2016) or diffusion approximation (Ragsdale and Gutenkunst, 2017). Selection is most easily included using the forward-in-time diffusion equation (Kimura, 1955; Hill and Robertson, 1966), where a standard approach is to first solve for the continuous distribution *ψ* of the density of two-locus haplotype configurations in the full population, and then integrate *ψ* against the multinomial sampling function to obtain ψ_*n*_.

Alternatively, Ragsdale and Gravel (2019) showed that there exists a system of ordinary differential equations directly on the entries of ψ_*n*_. I briefly summarize this general approach below, but refer readers to that paper for detailed derivations of the drift, recombination, and mutation terms and the moment-closure approximation. Instead, here I focus on generalizing the selection operator to include epistasis, dominance, and other forms of two-locus interactions.

#### 4.2.1 Moment equation for ψ_*n*_

The system of linear ordinary differential equations for the entries of ψ_*n*_ takes the form

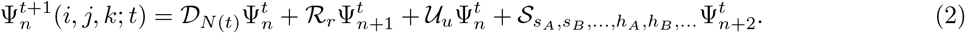

Here, 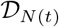 is a sparse linear operator accounting for drift with population size *N*(*t*), 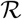 accounts for re-combination with per-generation recombination probability *r* between the two loci, 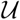 accounts for mutation, either under an infinite sites or biallelic reversible mutation model, and S accounts for selection.

The moment system for ψ_*n*_ can be derived directly from the diffusion approximation, or it can be found through a more intuitive process of tracking the dynamics of allelic states of a sample of size of *n* from the full population. We assume *n* ≪ *N*_e_, and *r* and *s* are *O*(1/*N*_e_) so that multiple coalescence, recombination, or selective events within the *n* lineages are rare in any given generation (Supporting Information; Jouganous *et al*., 2017; Ragsdale and Gravel, 2019). In typical diffusion approximation fashion, we multiple through by *2N_ref_* so that time is measured in *2N_ref_* generations, and we consider scaled parameters *ρ* = 4*Nr*, *θ* = 4*Nu*, and γ = 2*Ns*.

#### 4.2.2 Moment closure

In the absence of selection and for fully linked loci (i.e., *ρ* = 0), the system is closed and can be solved exactly. However, for nonzero recombination or selection, the entries of ψ_*n*_ rely on the slightly larger sampling distributions with sample sizes *n* +1 (for recombination and additive selection) or *n* + 2 (for nonadditive selection). This is because if a recombination event occurs within one of n lineages being tracked by ψ_*n*_, we need to draw an additional lineage from the full population to recombine with that chosen lineage, thus requiring 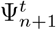 to find 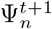. Selection events similarly require extra lineages from the full population, which replace a chosen lineage that fails to reproduce with probability proportional to its relative fitness.

This requirement of extra lineages for nonzero recombination and selection means that the system in (2) is not closed, so that we need a moment-closure approximation to solve for ψ_*n*_. As in Ragsdale and Gravel (2019), a jackknife approximation is used to estimate ψ_*n*+*l*_, for *l* = 1 or 2, from ψ_*n*_ (following the single-locus closure introduced in Jouganous *et al*., 2017), so that 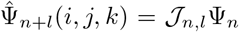, although other accurate closure approximations are possible (Friedlander and Steinrucken, 2022). This emits a closed approximate system,

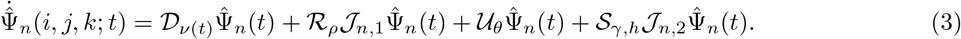

The jackknife approximation, which approximates an entry ψ_*n*+*l*_(*i*, *j*, *k*) using nearby entries in ψ_*n*_, is more accurate for larger sample sizes, creating a tension between efficiency and accuracy: larger sample sizes result in more accurate solutions, as error in the jackknife is diminished, but computational complexity also grows rapidly in the number of entries of ψ_*n*_, which is *O*(*n*^3^) (Figure S1). In the results presented in this paper, sample sizes between *n* = 30 and *n* = 80 are used. Derivations for the drift, recombination, and mutation operators and the jackknife moment-closure approximation can be found in section S1.3 of Ragsdale and Gravel (2019), and I repeat the main results in the Supporting Information of this paper.

#### 4.2.3 Selection models with epistasis and dominance

To include selection, we consider a model where we draw lineages uniformly from the previous generation, but keep lineages with probability proportional to their fitness. In the absence of dominance, selection reduces to a haploid model, with acceptance and rejection probabilities depending on the fitnesses of each haploid copy, where haplotype *Ab* has fitness 1 + *s_A_*, *aB* has fitness 1 + *s_B_*, and *AB* has fitness 1 + *s_AB_*. We assume the doubly ancestral haplotype *ab* has fitness 1, so fitnesses are relative to that of *ab* haplotypes. The standard multiplicative fitness function assumes that *s_AB_* ≈ *s_A_* + *s_B_* (assuming *s*^2^ ≈ 0), and a model for epistasis can be written as

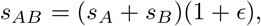

so that *ϵ* > 0 implies synergistic epistasis, while *ϵ* < 0 implies antagonistic epistasis.

To obtain the recursion equation under selection we consider drawing *n* lineages from generation *t*, which has an expected sampling distribution of haplotype counts given by 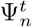. However, assuming *s* ≤ 0 for each derived haplotype, each of those sampled lineages has probability of being rejected equal to the absolute value of the selection coefficient assigned to its haplotype state. If a lineage is rejected, a replacement is drawn from the full population. Under the assumption that *ns* ≪ 1, the probability that more than one selection event occurs in any given generation is negligibly small, so that the case of multiple simultaneous rejections can be ignored. Then 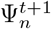 relies only on 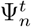 and 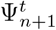 for additive selection. The full selection operator S for additive selection is given in the Supporting Information.

To account for dominance, or other general forms of two-locus selection, the selection operator no longer reduces to individual haplotypes, but instead we need to know the state of two-locus genotypes. For example, the fitness of an individual carrying an *Ab* haplotype depends on whether their second haplotype is *ab*, *Ab*, *aB*, or *AB*. We can therefore assign a selection coefficient to each possible diploid configuration, *s_Ab/ab_*, *s_Ab/Ab_*, and so on. Assuming that the doubly homozygous ancestral *ab/ab* genotype has relative fitness 1, this gives nine possible unique selection coefficients in the most general two-locus selection model. Note that *AB*/*ab* and *Ab*/*aB* genotypes need not have the same selection coefficient, which allows for simulation under a gene-based model of dominance (Table S1).

The general selection operator follows the same approach as the haploid selection operator with epistasis described above. Now, in the case of a selection event rejecting a lineage within our tracked samples, we need to draw not only the replacement lineage from the full population but also a second haplotype from the full population to form the diploid genotype, as this determines the probability that we reject the focal haplotype. We again assume that *ns* ≪ 1 for all genotype selection coefficients, so that we may assume at most a single selection event occurs in any given generation. This means that to find 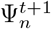 under a general two-locus selection model, which encompasses dominance within either locus, gene-based dominance, or a combination of dominance and epistatic effects, we need 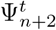. Again, a full derivation and expressions for the general selection operator are given in the Supporting Information.

#### 4.2.4 Low-order summaries of the sampling distribution

From ψ_*n*_, expectations for any two-locus statistic can be found by downsampling to the appropriate sample size. For example, to compute 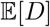, the sum is taken over all haplotype configurations **n** = (*n_AB_*, *n_Ab_*, *n_aB_*), weighted by the density ψ_*n*_ for that configuration:

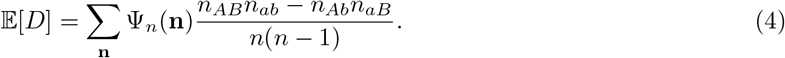

For large sample sizes, this is approximately equal to computing *D* by taking the maximum likelihood estimate for each allele frequency *f_i_* = *n_i_*/*n*, but the maximum likelihood-based estimate will be noticeably biased for small to moderate sample sizes. Other low-order two-locus statistics can be computed using the same approach, as implemented in moments following Ragsdale and Gravel (2020), which can be compared across sample sizes and between estimates from phased or unphased data. In this paper, I focus on 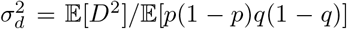 and 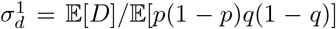, which can be averaged over pairs of variants at all frequencies. Allele-frequency conditioned statistics (such as keeping only loci below some frequency threshold as in Good (2022)) can be considered using this same approach.

#### 4.2.5 Simulations of non-steady-state demography

I considered four variable population size histories, two simple toy models and two inferred from human populations in African and Europe using `Relate` (Speidel *et al*., 2019). For each size history scenario, I tracked the evolution of ψ_*n*_(*t*) for varying selection models, plotting the trajectories of 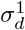 and 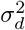 over time (Figures 6, 7, and S11–S16). The selection strength at both loci was fixed at either γ = –1 or –10 for the models with epistasis, or γ = –2 for models with dominance, and recombination was set to zero.

The simple size change models both had ancestral *N_e_* = 10,000, with one a 3-fold population expansion that occurs 3,000 generations ago, and the other a 5-fold reduction 2,000 generations ago followed by a recovery to its initial size 1, 000 generations ago. The size histories for YRI and CEU were inferred using `Relate` (Speidel *et al*., 2019) applied to the phase 3 haplotype-phased autosomal data from 1000 Genomes Project Consortium *et al*. (2015), using default parameters as recommended in the `Relate` online tutorial, assuming a mutation rate of 1.25 ×10^-8^per-bp per-meiosis and a human generation time of 29 years. `Relate` returns estimates of coalescence rates within specified time bins, and population sizes are estimated as their inverses. Estimates using `Relate` for population sizes in the very recent past (< 3,000 years, or ≈ 100 generations) diverged, so I truncated the history over this time period and assumed a constant size from the most recent non-diverged bin.

### 4.3 Analysis of human genomic data

Using the annotated variant call format (VCF) files from the phase 3 1000 Genomes Project Consortium *et al*. (2015) (Thousand Genomes) data release, I subset the genotype VCFs to autosomal variants that were annotated as either synonymous or nonsynonymous, including both missense mutations and more damaging “high impact” loss-of-function mutations. Loss-of-function annotations include frameshifts, splice acceptor, splice donor, start loss, stop gain, stop loss, and transcript ablation variants. I further subset to samples within each non-admixed population in the African, European, and East Asian continental groups (five populations each, Table S3). Signed LD is sensitive to ancestral state misidentification, so I only kept sites for which ancestral alleles were estimated with high confidence in both the VCF info field and the Thousand Genomes human ancestor reconstructed from a phylogeny of six primates.

In addition to ancestral state misidentification, measured LD is sensitive to phasing error, so I computed LD statistics using unphased genotypes following Ragsdale and Gravel (2020). This approach provides unbiased estimates for pairwise LD, under the assumption that individuals are not inbred. I considered pairs of mutations within the same mutation class (synonymous, missense, and loss-of-function) either within the same gene and inside or outside of annotated domains within the same protein-coding genes. I used a dataset of annotated protein domains mapped to the hg19 human reference build compiled by Stanek *et al*. (2020) to determine if a given mutation falls within a domain or not.

### 4.4 Data and software availability

All data and software used in this paper are publicly available and open source. I downloaded the Thousand Genomes annotations and genotypes VCFs from the ftp server at ftp://ftp.1000genomes.ebi.ac.uk/vol1/ftp/release/20130502/, and the Thousand Genomes human ancestor fasta file from ftp://ftp.1000genomes.ebi.ac.uk/vol1/ftp/phase1/analysis_results/supporting/ancestral_alignments/. Protein domain information from Stanek *et al*. (2020) was downloaded from http://prot2hg.com/dbdownload.php.

Implementation of moment equations to compute expectations for two-locus and linkage disequilibrium statistics are implemented in Python using Numpy (Harris et al., 2020) and sparse matrix solvers in Scipy (Virtanen et al., 2020). These methods are packaged within `moments`, and analyses here were performed using moments version 1.1.10, available from https://bitbucket.org/simongravel/moments/src/master/ and via `conda`, with extensive documentation at https://moments.readthedocs.org. Scripts to run all analyses, recreate figures, and compile this manuscript are available at https://github.com/apragsdale/two_locus_selection.

## 5 Acknowledgements

I thank Alex Diaz-Papkovich, Eric Friedlander, Simon Gravel, Mashaal Sohail, Matthias Steinrücken, and Kevin Thornton for helpful discussions and feedback on earlier versions of this manuscript.

