## Supporting Information for "Local fitness and epistatic effects lead to distinct patterns of linkage disequilibrium in protein-coding genes"

Aaron P. Ragsdale

Department of Integrative Biology, University of Wisconsin–Madison, WI, USA

April 1, 2022

**S3 Supporting references** 8

**S4 Supporting tables** 10

**S5 Supporting figures** 20

### S1 The diffusion equation and moment system for the two-locus sampling distribution

The two-locus diffusion equation with additive selection was first described by Kimura (1955) and studied extensively in the 1960s and 70s (e.g., Hill and Robertson (1966); Ohta and Kimura (1969)). The continuous distribution $\psi(x_1, x_2, x_3)$ of haplotype frequencies in a population, where $x_1$ is the frequency of $AB$, $x_2$ of $Ab$, and $x_3$ of $aB$, is governed by the multi-dimensional Fokker-Planck equation:

$$\begin{aligned}\frac{\partial \psi}{\partial \tau} =& \frac{1}{2}\sum\sum_{1\leq i,j\leq 3} \frac{\partial^2}{\partial x_i \partial x_j}\left[\frac{x_i(\delta_{i=j} - x_j)\psi}{\nu(\tau)}\right] \\ &- \frac{\rho}{2}\left(-\frac{\partial}{\partial x_1}D\psi + \frac{\partial}{\partial x_2}D\psi + \frac{\partial}{\partial x_3}D\psi\right) \\ &- \frac{\gamma_A}{2}\left[\frac{\partial}{\partial x_1}x_1(1-x_1-x_2)\psi + \frac{\partial}{\partial x_2}x_2(1-x_1-x_2)\psi - \frac{\partial}{\partial x_3}x_3(x_1+x_2)\psi\right] \\ &- \frac{\gamma_B}{2}\left[\frac{\partial}{\partial x_1}x_1(1-x_1-x_3)\psi - \frac{\partial}{\partial x_2}x_2(x_1+x_3)\psi + \frac{\partial}{\partial x_3}x_3(1-x_1-x_3)\psi\right].\end{aligned} \tag{S1}$$

$D$ is the standard covariance measure of linkage disequilibrium,

$$D = x_1 - (x_1 + x_2)(x_1 + x_3) = x_1x_4 - x_2x_3,$$

$\gamma_A$ and $\gamma_B$ are the (additive) scaled selection coefficients at the left and right locus, and $\rho$ is the scaled recombination rate between the two loci. Time $\tau$ is measured in $2N_e$ generations, and $\nu(\tau)$ is the population size relative to the ancestral size or some reference size at time $\tau$.

Given a function $\psi$ that solves Equation S1, the two-locus sampling distribution for a sample size of $n$ haploids can be found by integrating $\Psi$ against the multinomial sampling function, so that

$$\Psi_n(i,j,k) = \binom{n}{i,j,k,n-i-j-k} \iiint_{\substack{x_1,x_2,x_3\geq 0\\ x_1+x_2+x_3\leq 1}} \psi(x_1,x_2,x_3)x_1^i x_2^j x_3^k(1-x_1-x_2-x_3)^{n-i-j-k}dx_1dx_2dx_3. \tag{S2}$$

In the method-of-moments approach, instead of solving the differential equation for $\psi$, we instead integrate both sides of the differential equation against the multinomial sampling function for a given sampling configuration $(i,j,k)$. On the left side, we get $\partial_t \Psi_n(i,j,k)$, and on the right we obtain, after some simple integration by parts and somewhat tedious simplification, terms for drift, recombination, and selection that can be written as sparse linear operators of $\Psi_n$. Written compactly, this takes the form

$$\partial_\tau \Psi_n = \frac{1}{2\nu(\tau)}\mathcal{D}_n\Psi_n + \frac{\rho}{2}\mathcal{R}_n\Psi_n + \frac{\theta}{2}\mathcal{U}_n\Psi_n + \mathcal{S}_{n,\boldsymbol{\gamma},\mathbf{h}}\Psi_n. \tag{S3}$$

.

Alternatively, we arrive at this same linear system of equations by tracking the expected sampling distribution over $n$ lineages within the full population and how that changes over time by drawing lineages from one generation to the next in the style of Wright (1931). Both Jouganous *et al.* (2017), for the single-locus SFS, and Ragsdale and Gravel (2019) drew this connection in detail, so I refer readers to those previous papers for a fuller description of those derivations and discussion. In the next section I simply repeat the results for $\mathcal{D}$, $\mathcal{R}$, and $\mathcal{U}$, briefly describe the moment closure approximation (which is the same as presented in Ragsdale and Gravel (2019)), and then describe the selection operator $\mathcal{S}$ for selection with epistasis, both with and without dominance.

#### S1.1 Drift, mutation, recombination, and moment closure

##### S1.1.1 Drift

Drift for an entry $(i,j,k)$ depends only on $\Psi_n$ and therefore closes. The entries of $\mathcal{D}$ are found by considering the possibility of a coalescence event occurring within a given generation within $n$ lineages in the full population. If $n \ll N$, we can safely assume that at most a single such event occurs in any given generation.

$$
\begin{aligned}
\mathcal{D}_n(i,j,k)\Psi_n =&(i-1)(n-i-j-k+1)\Psi_n(i-1,j,k) \\
&+(i+1)(n-i-j-k-1)\Psi_n(i+1,j,k) \\
&+(i-1)(k+1)\Psi_n(i-1,j,k+1) \\
&+(i+1)(k-1)\Psi_n(i+1,j,k-1) \\
&+(i-1)(j+1)\Psi_n(i-1,j+1,k) \\
&+(i+1)(j-1)\Psi_n(i+1,j-1,k) \\
&+(j-1)(n-i-j-k+1)\Psi_n(i,j-1,k) \\
&+(j+1)(n-i-j-k-1)\Psi_n(i,j+1,k) \\
&+(j-1)(k+1)\Psi_n(i,j-1,k+1) \\
&+(j+1)(k-1)\Psi_n(i,j+1,k-1) \\
&+(k-1)(n-i-j-k+1)\Psi_n(i,j,k-1) \\
&+(k+1)(n-i-j-k-1)\Psi_n(i,j,k+1) \\
&-2\left(i(n-i-j-k)+ik+ij+j(n-i-j-k)+jk+k(n-i-j-k)\right)\Psi_n(i,j,k)
\end{aligned} \tag{S4}
$$

###### S1.1.2 Recombination

If a lineage in our sample of size $n$ recombines in a given generation, which occurs with probability $nr$, we need to draw an extra lineage from the full population for it to recombine with. This means we need $\Psi_{n+1}$ in the previous generation. After drawing that extra lineage, $\Psi_n$ changes as we draw one of the two recombinant types (each with probability 1/2) instead of the lineage that was chosen to recombine.

$$
\begin{aligned}
\mathcal{R}_n(i,j,k)\Psi_n =&\frac{(i+1)(n-i-j-k+1)}{n+1}\Psi_{n+1}(i+1,j-1,k) \\
&+\frac{(i+1)(n-i-j-k+1)}{n+1}\Psi_{n+1}(i+1,j,k-1) \\
&+\frac{(j+1)(k+1}{n+1)}\Psi_{n+1}(i-1,j+1,k+1) \\
&+\frac{(j+1)(k+1}{n+1)}\Psi_{n+1}(i,j+1,k+1) \\
&-\frac{(i+1)(n-i-j-k)}{n+1}\Psi_{n+1}(i+1,j,k) \\
&-\frac{(j+1)k}{n+1}\Psi_{n+1}(i,j+1,k) \\
&-\frac{j(k+1)}{n+1}\Psi_{n+1}(i,j,k+1) \\
&-\frac{i(n-i-j-k+1}{n+1}\Psi_{n+1}(i,j,k)
\end{aligned} \tag{S5}
$$

###### S1.1.3 Mutation

We assume an infinite sites mutation (ISM) model where new mutations occur at previously unmutated loci. In the two-locus ISM model, two-locus pairs of variable loci arise when a mutation occurs at one locus when the other locus is already variable. Thus, new mutations at the $B/b$ locus occur against the single-locus allele frequency distribution $\Phi_{n,A}$, and new mutations at the $A/a$ locus occur against $\Phi_{n,B}$, which are found via the single-locus system from Jouganous *et al.* (2017).

$$
\begin{aligned}
\mathcal{U}_n(i,j,k)\Psi_n =&(j+1)\frac{\theta_B}{2}\Phi_{n,A}(j+1)\delta_{i=1,k=0} \\
&+(n-j)\frac{\theta_B}{2}\Phi_{n,A}(j)\delta_{i=0,k=1} \\
&+(i+1)\frac{\theta_A}{2}\Phi_{n,B}(i+1)\delta_{i=1,j=0} \\
&+(n-i)\frac{\theta_A}{2}\Phi_{n,B}(i)\delta_{i=0,j=1}
\end{aligned}
\tag{S6}
$$

###### S1.1.4 Jackknife moment closure approximation

We use a jackknife approximation to write the entries of $\Psi_{n+1}$ and $\Psi_{n+2}$ as linear combinations of entries in $\Psi_n$. The general strategy is to assume the underlying continuous distribution $\psi(x_1, x_2, x_3)$ can be approximated locally as a quadratic, and then use entries in $\Psi_n$ that are close in frequency to a given entry in $\Psi_{n+l}$ to estimate the coefficients of that quadratic using the multinomial sampling formula. Then this quadratically local approximation to $\psi$ can be used to compute $\Psi_{n+l}(i,j,k)$ using Eq. (S2). Readers should refer to section S1.3.5 in the Supporting material for Ragsdale and Gravel (2019) for details.

#### S1.2 Selection

First consider the case of no dominance, so that the haplotypes $Ab$, $aB$, and $AB$ have selection coefficients $s_{Ab}$, $s_{aB}$, and $s_{AB}$, respectively. Note that the case with $s_{AB} = s_{Ab} + s_{aB}$ implies no epistasis between the $A/a$ and $B/b$ loci. Here, we assume all selection coefficients are negative. In a given generation, a selection event could occur in which a haplotype is rejected (selected against) with probability proportional to its selection coefficient, $-s$. We then draw an extra lineage from the full population to replace that rejected lineage.

For example, the probability that an $AB$ haplotype is selected against and replaced by an $Ab$ haplotype is

$$
-ns_{AB}\frac{i}{n+1}j+1n\Psi_{n+1}(i,j+1,k),
$$

where the additional $j+1$ lineage in a sample of size $n+1$ accounts drawing that extra $Ab$ haplotype. Taking all such selective events together, for additive selection we get

$$
\begin{aligned}
\mathcal{S}_n(i,j,k)\Psi_n =&\frac{i+1}{n+1}\left(-s_{AB}(n-i)+s_{Ab}j+s_{aB}k\right)\Psi_{n+1}(i+1,j,k) \\
&+\frac{j+1}{n+1}\left(s_{AB}i-s_{Ab}(n-j)+s_{aB}k\right)\Psi_{n+1}(i,j+1,k) \\
&+\frac{k+1}{n+1}\left(s_{AB}i+s_{Ab}j-s_{aB}(n-k)\right)\Psi_{n+1}(i,j,k+1) \\
&+\frac{n-i-j-k+1}{n+1}\left(s_{AB}i+s_{Ab}j+s_{aB}k\right)\Psi_{n+1}(i,j,k)
\end{aligned}
\tag{S7}
$$

For a general diploid selection model, the idea is nearly the same, but we need to draw an extra lineage to determine the fitness of a diploid individual. For example, the probability that an $AB$ haplotype is paired with an additional lineage $Ab$ and selected against, and then replaced by an $aB$ haplotype is

$$
-ns_{AB/Ab}\frac{i}{n+2}\frac{j+1}{n+1}\frac{k+1}{n}\Psi_{n+2}(i,j+1,k+1).
$$

There are now many more possible selective events to consider, but after accounting for all possible diploid pairs and replacements (90 in total) and simplifying, we find

$$\begin{aligned}
\mathcal{S}_n(i,j,k)\Psi_n =& \frac{n-i-j-k+2}{n+2}\frac{n-i-j-k+1}{n+1}\left(s_{AB/ab}i + s_{Ab/ab}j + s_{aB/ab}k\right)\Psi_{n+2}(i,j,k) \\
&+ \frac{i+1}{n+2}\frac{n-i-j-k+1}{n+1}(s_{AB/AB}i + s_{AB/Ab}j + s_{AB/aB}k + s_{Ab/ab}j \\
&\quad + s_{aB/ab}k - s_{AB/ab}(n+j+k))\Psi_{n+2}(i+1,j,k) \\
&+ \frac{i+2}{n+2}\frac{i+1}{n+1}(s_{AB/Ab}j + s_{AB/aB}k + s_{AB/ab}(n-i-j-k) - s_{AB/AB}(n-i))\Psi_{n+2}(i+2,j,k) \\
&+ \frac{i+1}{n+2}\frac{j+1}{n+2}(s_{AB/AB}i + s_{AB/aB}k + s_{AB/ab}(n-i-j-k) + s_{Ab/Ab}j \\
&\quad + s_{Ab/aB}k + s_{Ab/ab}(n-i-j-k) - s_{AB/Ab}(2n-i-j))\Psi_{n+2}(i+1,j+1,k) \\
&+ \frac{i+1}{n+2}\frac{k+1}{n+1}(s_{AB/AB}i + s_{AB/Ab}j + s_{AB/ab}(n-i-j-k) + s_{Ab/aB}j \\
&\quad + s_{aB/aB}k + s_{aB/ab}(n-i-j-k) - s_{AB/aB}(2n-i-k))\Psi_{n+2}(i+1,j,k+1) \\
&+ \frac{j+1}{n+2}\frac{n-i-j-k+1}{n+1}(s_{AB/Ab}i + s_{AB/ab}i + s_{Ab/Ab}j + s_{Ab/aB}k \\
&\quad + s_{aB/ab}k - s_{Ab/ab}(n+i+k))\Psi_{n+2}(i,j+1,k) \\
&+ \frac{j+2}{n+2}\frac{j+1}{n+1}(s_{AB/Ab}i + s_{Ab/aB}k + s_{Ab/ab}(n-i-j-k) - s_{Ab/Ab}(n-j))\Psi_{n+2}(i,j+2,k) \\
&+ \frac{j+1}{n+2}\frac{k+1}{n+1}(s_{AB/Ab}i + s_{AB/aB}i + s_{Ab/Ab}j + s_{Ab/ab}(n-i-j-k) \\
&\quad + s_{aB/aB}k + s_{aB/ab}(n-i-j-k) - s_{Ab/aB}(2n-j-k))\Psi_{n+2}(i,j+1,k+1) \\
&+ \frac{k+1}{n+2}\frac{n-i-j-k+1}{n+1}(s_{AB/aB}i + s_{AB/ab}i + s_{Ab/aB}j + s_{Ab/ab}j \\
&\quad + s_{aB/aB}k - s_{aB/ab}(n+i+j))\Psi_{n+2}(i,j,k+1) \\
&+ \frac{k+2}{n+2}\frac{k+1}{n+1}(s_{AB/aB}i + s_{Ab/aB}j + s_{aB/ab}(n-i-j-k) - s_{aB/aB}(n-k))\Psi_{n+2}(i,j,k+2).
\end{aligned} \tag{S8}$$

Multiplying through by $2N_{ref}$ gives us selection operators in terms of $\gamma$ instead of $s$.

#### S1.3 Validation against simulation

To assess the accuracy of the numerical approximation, I carried out discrete two-locus simulations over a range of recombination rates and selection parameterizations. Such discrete simulations track allele and haplotype frequencies in a population of a given size, with offspring generations constructed via multinomial sampling from the haplotype frequencies in the parental generation. Because we only track the two focal loci, these simulations do not contain any additional effects of linked selection (see Section S1.4 for simulations with many linked loci).

Simulations had diploid population sizes of 5000 and I drew haploid sample sizes of 50 (25 diploids), with recombination rates varying from 0 to 0.001 (or $\rho = 0 - 20$). I tested selection coefficients of $s$ of 0, $-1 \times 10^{-5}$, $-2 \times 10^{-4}$, and $-2 \times 10^{-3}$ (or $\gamma = 0$, $-0.1$, $-2$, and $-20$). Epistasis was set to $\epsilon = 0$, $-0.5$, and 0.5, and dominance was set to either $h = 0.5$ (additive) or 0.1 (recessive). Conditional distributions of the two-locus sampling distribution show good agreement between numerical solutions and simulations, and residuals do not show any directional bias (Figures S2–S6). The Supporting Figures includes a handful of these comparisons, and the full set of comparisons (over 40 in total across the range of parameters listed above), simulation, and plotting scripts in Python are available at `https://github.com/apragsdale/two_locus_selection`.

To assess the accuracy under dominance, I simulated under the Roze (2021) scenarios ($N = 1000$, $r = 0.0001$ and 0.001, and $sh = -0.0001$, $-0.0004$, $-0.001$, and $-0.01$). Note that $sh = -0.01$ is extremely strong selection, especially for small $h$, and `moments` was unstable in that parameter regime. For weak and moderate selection, there good agreement between simulated data and `moments` solutions (Figure 4D, E).

##### S1.4 Background selection and associative overdominance

To explore the effects of many linked selected loci on patterns of LD, I used `fwdpy11` (Thornton, 2014, 2019) to simulate large genomic segments with many selected mutations. Each simulation had population size of $N = 1000$, a sequence length of 1Mb, with mutation and recombination rates both $u = r = 10^{-8}$. I considered four scenarios for selected mutaions: additive selection $h = 0.5$ with $s = -0.0005$ and $s = -0.01$ ($\gamma = -1$ and $-20$), and recessivity $h = 0.0$ or $h = 0.1$ with the same selection coefficients. These sets of parameters should give rise to strong effects of linked selection, as there are many cosegregating, tightly linked selected variants. For each setting, I examined site-frequency spectra and LD decay for both selected and neutral mutations across the region.

For each simulation, I ran a large number of replicates (100 independent replicates, and within each replicate, I sampled 25 diploid individuals every 100 generations for a total of 20,000 samples from each replicate, after burn-in of $100N$ generations). Additive selection caused minor distortions to the SFS, and reduced observed $\sigma_d^1$ and $\sigma_d^2$ in the case of weak selection ($\gamma = -1$) (Figure S7). LD was largely unaffected in the stronger selection scenario, consistent with intuition that interference (including interference from additional linked sites) is most noticeable for $s \approx 1/N$ (Figure S8).

Larger deviations from two-locus expectations were observed for recessive mutations. For recessive weakly deleterious mutations (Figure S9), there is a large excess of neutral and selected mutations at common frequencies (due to associative overdominance (Zhao and Charlesworth, 2016)). $\sigma_d^2$ for neutral and selected mutations is also increased over two-locus expectations. Stronger recessive mutations (Figure S10) also show distorted allele frequencies, though the distortion to LD is not as strong as observed for weak recessive variants.

##### S1.5 Frequency dependence of signed LD

Mutations at different frequencies are expected to have different signs and magnitudes of LD. For example, rare mutations show highly positive LD, even for pairs of neutral mutations (Good, 2022). Mutations of different functional classes (synonymous, missense, and loss-of-function, for example) are expected to segregate at different average frequencies, and average selection pressures are observed to be greater within annotated conserved domains, with stronger purifying selection driving allele frqeuencies to relatively lower levels among missense and loss-of-function variants. Since mutations of difference classes and locations have different average frequencies, this may drive differences in patterns of LD when comparing between them.

One approach to lessen the impact of differing allele frequencies when comparing between classes of mutations is to condition on allele frequencies. From sampling data with observed derived allele counts, it is simplest to condition on those allele counts (e.g.,$n_A, n_B = 2$, Garcia and Lohmueller, 2021), and to compute $\sigma_d^1$ and $\sigma_d^2$ from pairs of mutations that satisfy that condition. In the simulations with haploid sample sizes $n = 50$ (Figures 6, 7, S11–S16), I partitioned allele counts as uncommon ($n_A, n_B \leq 4$, or $\hat{f_A}, \hat{f_B} \leq 0.08$), or common ($n_A, n_B \geq 5$). In the 1000 Genomes Project Consortium *et al.* (2015) data, in which most populations have diploid sample sizes between 80 and 100, I considered $n_A, n_B \geq 2$, $3 \leq n_A, n_B \leq 8$, and $n_A, n_B \geq 9$ (or rare, uncommon, and common ($\gtrsim 0.05$)). Functions to compute allele-count-conditioned statistics from $\Psi_n$ are packaged within `moments`.

#### S2 Data analysis

##### S2.1 DFE for missense and LOF variants

Loss-of-function (LOF) variants show a dramatic skew toward low-frequency variants across all human populations (Table S4). Here, using the folded SFS for synonymous, missense, and LOF mutations across all autosomal genes, I inferred DFEs for missense and LOF mutations independently. I considered a few different dominance coefficients to explore the effect of the assumed recessivity of the two classes of mutations.

The standard SFS approach to fitting the DFE involves first inferring a demographic history for the population using putatively neutral variants (here, synonymous mutations), and then fixing that demography and fitting a parameterized function for the distribution of selection coefficients for new mutations for the selected classes. DFE inference also requires an estimate for the total mutation rate of the different mutation classes, as much of the signal for strongly selected mutations comes from observing fewer mutations than expected given a known mutation rate (with the assumption that selection purges some fraction of strongly deleterious mutations which are unseen in the sample). Here, I fit demography and DFEs to the folded SFS from the Mende in Sierra Leone (MSL) using *moments* version 1.1.0 (Jouganous *et al.*, 2017).

I used the mutation model from Karczewski *et al.* (2020) to estimate the total mutation rate across autosomal genes ($uL$, where $u$ is the per-base mutation rate of a given mutation class, and $L$ is the total length of the coding genome). These values were $(0.1442, 0.3426, 0.0256)$ for synonymous, missense, and LOF mutations, respectively. Roughly two thirds of new mutations in coding regions are expected to be missense mutations, while only 5% of new mutations are LOF. I fit a demographic model to the synonymous variants, which included a population expansion in the deeper past and exponential growth in the recent past (Figure S27A). Using the inferred optimal scaled mutation rate, $\theta = 4N_e uL$, I estimated $Ne \approx 12,300$, and assuming an average generation time of 29 years I converted the inferred genetic units to physical units. The best-fit model had a roughly two-fold expansion 400 thousand years ago, and then exponential growth over the past 20-30 thousand years, with a current effective size of $\approx 63,000$.

Under this demographic model, I fit a gamma distribution for the distribution of fitness effects to missense and LOF mutations (Table S5). For each fit, I fixed the scaled mutation rate for each mutation class, so that $\theta_{mis} = \frac{u_{mis}}{u_{syn}}\hat{\theta}_{syn}$ and $\theta_{lof} = \frac{u_{lof}}{u_{syn}}\hat{\theta}_{syn}$, where values of $u$ were found using the GNOMAD mutation model (Karczewski *et al.*, 2020). I tested three values for the dominance coefficient $h$: 0, 0.2 and 0.5. For missense mutations, $h = 0$ gave a poor fit to the data, and $h = 0.5$ fit best among the three tested dominance coefficients. For LOF variants, $h = 0$ also fit poorly, but $h = 0.2$ and $h = 0.5$ gave similar likelihoods, highlighting that inferring dominance using the SFS is poorly constrained. Regardless of the dominance coefficient assumed, however, the vast majority of LOF variants were inferred to be strongly deleterious, with only $\sim 10\%$ of new mutations having selection coefficients on the order $1/N_e$ or less.

#### S2.2 Multinucleotide mutations and positive LD between linked synonymous variants

Multinucleotide mutations (MNMs) are complex mutational events that result in multiple mutations occurring on the same haplotype background in a single generation. Because MNMs fall on the same haplotype, those mutations will be in positive LD, and LD between those pairs that are very tightly linked will not be broken down all that rapidly. MNMs are expected to occur over relatively short distances, on the order of 10s or 100s of base pairs, making them a likely culprit of the observed positive LD among synonymous mutations at short distances.

MNM events can be easily incorporated into the moment system with a simple adjustment to the mutation operator. Instead of all mutations occurring independently in haplotypes with mutations already segregating at the other locus, some fraction of new mutations could instead occur spontaneously and create a new pair of mutations with initial counts $n_{AB} = 1$ and $n_{ab} = n-1$. We can partition $\Psi_n$ into pairs of mutations due to MNM events ($\Psi_n^{MNM}$) and those due to sequential mutation events ($\Psi_n^{ISM}$), so that $\Psi_n = \Psi_n^{MNM} + \Psi_n^{ISM}$. For given scaled mutation rates $\theta = 4N_e\mu$, $\Psi_n^{ISM} \propto \theta^2$ and $\Psi_n^{MNM} \propto p_{MNM}\theta$, where $p_{MNM}$ is the probability that a mutational event causes a multinucleotide mutation. This difference in scaling means that $\Psi_n$ is sensitive to both the overall mutation rate $\theta$ as well as the probability that a mutation is a MNM at a given distance.

##### S2.2.1 Optimization details

Here, I fit a simple exponential model for the fraction of new mutations at a given distance that give rise to a MNM event, so that $p_{MNM,d} = P(MNM|d) = Ae^{-\lambda d}$, where $d$ is the distance separating pairs of mutations in base pairs. I considered all synonymous mutations within genes in the MSL data and used the same population size history model as inferred in the DFE section above for a demographic control. This left two parameters to be fit, $A$ and $\lambda$, which I fit to the binned decay curve of $\sigma_d^1$ using the midpoint of each bin as the distance for that bin. I needed to assume an average per-base recombination rate $r$ across gene regions, and tested a number of values between $10^{-9}$ and $2 \times 10^{-8}$. Optimization was insensitive to the chosen value of $r$, because the decay of positive LD occurs rapidly. For any plausible value of $r$, $\sigma_d^1$ decays to zero well before distances between pairs have scaled recombination rates $\rho = 4N_erd \sim 1$, and expected statistics for $\rho \ll 1$ vary negligibly.

To perform optimization, I assumed a Guassian likelihood function within each bin using variances estimated via bootstrap replicates from sampling genes with replacement. A composite likelihood was computed by taking the product of likelihoods across bins. I used the estimate of $Ne \approx 12,300$ from the DFE optimization using MSL data, a coding mutation rate of $\mu = 1.33 \times 10^{-8}$ (Karczewski *et al.*, 2020), and the fraction of coding mutations that result in synonymous mutations was assumed to be (assuming a ration of nonsynonymous mutations to synonymous mutations of 2.5:1). Thus, the scaled mutation rate was $\theta = 4 \times N_e \times \mu \times \frac{1}{3.5} \approx 1.87 \times 10^{-4}$.

###### S2.2.2 MNM optimization results and discussion

In fitting the LD decay of $\sigma_d^1$, the best fit parameters were $A = 2.44 \times 10^{-5}$, and $\lambda = 0.00919$. Thus, only a small fraction of new mutations were inferred to cause MNM events ($\ll 1\%$), and $1/\lambda \approx 100$ means that the proportion of mutation events that cause MNM decays quickly within a few hundred base pairs.

However, because of the differences in scaling of $\Psi_n^{ISM}$ and $\Psi_n^{MNM}$ (with the square of $\theta$ vs linearly as $p_{MNM}\theta$), the number of *observed* pairs of mutations at a given distance that arose due to a MNM event is much larger. A rough estimate given the assumed mutation rate and inferred parameters is that $p_{MNM}\theta/\theta^2 \approx 10\%$ of pairs of synonymous mutations observed at $d \approx 0$ are due to MNM events, roughly $p_{MNM}e^{-\lambda 100}\theta/\theta^2 \approx 4\%$ at $d = 100$ base pairs, and only about $p_{MNM}e^{-\lambda 400}\theta/\theta^2 \approx 0.25\%$ at $d = 400$ base pairs. This inference is very sensitive to the assumed mutation rate and the proportion of mutations that cause synonymous vs nonsynonymous variants, and future work will be needed to refine these estimates.

##### S2.3 Grouping Thousand Genomes populations based on clustering

The large confidence intervals for measurements of signed LD could be driven by either averaging over relatively few observed pairs of mutations, or due to small sample sizes that make each individual measurement a noisy estimate of the LD for that pair of mutations in the full population. To explore the underlying cause of measurement uncertainty in the 1000 Genomes Project Consortium *et al.* (2015) data, I considered larger sets of samples by combining populations that consistently cluster together in PCA and UMAP space and have low differentiation (Diaz-Papkovich *et al.*, 2020). I took combinations of CEU/GBR, CHB/CHS, CDX/KHV, and MSL/GWD. While recognizing that residual population structure in these population combinations could alter expected LD statistics compared to the respective single-population estimates, I was more interested in the effect that increasing the sample sizes would have on estimated measurement error.

Across each of the four combinations tested, confidence intervals were roughly equivalent to those of each of the individual populations. This suggests that the limiting factor to accurate LD measurement is not sample size but rather the overall levels of diversity and number of pairs of mutations that we compare. $\mathbb{E}[D]$ is most affected by common variants, and the sample sizes of the Thousand Genomes Project data are likely sufficient to accurately estimate common allele frequencies. Adding additional samples will increase the number of rare variants that we observe, but rare variants have minimal impact on $\sigma_d^1$. Thus, the accuracy of estimates of $\sigma_d^1$ is more fundamentally limited by evolutionary history and genome biology (i.e. past population sizes, mutation and recombination rates) than by sample sizes.

### S4 Supporting tables

| Diploid genotype | General model | Simple dominance | Gene-based dominance |
|---|---|---|---|
| $AB$ / $AB$ | $1 + s_{AB/AB}$ | $1 + 2s_A + 2s_B$ | $1 + 2s$ |
| $AB$ / $Ab$ | $1 + s_{AB/Ab}$ | $1 + 2s_A + 2s_B h_B$ | $1 + 2s$ |
| $AB$ / $aB$ | $1 + s_{AB/aB}$ | $1 + 2s_A h_A + 2s_B$ | $1 + 2s$ |
| $AB$ / $ab$ | $1 + s_{AB/ab}$ | $1 + 2s_A h_A + 2s_B h_B$ | $1 + 2sh$ |
| $Ab$ / $Ab$ | $1 + s_{Ab/Ab}$ | $1 + 2s_A$ | $1 + 2s$ |
| $Ab$ / $aB$ | $1 + s_{Ab/aB}$ | $1 + 2s_A h_A + 2s_B h_B$ | $1 + 2s$ |
| $Ab$ / $ab$ | $1 + s_{Ab/ab}$ | $1 + 2s_A h_A$ | $1 + 2sh$ |
| $aB$ / $aB$ | $1 + s_{aB/aB}$ | $1 + 2s_B$ | $1 + 2s$ |
| $aB$ / $ab$ | $1 + s_{aB/ab}$ | $1 + 2s_B h_B$ | $1 + 2sh$ |
| $ab$ / $ab$ | 1 | 1 | 1 |

Table S1: **General selection model for diploids and dominance models.** Different dominance and interactive effects can be implemented by assigning the appropriate values of $s$.

| Haplotype | Fitness |
|---|---|
| $AB$ | $(1 + s_A + s_B)(1 + \epsilon)$ |
| $Ab$ | $1 + s_A$ |
| $aB$ | $1 + s_B$ |
| $ab$ | 1 |

Table S2: **Haploid epistasis model.**

| Code | Description | Region |
|---|---|---|
| ESN | Esan in Nigeria | Africa |
| GWD | Gambian in Western Divisions in the Gambia | Africa |
| LWK | Luhya in Webuye, Kenya | Africa |
| MSL | Mende in Sierra Leone | Africa |
| YRI | Yoruba in Ibadan, Nigeria | Africa |
| CEU | Utah Residents (CEPH) with Northern and Western European Ancestry | Europe |
| GBR | British in England and Scotland | Europe |
| FIN | Finnish in Finland | Europe |
| IBS | Iberian Population in Spain | Europe |
| TSI | Toscani in Italia | Europe |
| CDX | Chinese Dai in Xishuangbanna, China | East Asia |
| CHB | Han Chinese in Beijing, China | East Asia |
| CHS | Southern Han Chinese | East Asia |
| JPT | Japanese in Tokyo, Japan | East Asia |
| KHV | Kinh in Ho Chi Minh City, Vietnam | East Asia |

Table S3: **Thousand Genomes Project population descriptions for populations used in this study.**

Table S4: **Tamija's $D$ for classes of coding mutations.** Here, we partitioned by synonymous, missense, and nonsense mutationsi, and considered all mutations gene-wide, mutations falling within annotated conserved elements, and mutations falling outside of those elements.

| Population | Mutation type | Region | Tajima's D |
|---|---|---|---|
| ESN | Synonymous | All | -0.882 |
| | | In domain | -0.854 |
| | | Not in domain | -0.921 |
| | Missense | All | -1.414 |
| | | In domain | -1.535 |
| | | Not in domain | -1.293 |
| | Loss of function | All | -1.483 |
| | | In domain | -2.156 |
| | | Not in domain | -1.282 |
| GWD | Synonymous | All | -1.011 |
| | | In domain | -0.981 |
| | | Not in domain | -1.052 |
| | Missense | All | -1.566 |
| | | In domain | -1.678 |
| | | Not in domain | -1.452 |
| | Loss of function | All | -1.697 |
| | | In domain | -2.328 |
| | | Not in domain | -1.501 |
| LWK | Synonymous | All | -1.109 |
| | | In domain | -1.088 |
| | | Not in domain | -1.139 |
| | Missense | All | -1.589 |
| | | In domain | -1.700 |
| | | Not in domain | -1.477 |
| | Loss of function | All | -1.666 |
| | | In domain | -2.278 |
| | | Not in domain | -1.477 |
| MSL | Synonymous | All | -0.983 |
| | | In domain | -0.959 |
| | | Not in domain | -1.017 |
| | Missense | All | -1.501 |
| | | In domain | -1.603 |
| | | Not in domain | -1.400 |
| | Loss of function | All | -1.559 |
| | | In domain | -2.303 |
| | | Not in domain | -1.332 |
| YRI | Synonymous | All | -0.928 |
| | | In domain | -0.898 |
| | | Not in domain | -0.971 |
| | Missense | All | -1.467 |
| | | In domain | -1.586 |
| | | Not in domain | -1.348 |
| | Loss of function | All | -1.624 |
| | | In domain | -2.237 |
| | | Not in domain | -1.424 |

Table S4: Tamija's $D$ for classes of coding mutations. *(continued)*

| Population | Mutation type | Region | Tajima's D |
|---|---|---|---|
| CEU | Synonymous | All | -0.417 |
| | | In domain | -0.392 |
| | | Not in domain | -0.452 |
| | Missense | All | -1.248 |
| | | In domain | -1.404 |
| | | Not in domain | -1.082 |
| | Loss of function | All | -1.501 |
| | | In domain | -2.196 |
| | | Not in domain | -1.280 |
| FIN | Synonymous | All | -0.058 |
| | | In domain | -0.047 |
| | | Not in domain | -0.075 |
| | Missense | All | -0.883 |
| | | In domain | -1.048 |
| | | Not in domain | -0.710 |
| | Loss of function | All | -1.200 |
| | | In domain | -2.034 |
| | | Not in domain | -0.906 |
| GBR | Synonymous | All | -0.319 |
| | | In domain | -0.300 |
| | | Not in domain | -0.345 |
| | Missense | All | -1.120 |
| | | In domain | -1.276 |
| | | Not in domain | -0.954 |
| | Loss of function | All | -1.313 |
| | | In domain | -2.178 |
| | | Not in domain | -0.997 |
| IBS | Synonymous | All | -0.689 |
| | | In domain | -0.664 |
| | | Not in domain | -0.724 |
| | Missense | All | -1.424 |
| | | In domain | -1.560 |
| | | Not in domain | -1.279 |
| | Loss of function | All | -1.636 |
| | | In domain | -2.349 |
| | | Not in domain | -1.378 |
| TSI | Synonymous | All | -0.650 |
| | | In domain | -0.625 |
| | | Not in domain | -0.685 |
| | Missense | All | -1.422 |
| | | In domain | -1.568 |
| | | Not in domain | -1.266 |
| | Loss of function | All | -1.655 |
| | | In domain | -2.349 |
| | | Not in domain | -1.397 |
| CDX | Synonymous | All | -0.374 |
| | | In domain | -0.366 |
| | | Not in domain | -0.385 |

Table S4: Tamija's $D$ for classes of coding mutations. *(continued)*

| Population | Mutation type | Region | Tajima's D |
|---|---|---|---|
| | Missense | All | -1.179 |
| | | In domain | -1.323 |
| | | Not in domain | -1.026 |
| | Loss of function | All | -1.360 |
| | | In domain | -2.194 |
| | | Not in domain | -1.062 |
| CHB | Synonymous | All | -0.598 |
| | | In domain | -0.593 |
| | | Not in domain | -0.606 |
| | Missense | All | -1.389 |
| | | In domain | -1.528 |
| | | Not in domain | -1.239 |
| | Loss of function | All | -1.586 |
| | | In domain | -2.344 |
| | | Not in domain | -1.298 |
| CHS | Synonymous | All | -0.544 |
| | | In domain | -0.545 |
| | | Not in domain | -0.544 |
| | Missense | All | -1.334 |
| | | In domain | -1.499 |
| | | Not in domain | -1.150 |
| | Loss of function | All | -1.559 |
| | | In domain | -2.290 |
| | | Not in domain | -1.292 |
| JPT | Synonymous | All | -0.371 |
| | | In domain | -0.368 |
| | | Not in domain | -0.376 |
| | Missense | All | -1.194 |
| | | In domain | -1.355 |
| | | Not in domain | -1.019 |
| | Loss of function | All | -1.410 |
| | | In domain | -2.272 |
| | | Not in domain | -1.086 |
| KHV | Synonymous | All | -0.576 |
| | | In domain | -0.562 |
| | | Not in domain | -0.596 |
| | Missense | All | -1.346 |
| | | In domain | -1.473 |
| | | Not in domain | -1.210 |
| | Loss of function | All | -1.535 |
| | | In domain | -2.294 |
| | | Not in domain | -1.269 |

| Class | $h$ | shape | scale | LL | $[0, 10^{-5})$ | $[10^{-5}, 10^{-4})$ | $[10^{-4}, 10^{-3})$ | $[10^{-3}, 10^{-2})$ | $[10^{-2}, \infty)$ |
|---|---|---|---|---|---|---|---|---|---|
| Missense | 0.0 | 0.093 | 768505 | -678.2 | 0.260 | 0.062 | 0.077 | 0.096 | 0.505 |
| | 0.2 | 0.138 | 6660 | -416.7 | 0.260 | 0.098 | 0.134 | 0.182 | 0.327 |
| | 0.5 | 0.147 | 2117 | -392.0 | 0.282 | 0.114 | 0.159 | 0.214 | 0.231 |
| LOF | 0.0 | 0.132 | 99999054 | -248.3 | 0.077 | 0.028 | 0.037 | 0.051 | 0.807 |
| | 0.2 | 0.177 | 477994 | -226.7 | 0.083 | 0.042 | 0.063 | 0.095 | 0.717 |
| | 0.5 | 0.188 | 121419 | -224.2 | 0.092 | 0.050 | 0.077 | 0.119 | 0.662 |

Table S5: **Distribution of fitness effects inferred for nonsynonymous mutations.** DFEs inferred for missense and loss-of-function variants in MSL for varying values of $h$. General patterns are consistent across different chosen values of $h$, although $h = 0$ results in poorer fits for both missense and LOF variants. Columns to the right of the log-likelihood (LL) column show proportions of new mutations with $|s|$ in each given bin.

| Population | Syn. $\sigma_d^1$ (std. err.) | Mis. $\sigma_d^1$ (std. err.) | LOF $\sigma_d^1$ (std. err.) |
|---|---|---|---|
| ESN | | | |
| No matching | 0.0186 (0.0150) | 0.0224 (0.0169) | -0.1552 (0.5320) |
| Matched dist. | 0.0186 (0.0155) | 0.0215 (0.0156) | -0.3079 (0.5046) |
| Matched freq. | 0.0186 (0.0150) | 0.0185 (0.0175) | -0.1706 (0.4776) |
| Dist. and freq. | 0.0186 (0.0144) | 0.0164 (0.0147) | 0.1434 (0.4557) |
| GWD | | | |
| No matching | 0.0235 (0.0153) | 0.0211 (0.0165) | -0.2019 (0.4006) |
| Matched dist. | 0.0235 (0.0152) | 0.0195 (0.0154) | -0.1307 (0.4310) |
| Matched freq. | 0.0235 (0.0164) | 0.0158 (0.0182) | -0.1946 (0.3471) |
| Dist. and freq. | 0.0235 (0.0144) | 0.0105 (0.0146) | -0.0330 (0.3348) |
| LWK | | | |
| No matching | 0.0224 (0.0148) | 0.0215 (0.0178) | -0.1327 (0.5109) |
| Matched dist. | 0.0224 (0.0157) | 0.0202 (0.0175) | -0.1880 (0.5058) |
| Matched freq. | 0.0224 (0.0149) | 0.0187 (0.0198) | -0.1510 (0.4780) |
| Dist. and freq. | 0.0224 (0.0137) | 0.0138 (0.0150) | -0.0422 (0.4287) |
| MSL | | | |
| No matching | 0.0170 (0.0153) | 0.0215 (0.0146) | -0.1851 (0.4233) |
| Matched dist. | 0.0170 (0.0146) | 0.0206 (0.0154) | -0.0342 (0.3908) |
| Matched freq. | 0.0170 (0.0161) | 0.0170 (0.0175) | -0.1204 (0.3335) |
| Dist. and freq. | 0.0170 (0.0145) | 0.0141 (0.0144) | -0.1337 (0.3619) |
| YRI | | | |
| No matching | 0.0254 (0.0165) | 0.0220 (0.0187) | -0.0962 (0.3847) |
| Matched dist. | 0.0254 (0.0169) | 0.0200 (0.0187) | -0.1982 (0.4178) |
| Matched freq. | 0.0254 (0.0172) | 0.0195 (0.0232) | -0.1462 (0.3848) |
| Dist. and freq. | 0.0254 (0.0152) | 0.0147 (0.0171) | -0.0006 (0.4040) |

Table S6: **Signed LD in AFR-labeled populations matched to distances and allele frequencies of synonymous mutations.** Gene-wide signed LD is observed to be at similar slightly positive levels between pairs of synonymous and pairs of missense variants. This observation is consistest when weighting missense variants to match distances between pairs of synonymous variants and when weighting to match synonymous allele frequencies, with only a slight reduction in missense $\sigma_d^1$. The distribution of distances between synonymous and missense mutations within genes are very similar (Figure S26), so matching by distances separating mutations has a minor effect on measured average LD. The large measurement uncertainty of loss-of-function mutation pairs due to few observed pairs of such mutations within the same gene causes signed LD to fluctuate when conditioning on distances and frequencies matched to synonymous mutations.

| Population | Syn. $\sigma_d^2$ (std. err.) | Mis. $\sigma_d^2$ (std. err.) | LOF $\sigma_d^2$ (std. err.) |
|---|---|---|---|
| ESN | | | |
| No matching | 0.1572 (0.0053) | 0.1621 (0.0089) | 0.1833 (0.0795) |
| Matched dist. | 0.1572 (0.0054) | 0.1586 (0.0085) | 0.1707 (0.0729) |
| Matched freq. | 0.1572 (0.0044) | 0.1748 (0.0087) | 0.1844 (0.0740) |
| Dist. and freq. | 0.1572 (0.0031) | 0.1623 (0.0047) | 0.1672 (0.0678) |
| GWD | | | |
| No matching | 0.1591 (0.0049) | 0.1641 (0.0097) | 0.1081 (0.0398) |
| Matched dist. | 0.1591 (0.0051) | 0.1623 (0.0092) | 0.1092 (0.0389) |
| Matched freq. | 0.1591 (0.0043) | 0.1794 (0.0089) | 0.1038 (0.0380) |
| Dist. and freq. | 0.1591 (0.0031) | 0.1679 (0.0051) | 0.0957 (0.0353) |
| LWK | | | |
| No matching | 0.1490 (0.0045) | 0.1562 (0.0093) | 0.1286 (0.0517) |
| Matched dist. | 0.1490 (0.0045) | 0.1529 (0.0092) | 0.1366 (0.0548) |
| Matched freq. | 0.1490 (0.0040) | 0.1697 (0.0085) | 0.1266 (0.0503) |
| Dist. and freq. | 0.1490 (0.0030) | 0.1556 (0.0047) | 0.1219 (0.0445) |
| MSL | | | |
| No matching | 0.1502 (0.0050) | 0.1570 (0.0093) | 0.1241 (0.0447) |
| Matched dist. | 0.1502 (0.0048) | 0.1527 (0.0093) | 0.1196 (0.0394) |
| Matched freq. | 0.1502 (0.0043) | 0.1701 (0.0090) | 0.1099 (0.0396) |
| Dist. and freq. | 0.1502 (0.0031) | 0.1571 (0.0050) | 0.1078 (0.0317) |
| YRI | | | |
| No matching | 0.1600 (0.0052) | 0.1650 (0.0096) | 0.1217 (0.0498) |
| Matched dist. | 0.1600 (0.0055) | 0.1615 (0.0096) | 0.1240 (0.0517) |
| Matched freq. | 0.1600 (0.0051) | 0.1801 (0.0097) | 0.1175 (0.0480) |
| Dist. and freq. | 0.1600 (0.0034) | 0.1667 (0.0051) | 0.1158 (0.0454) |

Table S7: **Squared LD in AFR-labeled populations matched to distances and allele frequencies of synonymous mutations.**

| Population | Syn. $\sigma_d^1$ (std. err.) | Mis. $\sigma_d^1$ (std. err.) | LOF $\sigma_d^1$ (std. err.) |
|---|---|---|---|
| CEU | | | |
| No matching | 0.0232 (0.0265) | 0.0278 (0.0316) | 0.0161 (0.8613) |
| Matched dist. | 0.0232 (0.0250) | 0.0291 (0.0311) | -0.0439 (0.9127) |
| Matched freq. | 0.0232 (0.0299) | 0.0208 (0.0392) | 0.0669 (1.0064) |
| Dist. and freq. | 0.0232 (0.0277) | 0.0230 (0.0308) | 0.3833 (0.6455) |
| FIN | | | |
| No matching | 0.0342 (0.0246) | 0.0248 (0.0349) | -0.0593 (0.7886) |
| Matched dist. | 0.0342 (0.0253) | 0.0252 (0.0347) | -0.4903 (0.8358) |
| Matched freq. | 0.0342 (0.0301) | 0.0166 (0.0449) | 0.0042 (0.9103) |
| Dist. and freq. | 0.0342 (0.0280) | 0.0201 (0.0337) | 0.5981 (0.6003) |
| GBR | | | |
| No matching | 0.0293 (0.0249) | 0.0351 (0.0365) | -0.2074 (0.8241) |
| Matched dist. | 0.0293 (0.0244) | 0.0352 (0.0338) | -0.3794 (0.8403) |
| Matched freq. | 0.0293 (0.0278) | 0.0330 (0.0425) | -0.1755 (1.0623) |
| Dist. and freq. | 0.0293 (0.0274) | 0.0298 (0.0297) | 0.2995 (0.6625) |
| IBS | | | |
| No matching | 0.0194 (0.0246) | 0.0358 (0.0351) | -0.2830 (0.8064) |
| Matched dist. | 0.0194 (0.0241) | 0.0358 (0.0338) | -0.5134 (0.7790) |
| Matched freq. | 0.0194 (0.0261) | 0.0360 (0.0403) | -0.3616 (0.7154) |
| Dist. and freq. | 0.0194 (0.0259) | 0.0328 (0.0291) | -0.1194 (0.6295) |
| TSI | | | |
| No matching | 0.0185 (0.0237) | 0.0394 (0.0317) | -0.1689 (0.7143) |
| Matched dist. | 0.0185 (0.0244) | 0.0416 (0.0330) | -0.4632 (0.8200) |
| Matched freq. | 0.0185 (0.0272) | 0.0331 (0.0387) | -0.2158 (0.7336) |
| Dist. and freq. | 0.0185 (0.0258) | 0.0307 (0.0288) | -0.0262 (0.6130) |

Table S8: **Signed LD in EUR-labeled populations matched to distances and allele frequencies of synonymous mutations.** Similar to the pattern observed in AFR-labeled populations (Table S6), $sigma_d^1$ between missense mutations matched to distances and allele frequencies observed in synonymous mutation pairs has only a small effect on average signed LD, and gene-wide measurements are roughy equal between these two mutation classes. Again, measurement noise for loss-of-function mutation pairs is high.

| Population | Syn. $\sigma_d^2$ (std. err.) | Mis. $\sigma_d^2$ (std. err.) | LOF $\sigma_d^2$ (std. err.) |
|---|---|---|---|
| CEU | | | |
| No matching | 0.3017 (0.0097) | 0.2909 (0.0145) | 0.1818 (0.0724) |
| Matched dist. | 0.3017 (0.0100) | 0.2918 (0.0153) | 0.1820 (0.0885) |
| Matched freq. | 0.3017 (0.0102) | 0.3111 (0.0147) | 0.1884 (0.0730) |
| Dist. and freq. | 0.3017 (0.0063) | 0.2957 (0.0086) | 0.1837 (0.0703) |
| FIN | | | |
| No matching | 0.3062 (0.0106) | 0.2887 (0.0144) | 0.1660 (0.0504) |
| Matched dist. | 0.3062 (0.0111) | 0.2881 (0.0142) | 0.1772 (0.0650) |
| Matched freq. | 0.3062 (0.0106) | 0.3101 (0.0138) | 0.1793 (0.0522) |
| Dist. and freq. | 0.3062 (0.0066) | 0.2904 (0.0090) | 0.1425 (0.0421) |
| GBR | | | |
| No matching | 0.3024 (0.0101) | 0.2918 (0.0131) | 0.1862 (0.0653) |
| Matched dist. | 0.3024 (0.0095) | 0.2922 (0.0131) | 0.1876 (0.0706) |
| Matched freq. | 0.3024 (0.0101) | 0.3128 (0.0133) | 0.2048 (0.0719) |
| Dist. and freq. | 0.3024 (0.0065) | 0.2971 (0.0080) | 0.1881 (0.0684) |
| IBS | | | |
| No matching | 0.2953 (0.0087) | 0.2782 (0.0127) | 0.1940 (0.0728) |
| Matched dist. | 0.2953 (0.0089) | 0.2781 (0.0124) | 0.2066 (0.0752) |
| Matched freq. | 0.2953 (0.0084) | 0.3023 (0.0119) | 0.1851 (0.0663) |
| Dist. and freq. | 0.2953 (0.0058) | 0.2865 (0.0074) | 0.2008 (0.0660) |
| TSI | | | |
| No matching | 0.2987 (0.0121) | 0.2875 (0.0149) | 0.1724 (0.0414) |
| Matched dist. | 0.2987 (0.0119) | 0.2873 (0.0147) | 0.1834 (0.0563) |
| Matched freq. | 0.2987 (0.0124) | 0.3093 (0.0145) | 0.1661 (0.0413) |
| Dist. and freq. | 0.2987 (0.0068) | 0.2916 (0.0089) | 0.1748 (0.0703) |

Table S9: **Squared LD in EUR-labeled populations matched to distances and allele frequencies of synonymous mutations.**

| Population | Syn. $\sigma_d^1$ (std. err.) | Mis. $\sigma_d^1$ (std. err.) | LOF $\sigma_d^1$ (std. err.) |
|---|---|---|---|
| CDX | | | |
| No matching | 0.0316 (0.0265) | 0.0372 (0.0403) | -0.6390 (0.7834) |
| Matched dist. | 0.0316 (0.0257) | 0.0389 (0.0419) | -0.2830 (1.0085) |
| Matched freq. | 0.0316 (0.0277) | 0.0410 (0.0458) | -0.6881 (0.7660) |
| Dist. and freq. | 0.0316 (0.0272) | 0.0405 (0.0329) | -0.4208 (0.6103) |
| CHB | | | |
| No matching | 0.0321 (0.0278) | 0.0284 (0.0384) | -0.5260 (0.4933) |
| Matched dist. | 0.0321 (0.0268) | 0.0303 (0.0382) | -0.3899 (0.5167) |
| Matched freq. | 0.0321 (0.0276) | 0.0298 (0.0434) | -0.6868 (0.4521) |
| Dist. and freq. | 0.0321 (0.0263) | 0.0321 (0.0316) | -0.5496 (0.5710) |
| CHS | | | |
| No matching | 0.0489 (0.0272) | 0.0394 (0.0413) | -0.2479 (0.7257) |
| Matched dist. | 0.0489 (0.0280) | 0.0413 (0.0405) | 0.0569 (0.8391) |
| Matched freq. | 0.0489 (0.0272) | 0.0363 (0.0406) | -0.2869 (0.7035) |
| Dist. and freq. | 0.0489 (0.0276) | 0.0358 (0.0308) | -0.2268 (0.7151) |
| JPT | | | |
| No matching | 0.0219 (0.0267) | 0.0274 (0.0361) | -0.1412 (0.6743) |
| Matched dist. | 0.0219 (0.0269) | 0.0291 (0.0372) | 0.1718 (0.7430) |
| Matched freq. | 0.0219 (0.0270) | 0.0263 (0.0374) | -0.2164 (0.6342) |
| Dist. and freq. | 0.0219 (0.0281) | 0.0274 (0.0294) | -0.3457 (0.7562) |
| KHV | | | |
| No matching | 0.0333 (0.0272) | 0.0352 (0.0407) | -0.6546 (0.8281) |
| Matched dist. | 0.0333 (0.0268) | 0.0359 (0.0386) | -0.6442 (0.8931) |
| Matched freq. | 0.0333 (0.0278) | 0.0376 (0.0473) | -0.7193 (0.7782) |
| Dist. and freq. | 0.0333 (0.0270) | 0.0369 (0.0328) | -0.5834 (0.6057) |

Table S10: **Signed LD in EUR-labeled populations matched to distances and allele frequencies of synonymous mutations.** Similar to the pattern observed in AFR-labeled populations (Table S6), $sigma_d^1$ between missense mutations matched to distances and allele frequencies observed in synonymous mutation pairs has only a small effect on average signed LD, and gene-wide measurements are roughly equal between these two mutation classes. Again, measurement noise for loss-of-function mutation pairs is high.

| Population | Syn. $\sigma_d^2$ (std. err.) | Mis. $\sigma_d^2$ (std. err.) | LOF $\sigma_d^2$ (std. err.) |
|---|---|---|---|
| CDX | | | |
| No matching | 0.3162 (0.0094) | 0.3220 (0.0208) | 0.2857 (0.0978) |
| Matched dist. | 0.3162 (0.0096) | 0.3218 (0.0194) | 0.3072 (0.0863) |
| Matched freq. | 0.3162 (0.0080) | 0.3397 (0.0176) | 0.2730 (0.0889) |
| Dist. and freq. | 0.3162 (0.0064) | 0.3215 (0.0109) | 0.2611 (0.0641) |
| CHB | | | |
| No matching | 0.3144 (0.0110) | 0.3206 (0.0237) | 0.2301 (0.0734) |
| Matched dist. | 0.3144 (0.0114) | 0.3194 (0.0232) | 0.2408 (0.0789) |
| Matched freq. | 0.3144 (0.0090) | 0.3389 (0.0217) | 0.2622 (0.0906) |
| Dist. and freq. | 0.3144 (0.0068) | 0.3262 (0.0125) | 0.2129 (0.0850) |
| CHS | | | |
| No matching | 0.3250 (0.0105) | 0.3255 (0.0194) | 0.2294 (0.0623) |
| Matched dist. | 0.3250 (0.0107) | 0.3288 (0.0195) | 0.2358 (0.0698) |
| Matched freq. | 0.3250 (0.0090) | 0.3417 (0.0182) | 0.2201 (0.0579) |
| Dist. and freq. | 0.3250 (0.0069) | 0.3265 (0.0122) | 0.2604 (0.0631) |
| JPT | | | |
| No matching | 0.3226 (0.0102) | 0.3164 (0.0210) | 0.1971 (0.0577) |
| Matched dist. | 0.3226 (0.0099) | 0.3198 (0.0206) | 0.1904 (0.0582) |
| Matched freq. | 0.3226 (0.0087) | 0.3345 (0.0202) | 0.1914 (0.0583) |
| Dist. and freq. | 0.3226 (0.0069) | 0.3216 (0.0136) | 0.2646 (0.0633) |
| KHV | | | |
| No matching | 0.3136 (0.0101) | 0.3139 (0.0189) | 0.2159 (0.1185) |
| Matched dist. | 0.3136 (0.0099) | 0.3116 (0.0197) | 0.2282 (0.1104) |
| Matched freq. | 0.3136 (0.0080) | 0.3316 (0.0169) | 0.2064 (0.1021) |
| Dist. and freq. | 0.3136 (0.0063) | 0.3149 (0.0115) | 0.1889 (0.0711) |

Table S11: **Squared LD in EAS-labeled populations matched to distances and allele frequencies of synonymous mutations.**

### S5 Supporting figures

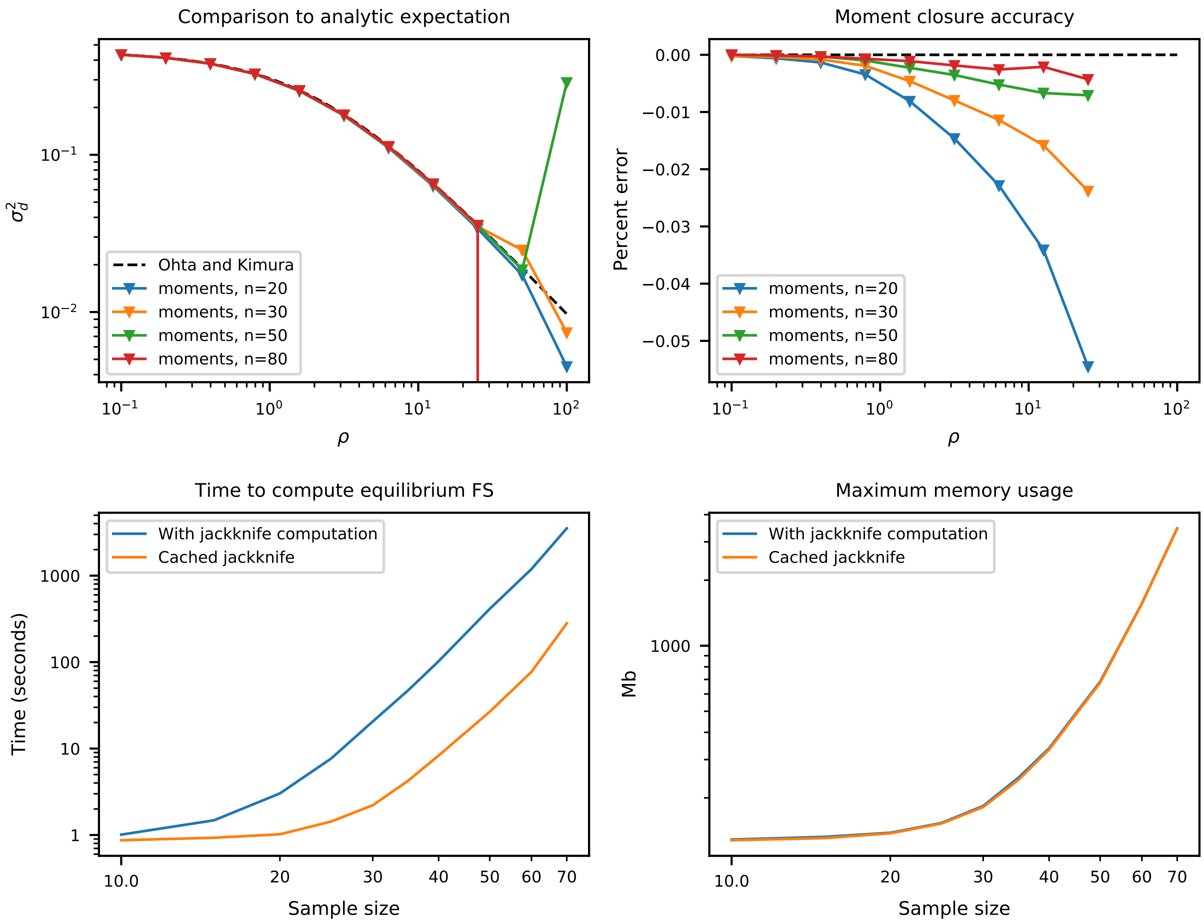

Figure S1: **Accuracy of the jackknife approximation and runtime.** Small sample sizes can lead to large error in the moment-closure approximation for larger recombination distances or selection coefficients. Generally, the jackknife approximation breaks down for recombination rates greater than $\rho \gtrsim 50$, depending on the sample size. While increasing sample size leads to more accurate solutions, it comes at the cost of both increased runtime and memory usage. Most analyses in this paper used sample sizes between 40 and 80.

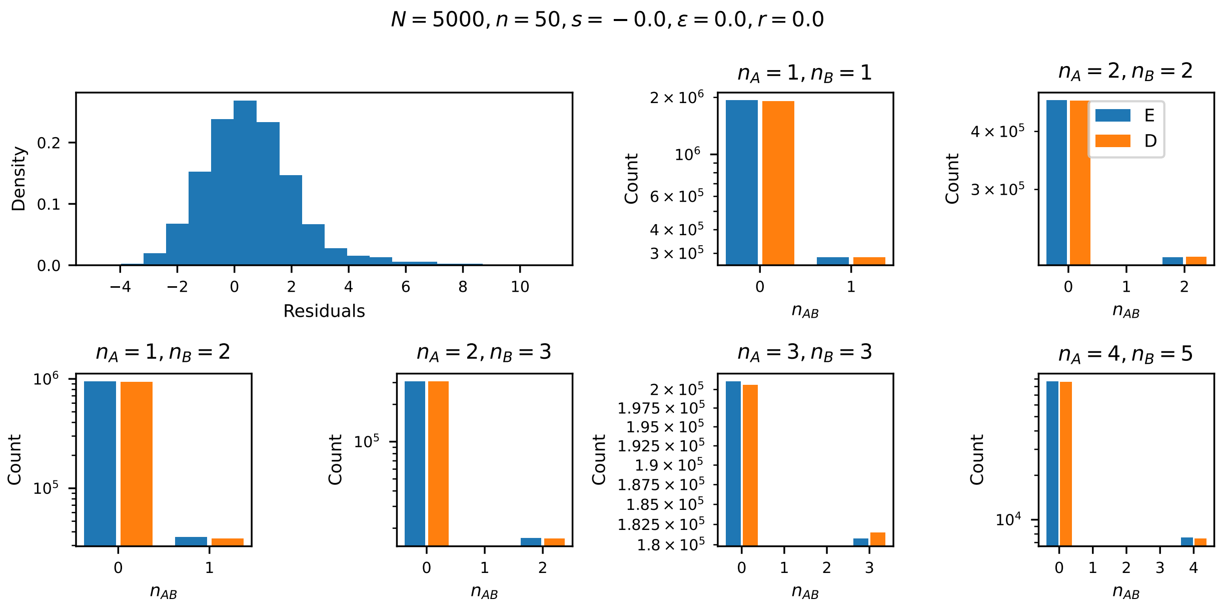

Figure S2: **Comparison between numerics and simulations for fully linked neutral loci.** With no recombination ($r = 0$) and no selection, the moments solution is exact, and computing $sigma_d^2$ from $\Psi_n$ gives 5/11, the analytic solution from Ohta and Kimura (1969). Discrepancies between numerical expectations and the simulations are due to simulation noise, with Anscombe Poisson residuals roughly normally distributed around zero (Pierce and Schafer, 1986). In the bar plots, "E" (blue) are expectations from `moments` and "D" are simulated data.

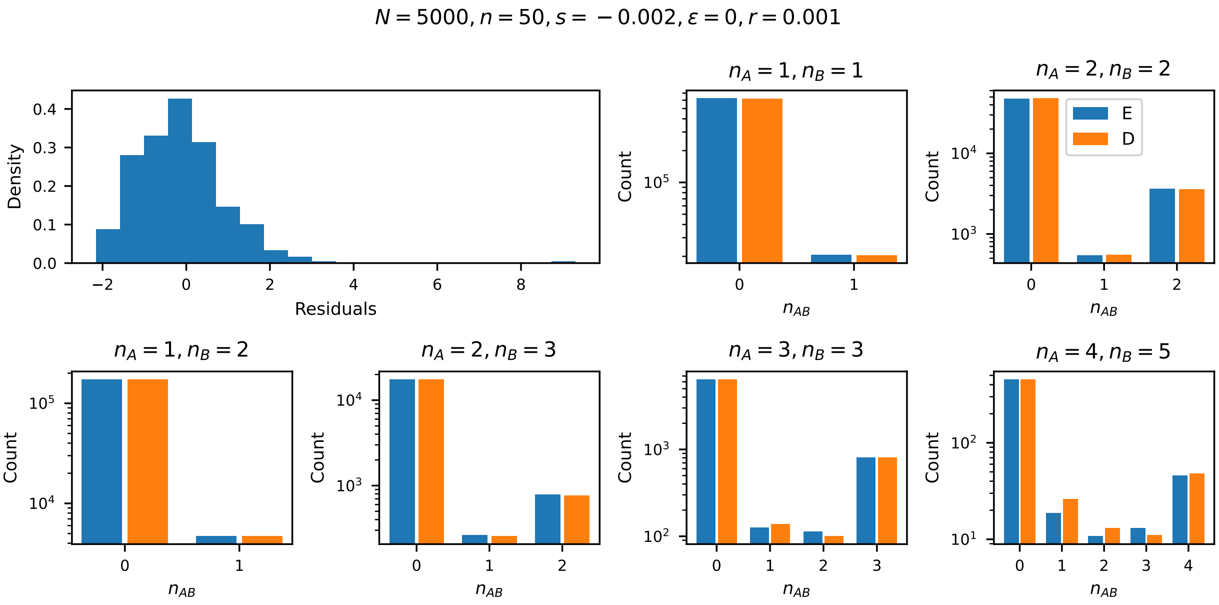

Figure S3: **Comparison to simulations for loosely linked additively selected loci.** $Ne = 5000$, $r = 0.001$, and $s = -0.002$ ($\rho = 20$ and $\gamma = -20$ at both loci), with no epistatic interactions ($\epsilon = 0$). E: `moments` expectations, D: simulated data.

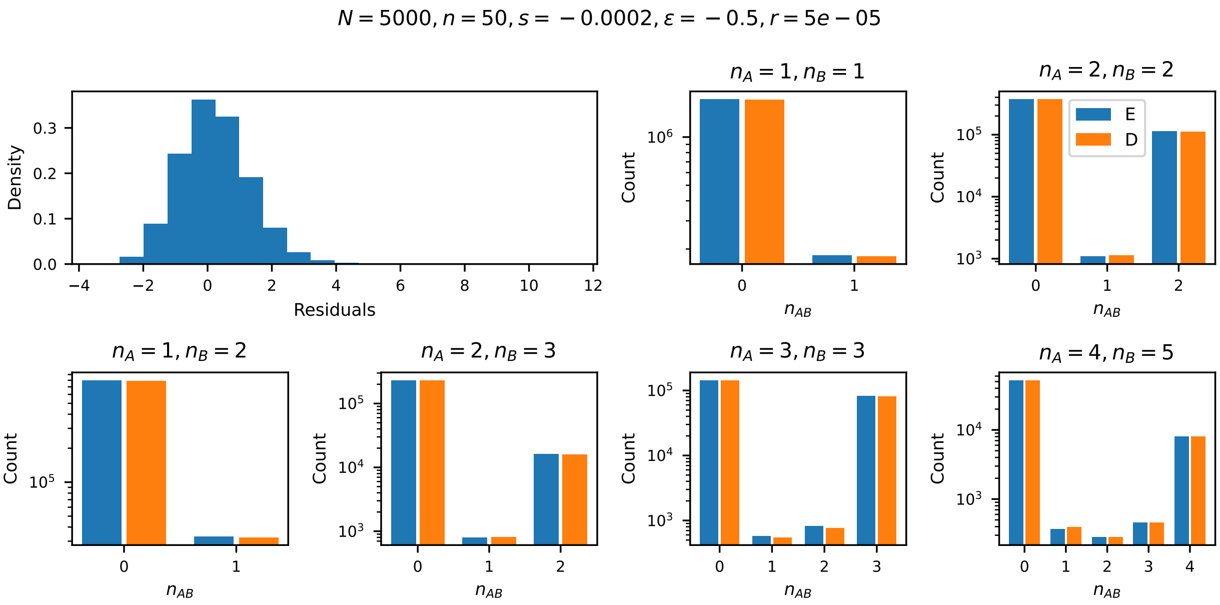

Figure S4: **Comparison to simulations with antagonistic epistasis.** $Ne = 5000$, $r = 0.00005$, $s = -0.0002$ ($\rho = 1$ and $\gamma = -2$ at both loci), and $\epsilon = -0.5$. E: `moments` expectations, D: simulated data.

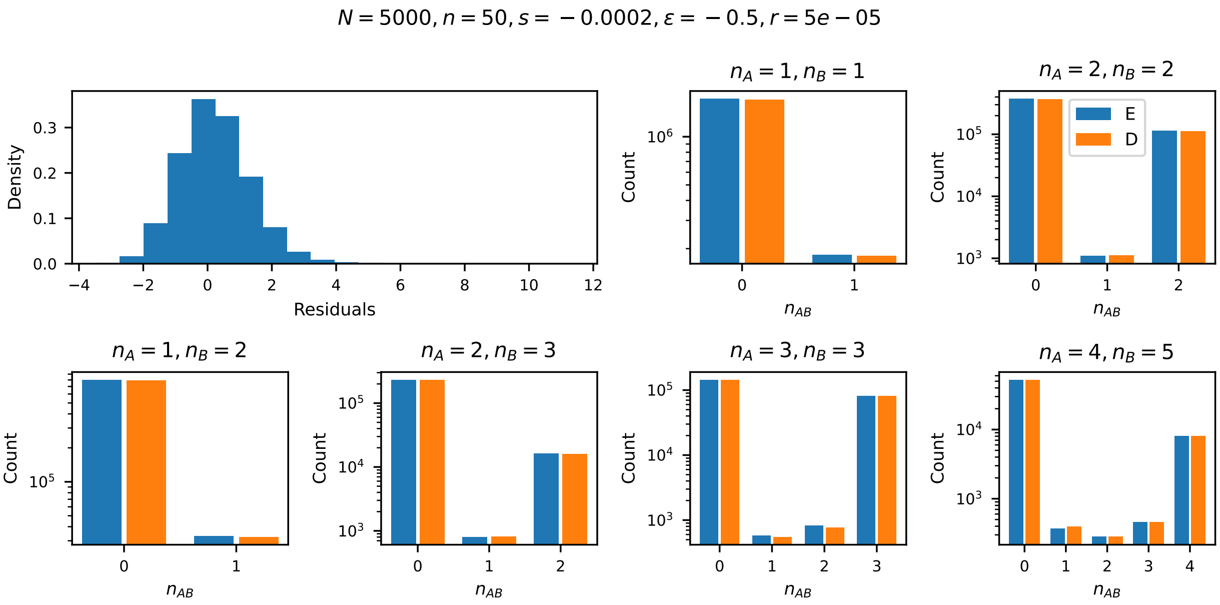

Figure S5: **Comparison to simulations with synergistic epistasis.** $Ne = 5000$, $r = 0.00005$, $s = -0.0002$ ($\rho = 1$ and $\gamma = -2$ at both loci), and $\epsilon = 0.5$. E: `moments` expectations, D: simulated data.

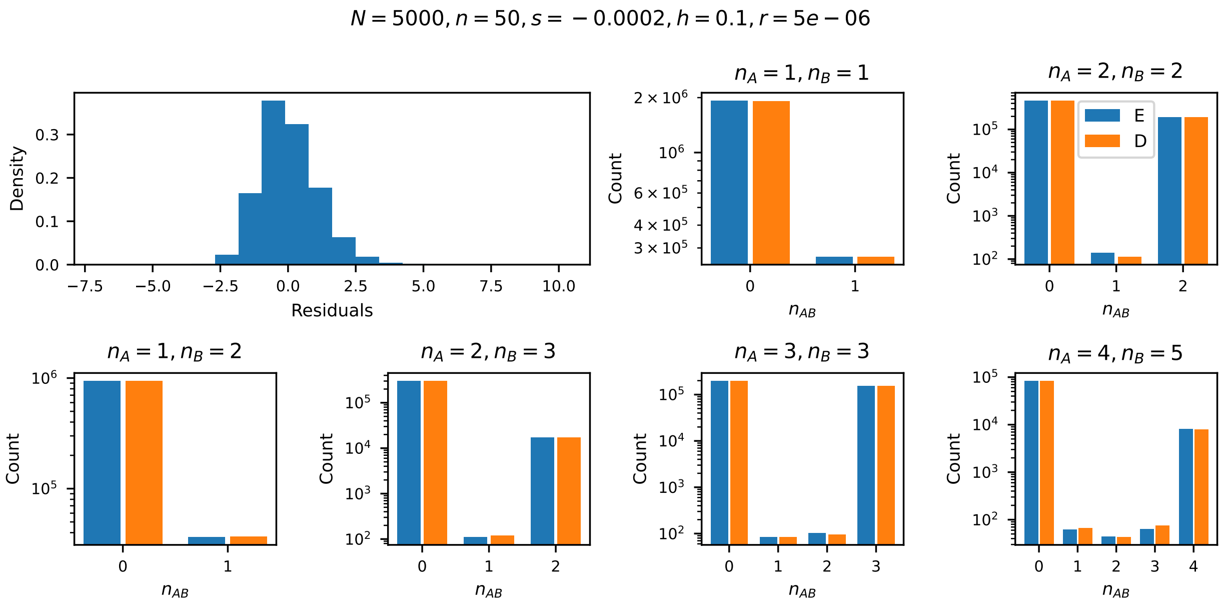

Figure S6: **Comparison to simulations with site-wise dominance.** $Ne = 5000$, $r = 5 \times 10^{-6}$, and $s = -0.0002$ ($\rho = 0.1$ and $\gamma = -2$), with $h = 0.1$ at both loci and no epistatic interactions ($\epsilon = 0$). E: `moments` expectations, D: simulated data. Additional comparisons to simulated data are availabe at `https://github.com/apragsdale/two_locus_selection`.

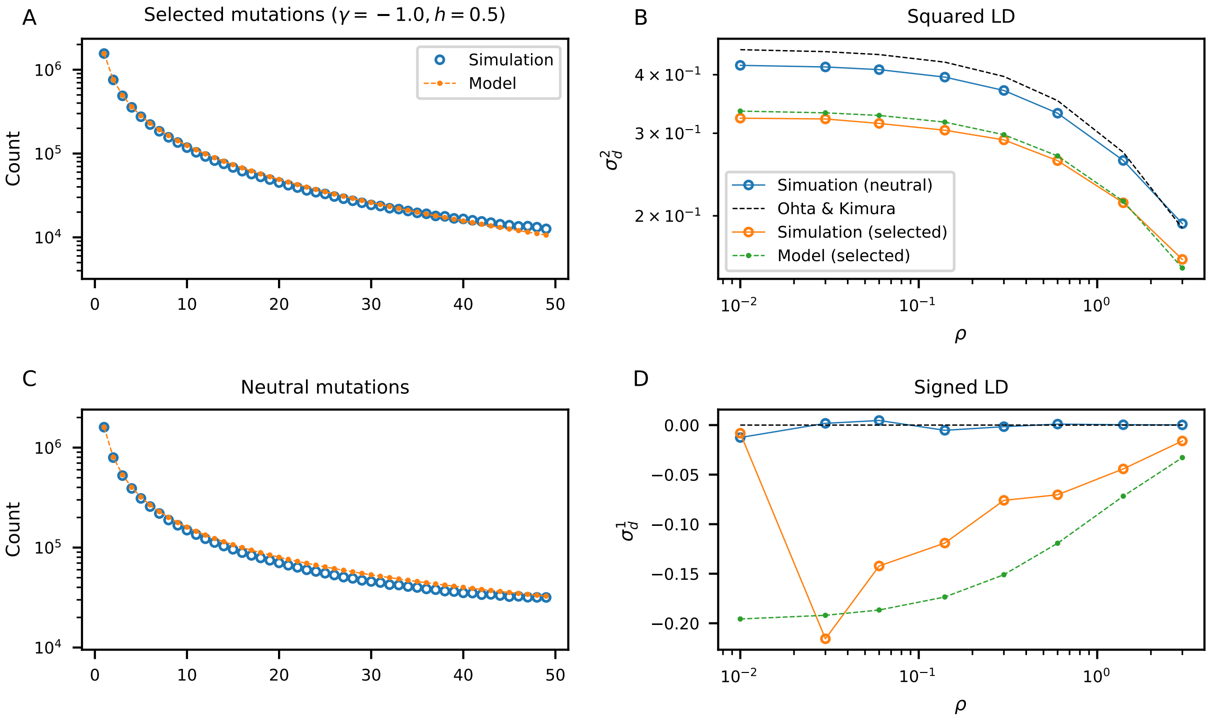

Figure S7: **Background selection with slightly deleterious mutations.** Using an individual based forward simulator (Thornton, 2019), I simulated many replicates of 1Mb regions with high mutation rates of selected mutations (see Section S1.4 for details). Here, many linked additive weakly selected mutations slightly distort the SFS for both selected and neutral mutations. $\sigma_d^2$ is reduced relative to two-locus expectations within background selection, and Hill-Robertson interference effects are slightly diminished with $\sigma_d^1$ not as strongly negative as expected.

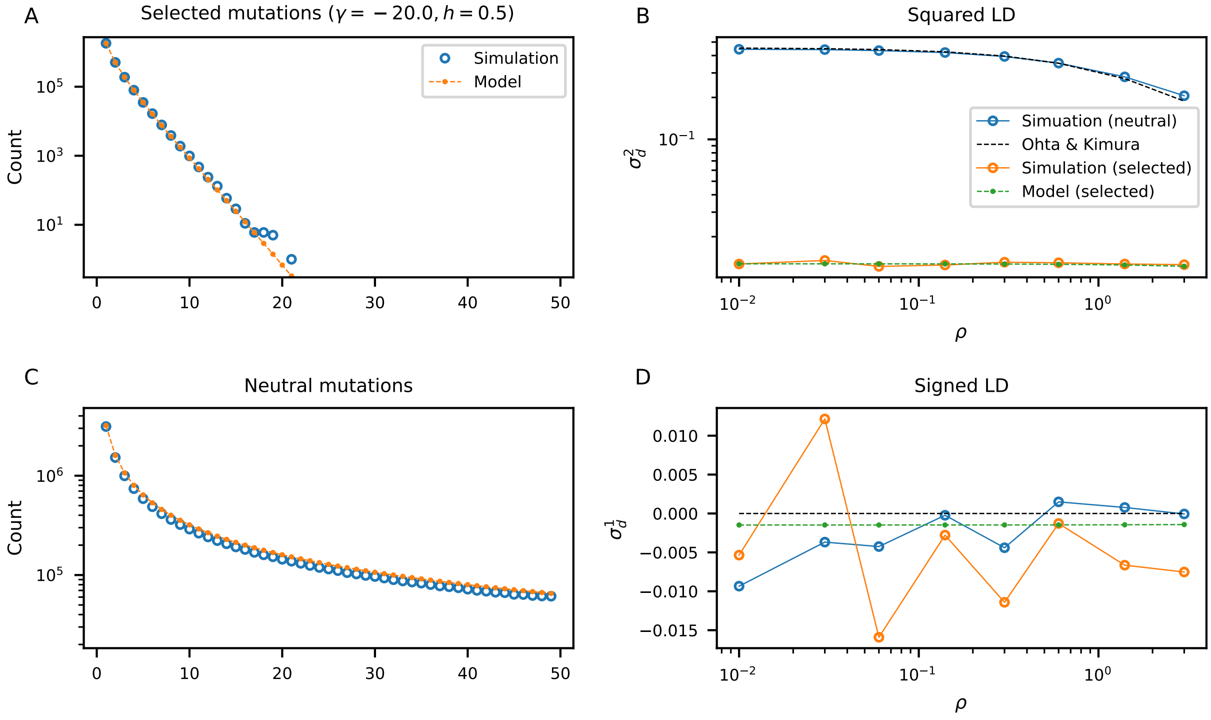

Figure S8: **Background selection with moderately deleterious mutations.** In the case of many linked strongly selected additive mutations, neither selected nor neutral statistics deviate dramatically from two-locus expectations. The SFS from the simulated data matches expectations, with a slight excess of selected mutations. LD is also largely unaffected, with $\sigma_d^2$ matching expectations and $\sigma_d^1$ fluctuating near zero as expected.

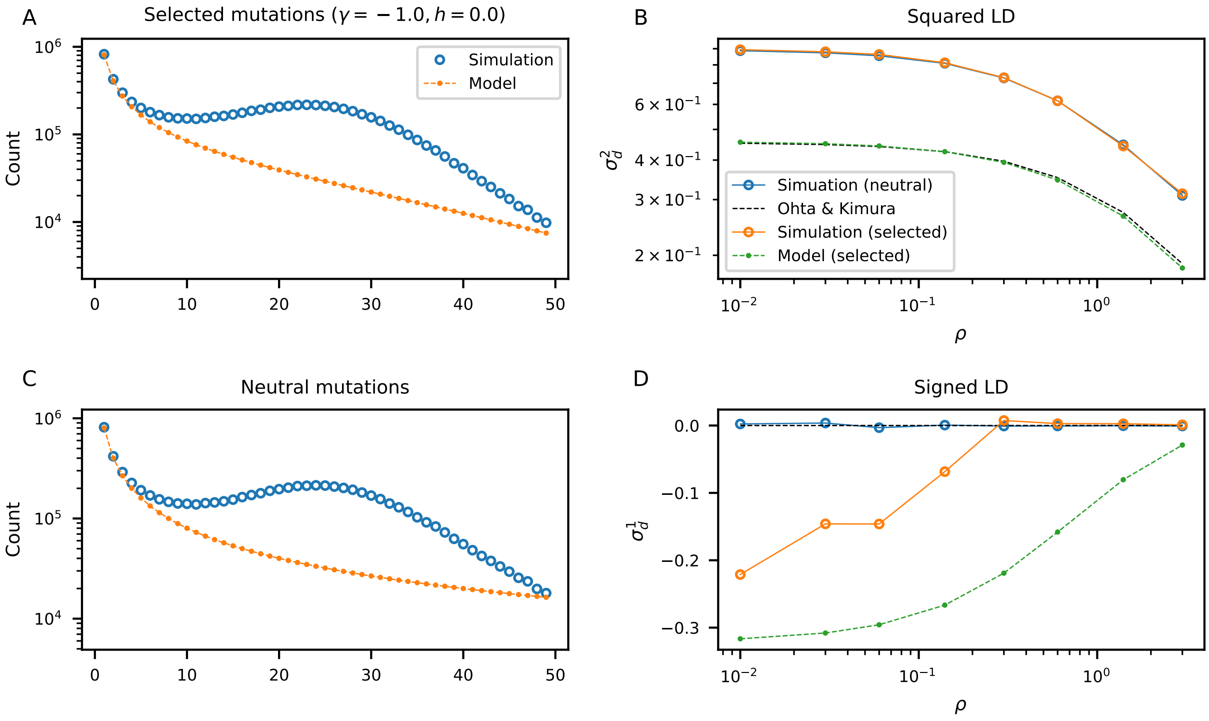

Figure S9: **Associative overdominance due to slightly deleterious recessive mutations.** The presence of many linked weakly selected recessive mutations can cause measues of diversity to have large increases over single- and two-locus expectations. The SFS for both neutral and selected mutations both show a very large excess of common variants (note the log scale), and $\sigma_d^2$ are also increased over two-locus expectations for both classes of mutations. Still, $\sigma_d^1$ remains 0 for neutral mutations and is not as strongly negative for pairs of selected mutations.

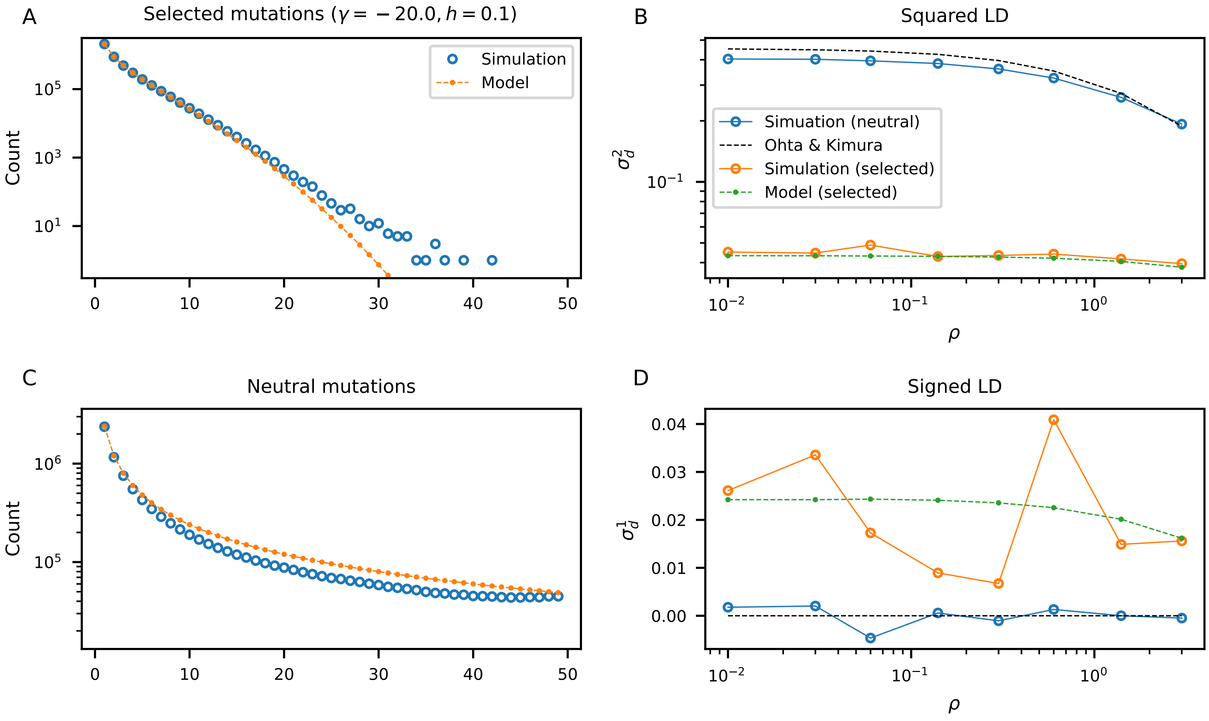

Figure S10: **Background selection with moderately deleterious recessive mutations.** More strongly deleterious recessive mutations do not show as dramatic of deviations from expectations as the case of weakly deleterious recessive mutations. The neutral SFS is reduced, and we observe more selected mutations at higher frequencies than expected. LD is also only weakly distorted from two-locus expectations. Notably, we see positive LD (matching between simulations and expectations) for strongly deleterious recessive mutations, as predicted in Roze (2021).

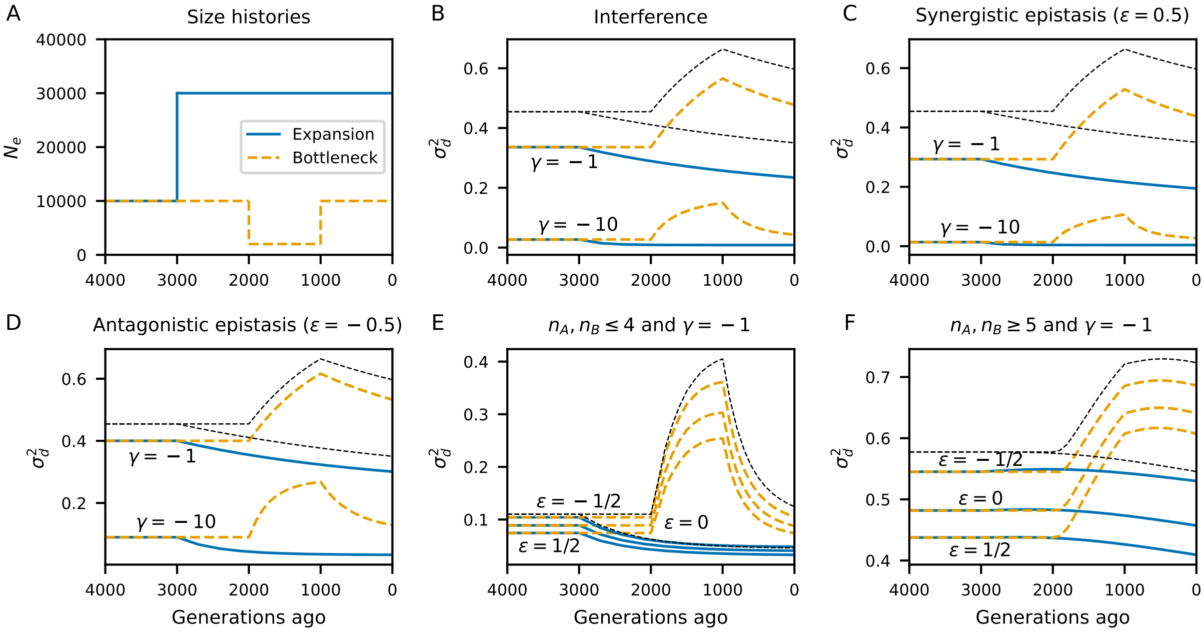

Figure S11: **Time series of squared LD under bottleneck and expansion histories with epistasis.** Bottlenecks increase $\sigma_d^2$ (squared LD), while expansions tend to decrease squared LD. Dashed lines show neutral expectations.

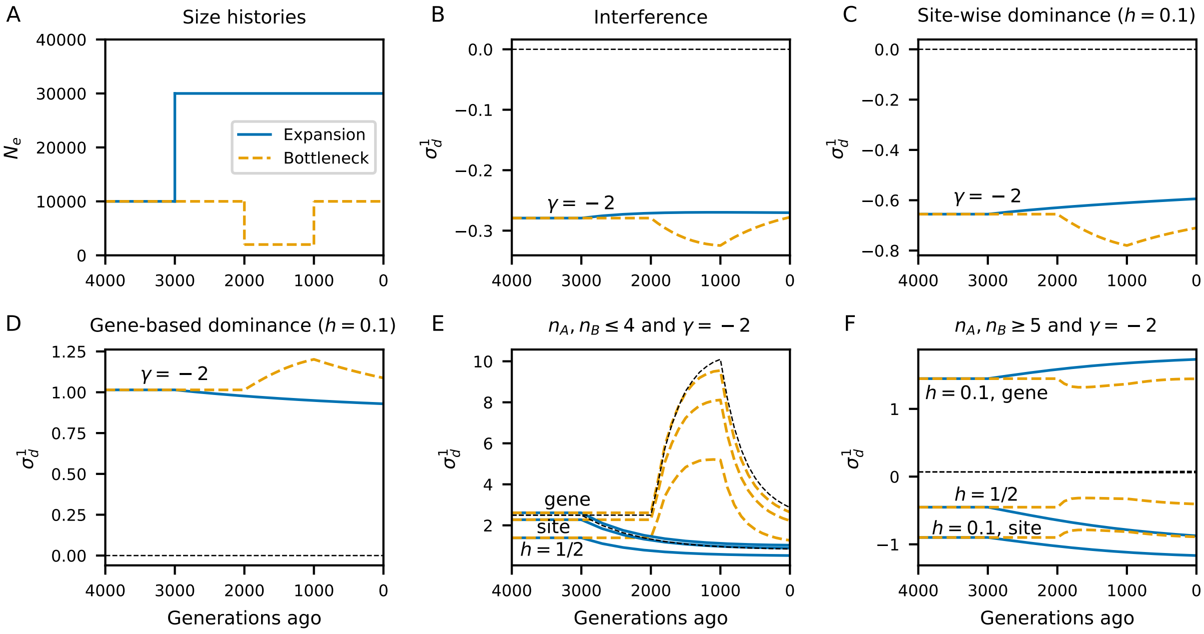

Figure S12: **Time series of signed LD under bottleneck and expansion histories with dominance.** Gene-based dominance, unlike site-wise dominance effects, can cause large positive signed LD, similar to compensatory mutations or antagonistic epistasis. In general, common variants are more stable under population size changes than rare variants. Dashed lines show neutral expectations.

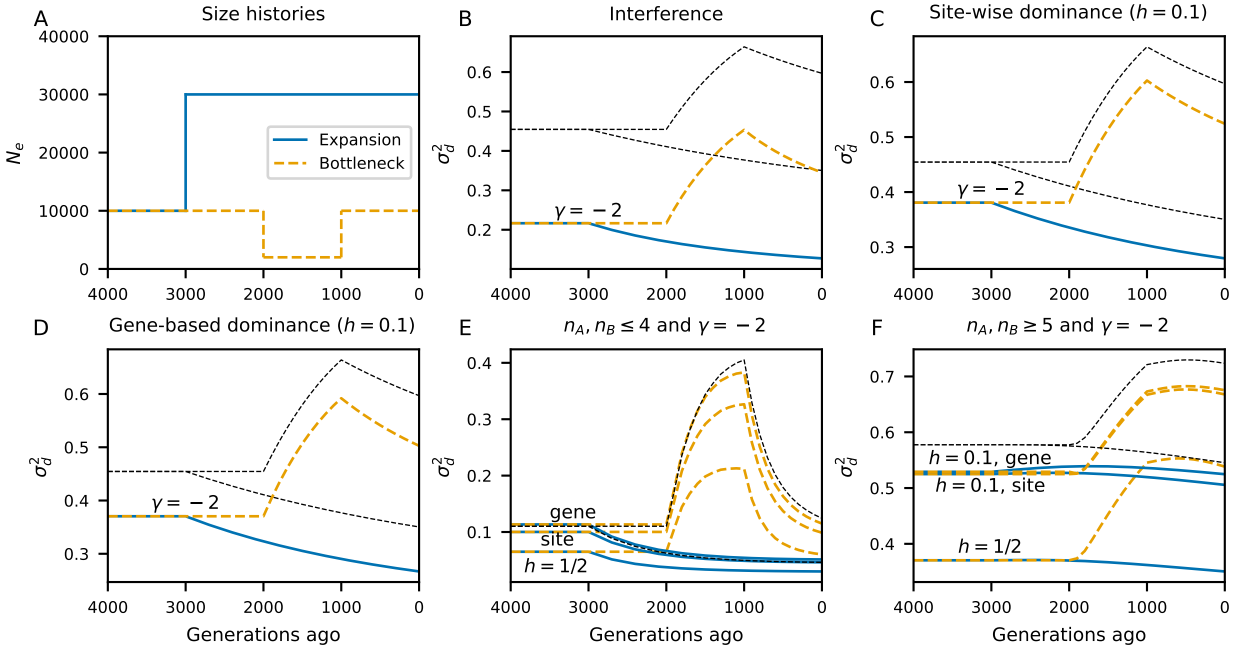

Figure S13: **Time series of squared LD under bottleneck and expansion histories with dominance.** Similar to the cases with epistasis, bottlenecks tend to increase $\sigma_d^2$ and expansions decrease squared LD. Dashed lines show neutral expectations.

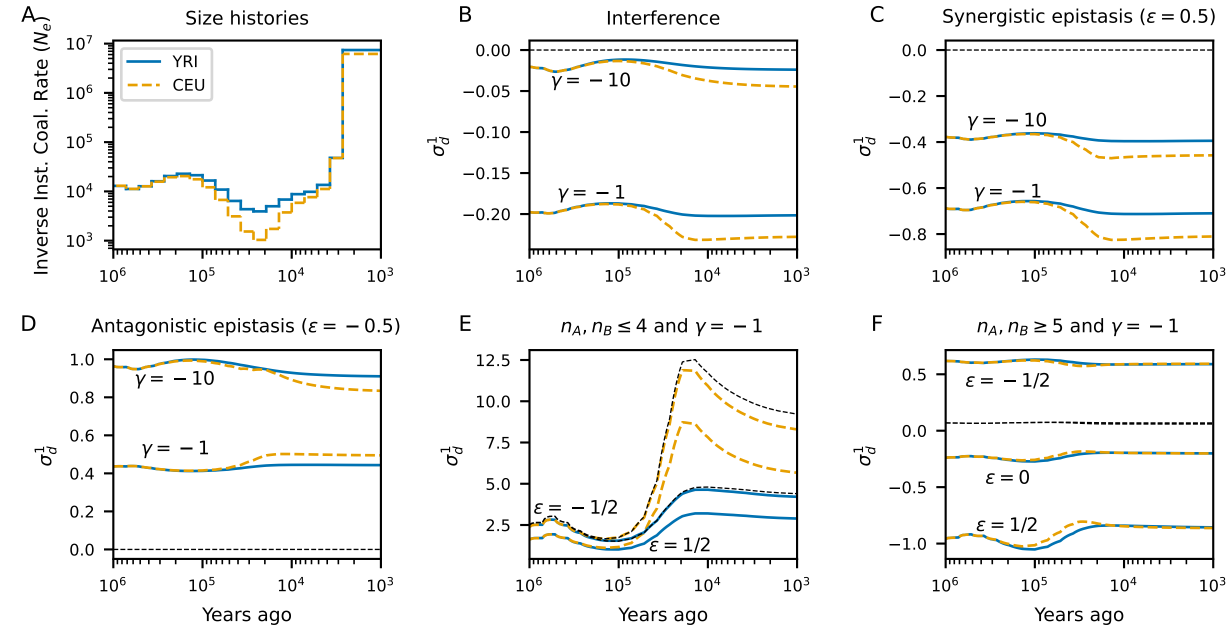

Figure S14: **Time series of signed LD under inferred models of human population size history with with epistasis.** Similar to the instantaneous size changes in the toy models (Figure 6), bottlenecks tend to make signed LD more extreme. However, in the case of strong selection and antagonistic epistasis, the CEU $\sigma_d^1$ trajector decreases relative to the YRI trajectory. Again, statistics for rare variants fluctuate more rapidly than common variants. Dashed lines show neutral expectations.

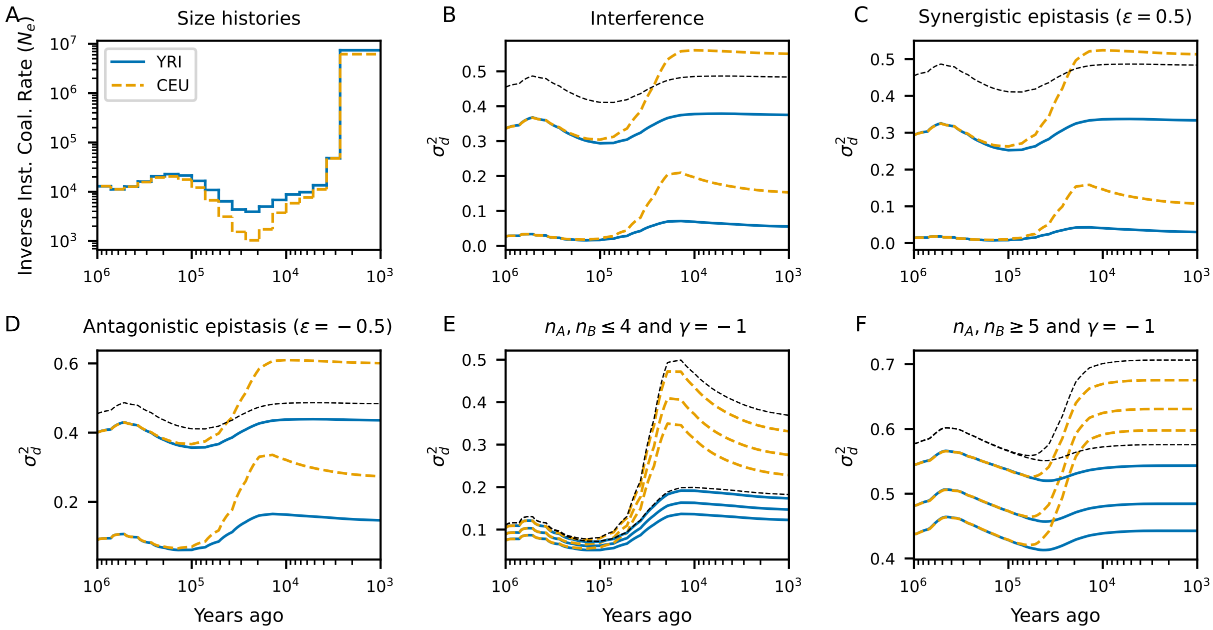

Figure S15: **Time series of squared LD under inferred models of human population size history with with epistasis.** Size reductions, such as bottlenecks in the recent past of Eurasian populations, increase squared LD. Dashed lines show neutral expectations.

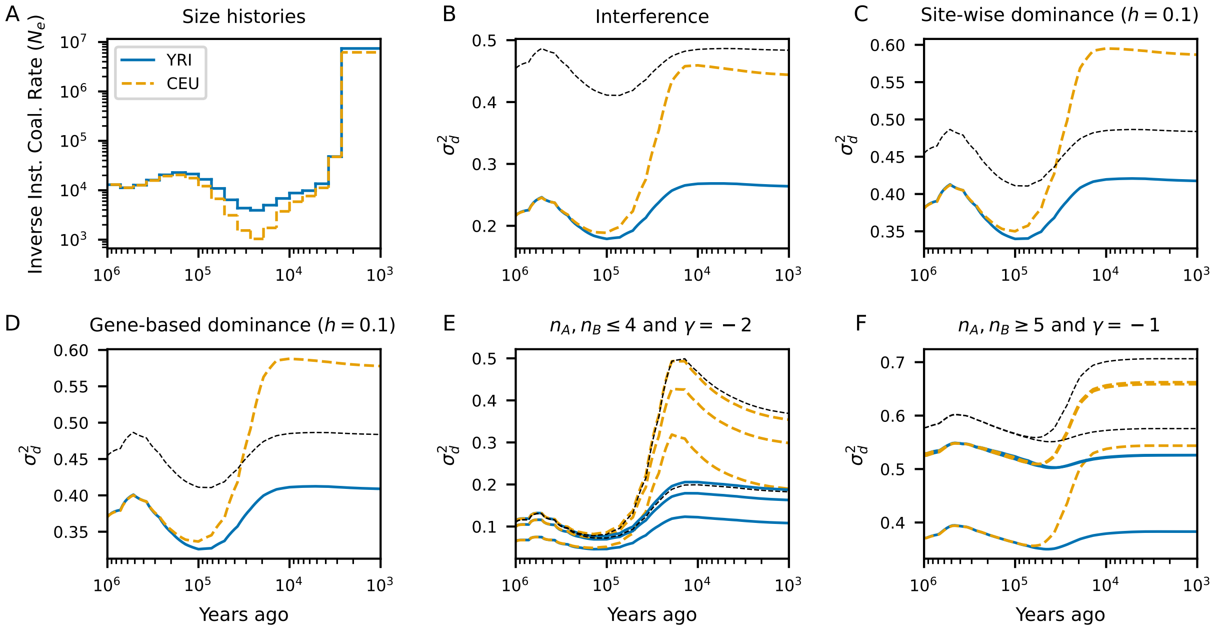

Figure S16: **Time series of squared LD under inferred models of human population size history with with dominance.** Again, bottlenecks increase squared LD relative to populations that did not experience such strong size reductions in their history. Dashed lines show neutral expectations.

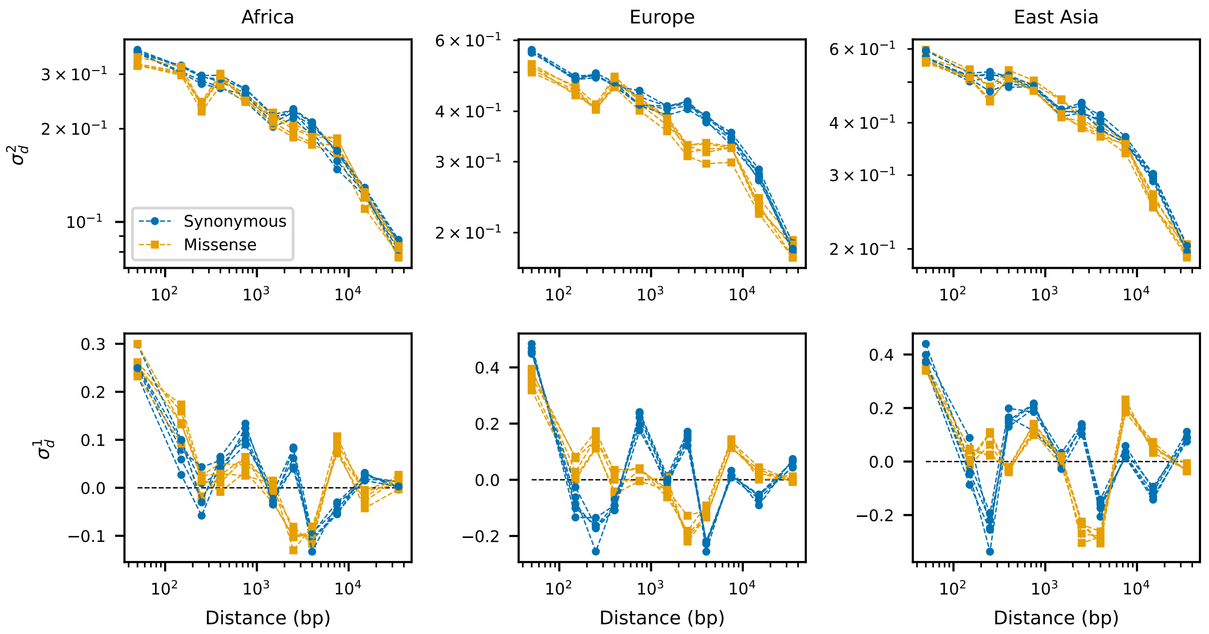

Figure S17: **LD decay for pairs of synonymous and missense mutations in the same gene.** When taking gene-wide averages of LD, there are no obvious differences between the LD decay curves of synonymous and missense mutations. Each panel contains five LD decay curves for each mutation class, one for each 1000 Genomes Project Consortium *et al.* (2015) population in the continental groupings. Shared demographic history leads to strong correlation in statistics between populations in the same continental groups.

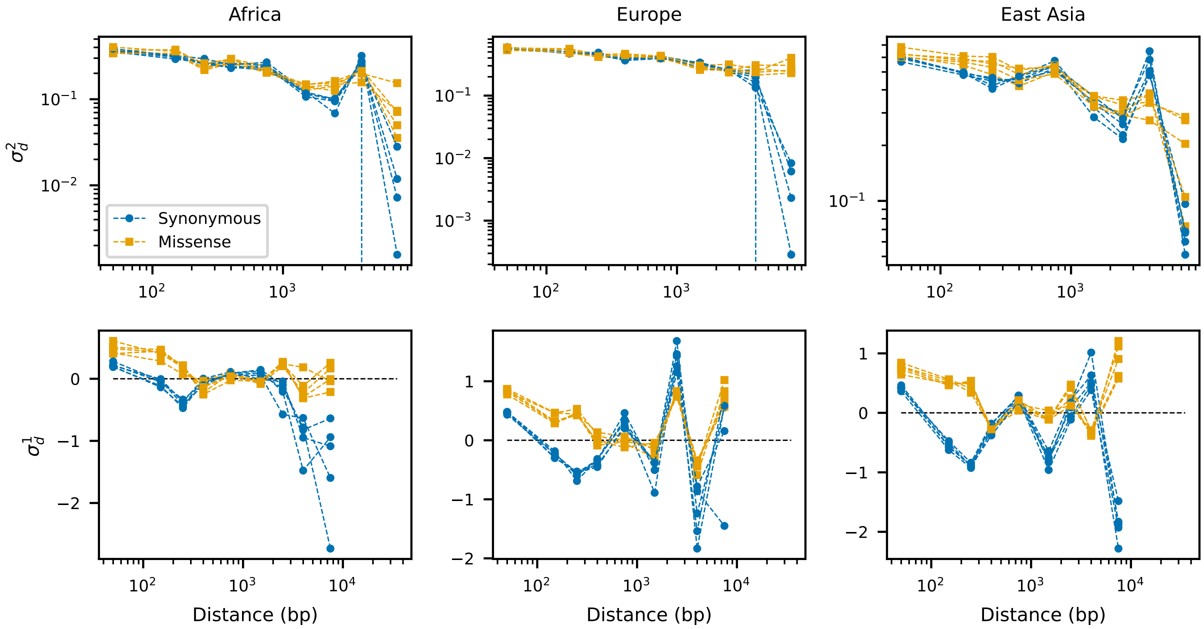

Figure S18: **LD decay for pairs of synonymous and missense mutations that fall inside the same domain.** At short distances between SNPs that both fall within sthe same conserved annotated domain, missense mtuations show consistently larger signed LD than synonymous variants. This may be caused by antagonistic epistasis or compensatory effects between pairs of nearby missense mutations.

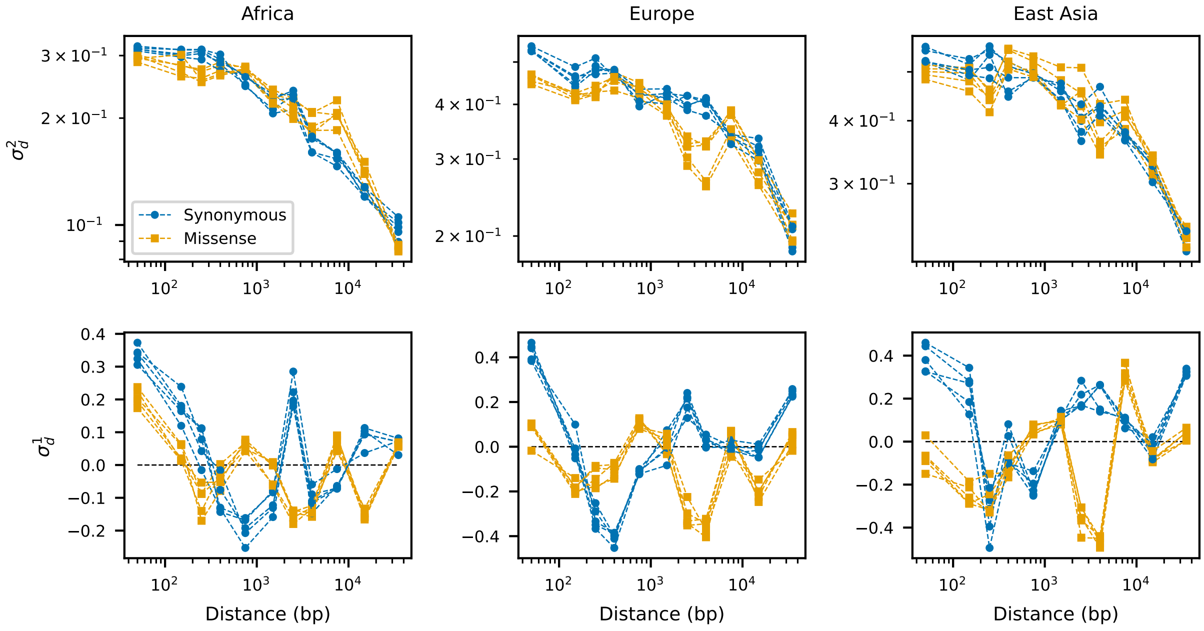

Figure S19: **LD decay for pairs of synonymous and missense mutations outside of domains.** Outside of conserved domains, missense mutations have reduced signed LD at short distances compared to synonymous mutations, opposite the pattern observed within domains (Figure S18). This pattern is consistent with Hill-Robertson interference between missense mutations outside of conserved elements.

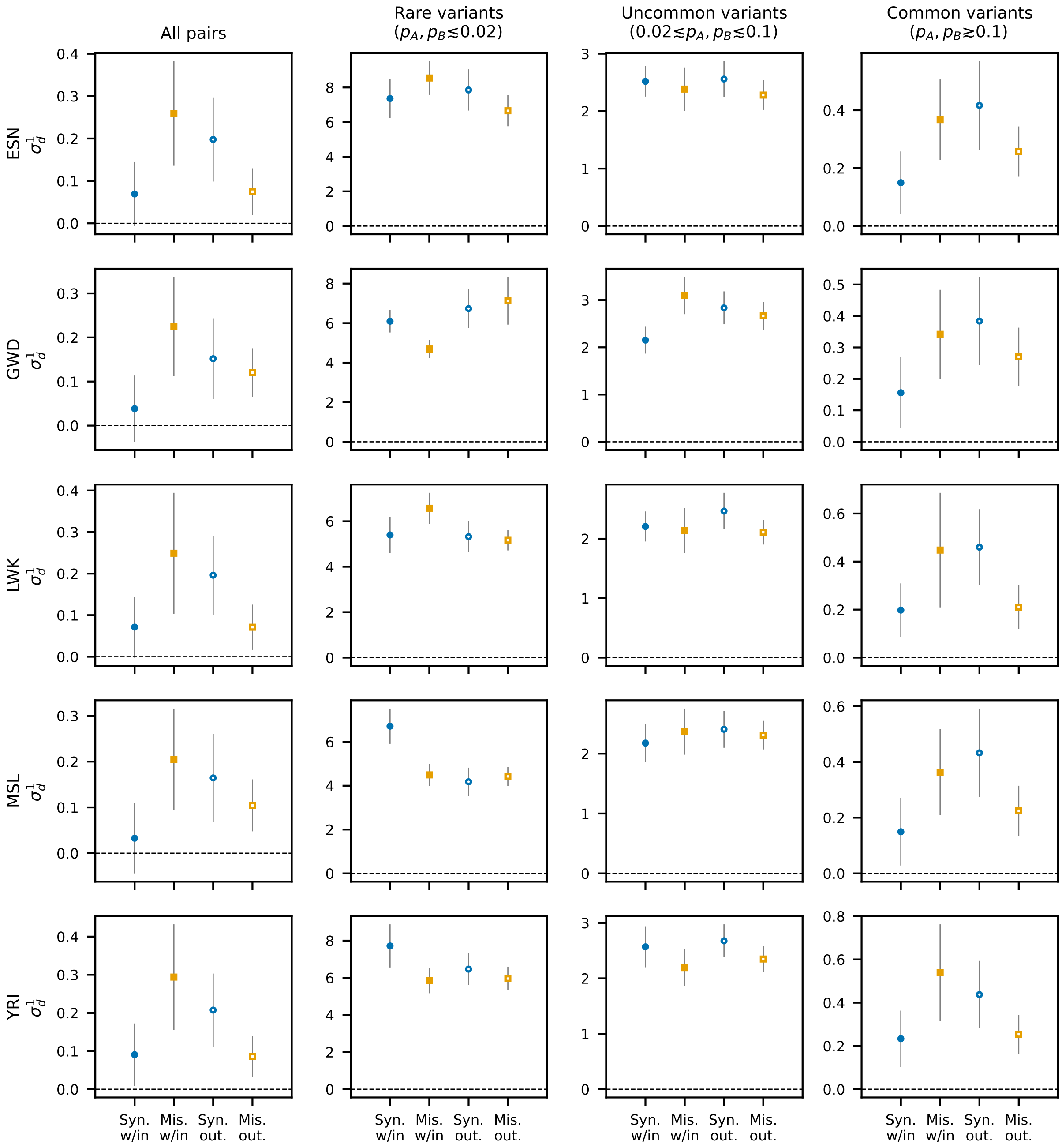

Figure S20: **LD for pairs of synonymous and missense mutations within the same domain and outside domains at matched distances.** Showing the five populations in the Thousand Genomes African group. We observe increased LD between missense mutations within domains, but decrased LD outside of domains at matched distances. This pattern holds when conditioning on non-rare variation ($n_A, n_B \geq 9$, MAF $\gtrsim 0.05$).

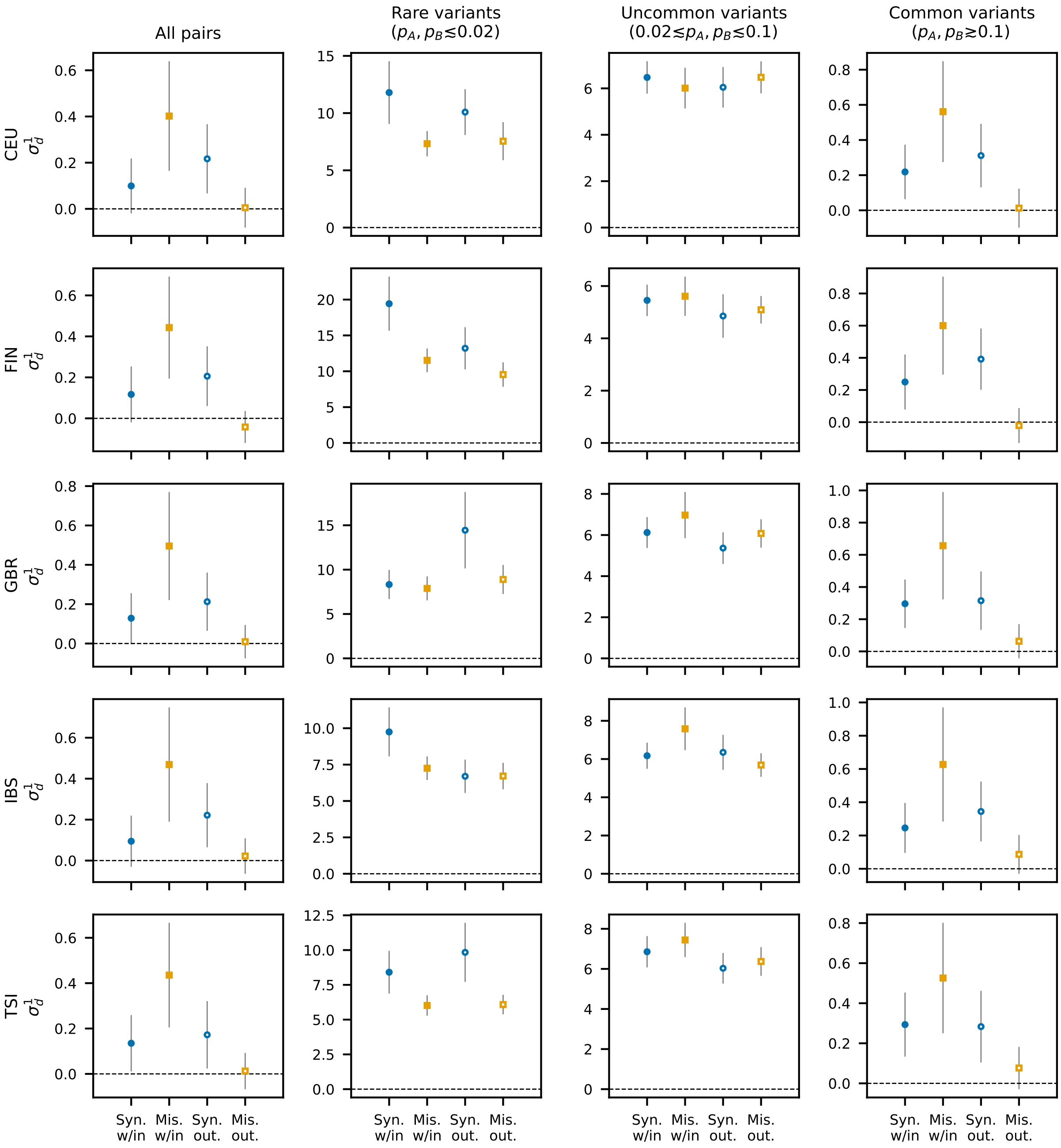

Figure S21: **LD for pairs of synonymous and missense mutations within the same domain and outside domains at matched distances.** Showing the five populations in the Thousand Genomes European group. We again observe increased LD between missense mutations within domains, but decrased LD outside of domains at matched distances. And again, this pattern holds when conditioning on non-rare variation ($n_A, n_B \geq 9$, MAF $\gtrsim 0.05$).

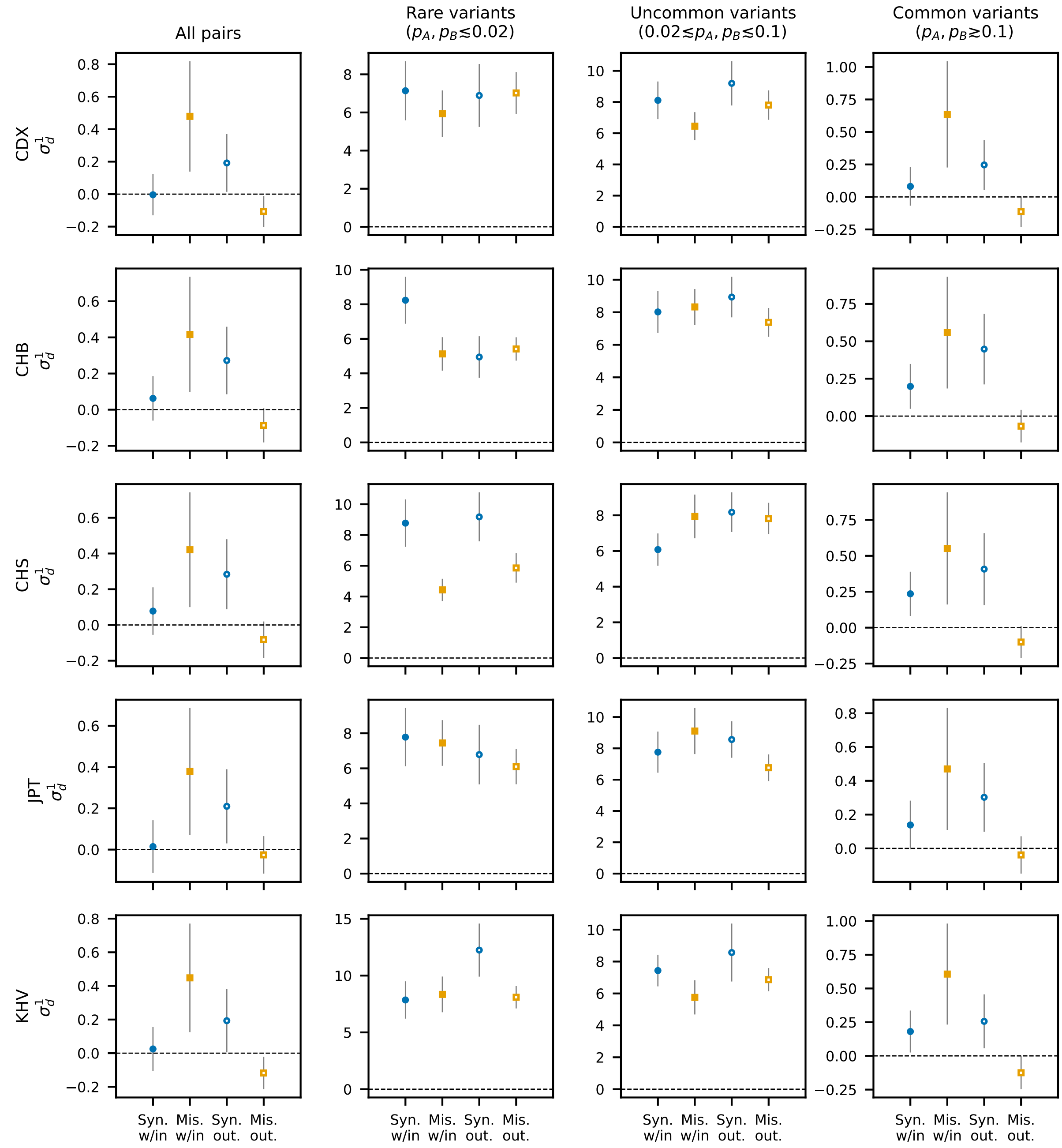

Figure S22: **LD for pairs of synonymous and missense mutations within the same domain and outside domains at matched distances.** Showing the five populations in the Thousand Genomes East Asian group. As with the African and European data, wobserve increased LD between missense mutations within domains, but decrased LD outside of domains at matched distances. And again, this pattern holds when conditioning on non-rare variation ($n_A, n_B \geq 9$, MAF $\gtrsim 0.05$).

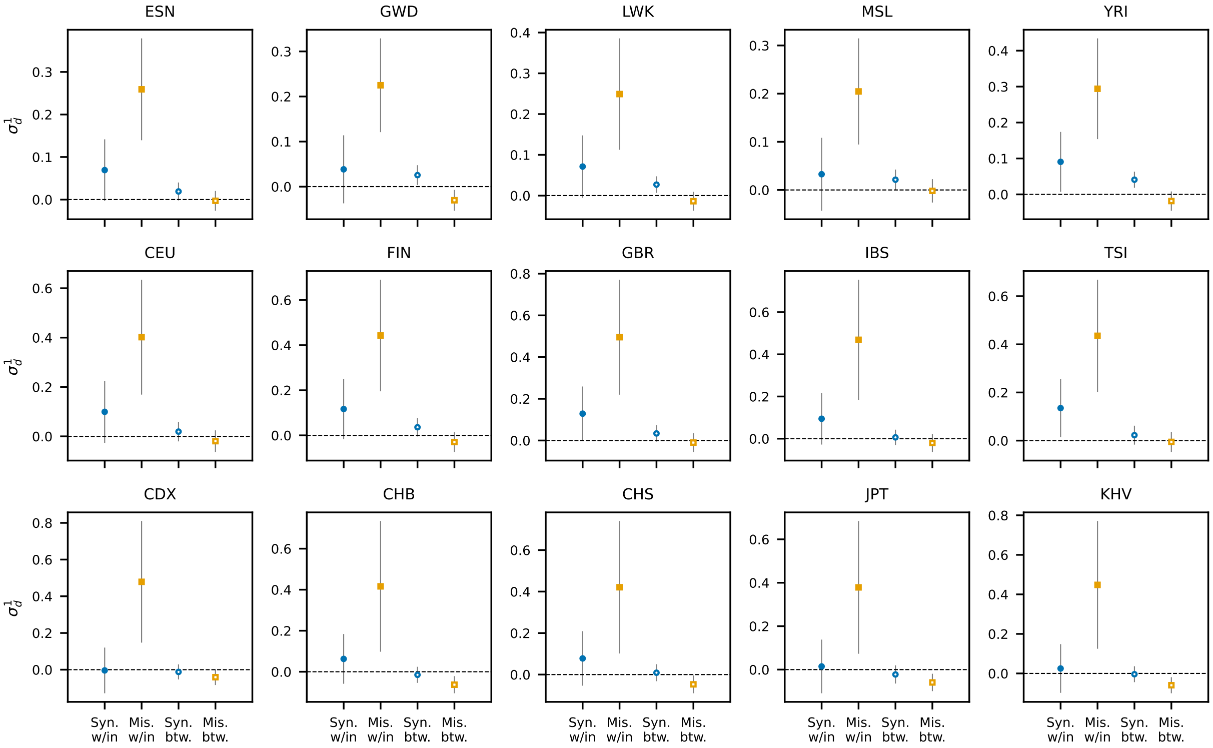

Figure S23: **LD for pairs of synonymous and missense mutations both within the same domain and in differing domains.** In each panel (one for each population), the left two data points show signed LD between mutations that fall within the same domain, and the right two data points show signed LD between mutations falling within domains but in *different* domains within the same protein-coding gene. Distances are not matched between these data classes (as between-domain pairs are typically more distant than within-domain pairs). Increased LD between missense mutations is only observed for pairs of mutations within the same domain, but not between domains.

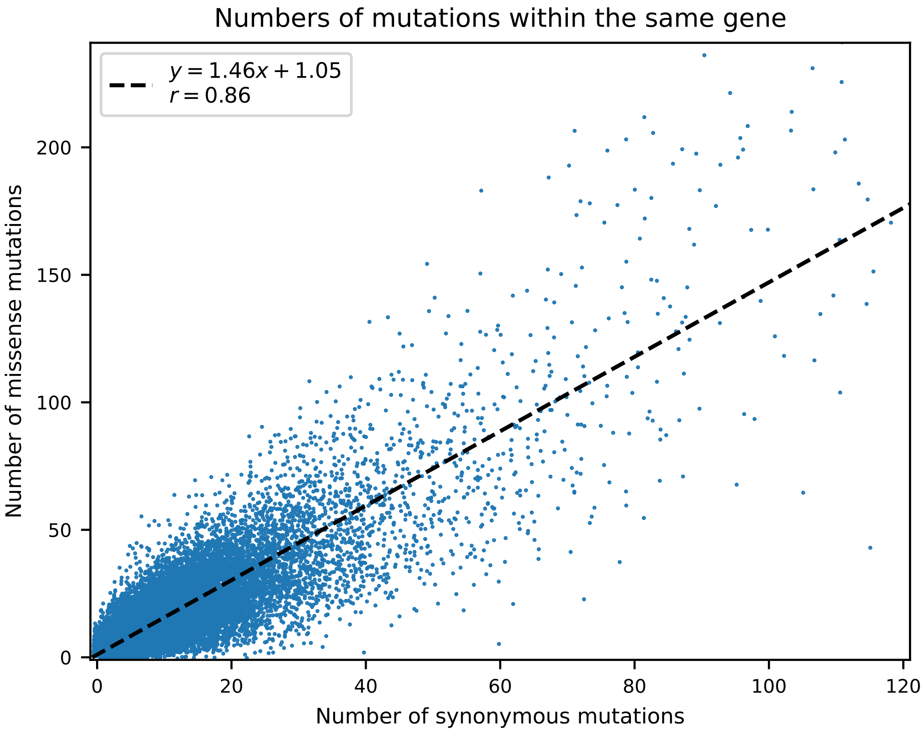

Figure S24: **Correlation between numbers of missense and synonymous mutations within each gene.** Human genes have different sizes and sequence content, so they may differ in the numbers of missense and synonymous mutations observed within them. The number of missense mutations within a gene is predicted well by the number of synonymous mutations ($r = 0.86$).

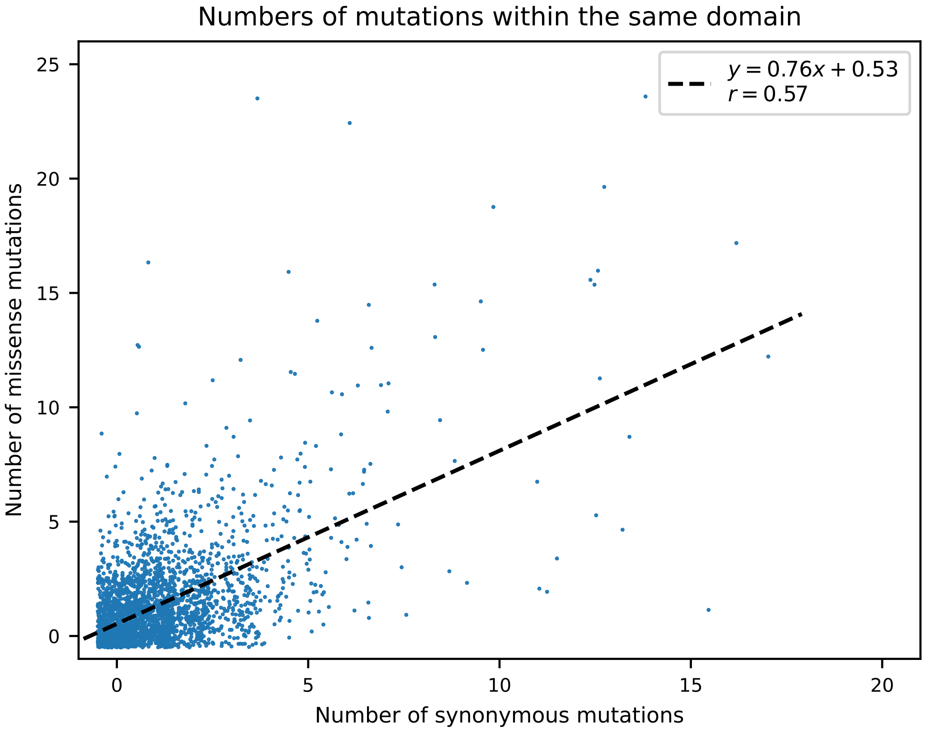

Figure S25: **Correlation between numbers of missense and synonymous mutations within each annotated domain.** Domains are typically much smaller than the average gene-size, and while the number of synonymous mutations within a domain is moderately predictive of the number of missense mutations within the same domain ($r = 0.57$), this correlation is weaker than the correlation in gene-wide mutation counts (Figure S24).

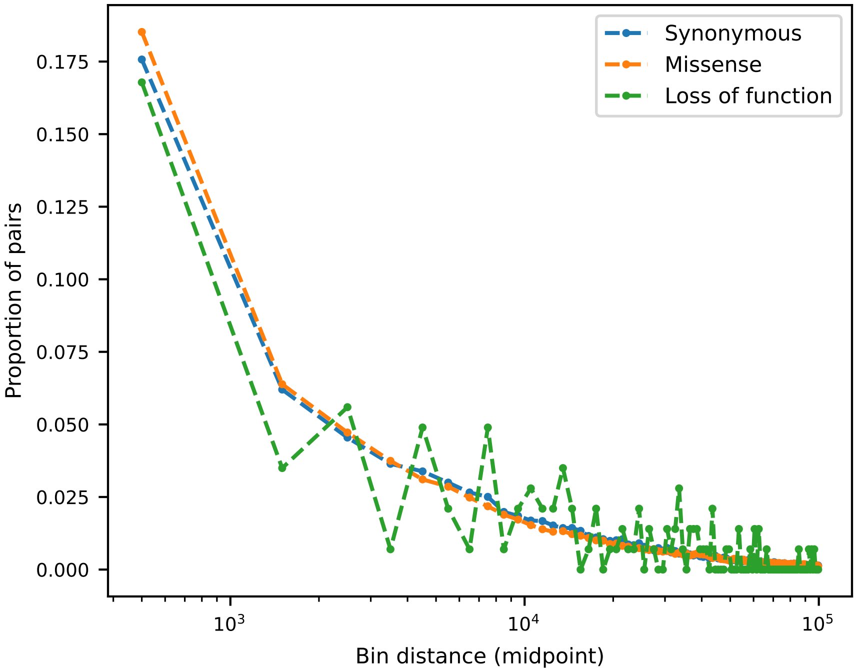

Figure S26: **Distances between mutation pairs for different classes of variants.** Despite different mutation rates between classes of mutations, the proportions of pairs of mutations at given distances are similar. Missense mutations, which have the highest mutation rate within genes, have slightly more pairs of mutations at close distances than synonymous, with loss of function mutations, which have the lowest mutation rate within genes, have the fewest pairs at short distances.

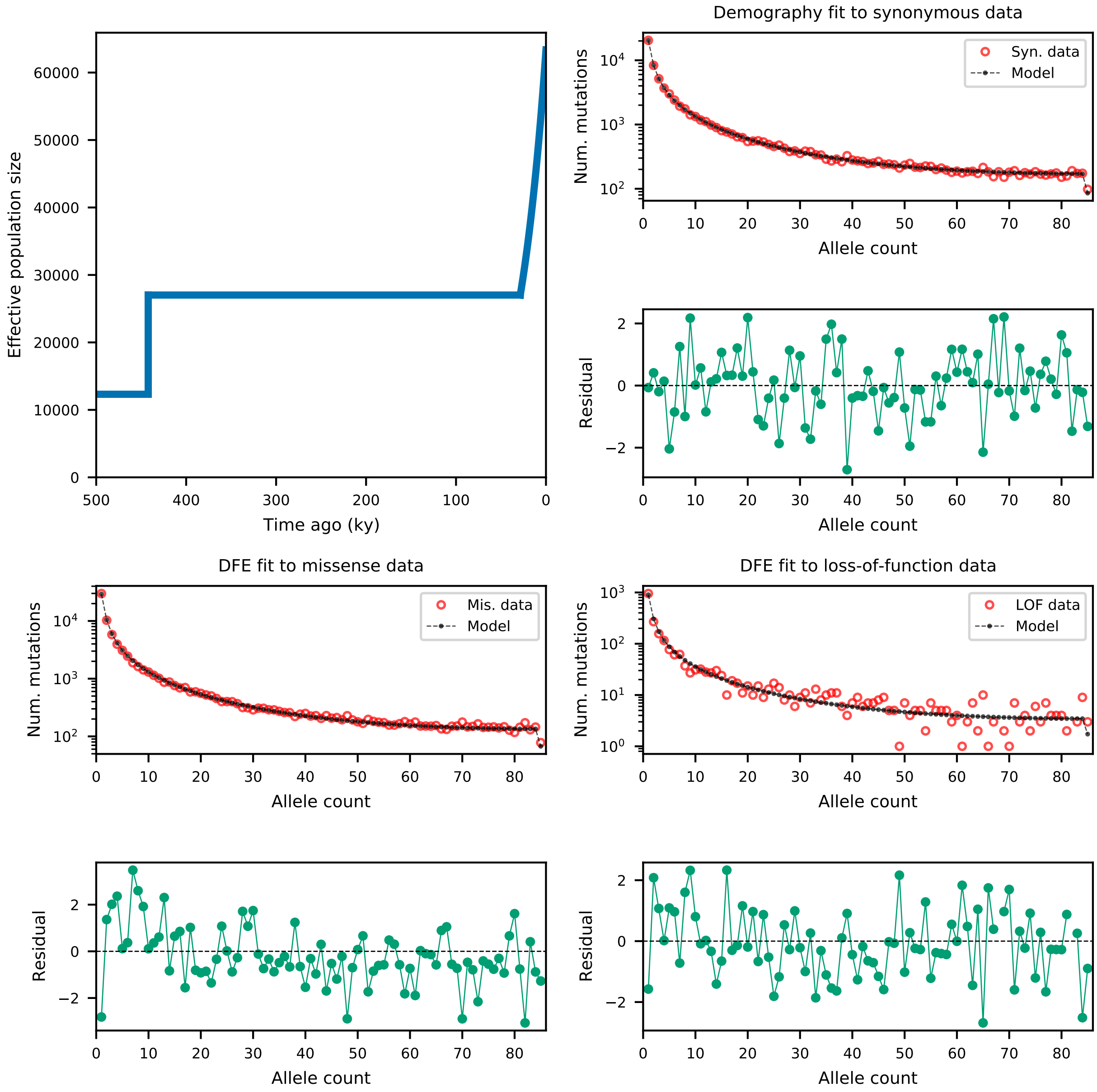

Figure S27: **Demography and DFE for MSL.** A demographic model was fit to the folded synonymous SFS, and DFEs were fit to missense and loss-of-function SFS. Shown here are DFEs fit with $h = 0.5$. See Table S5 for inferred best-fit parameters.

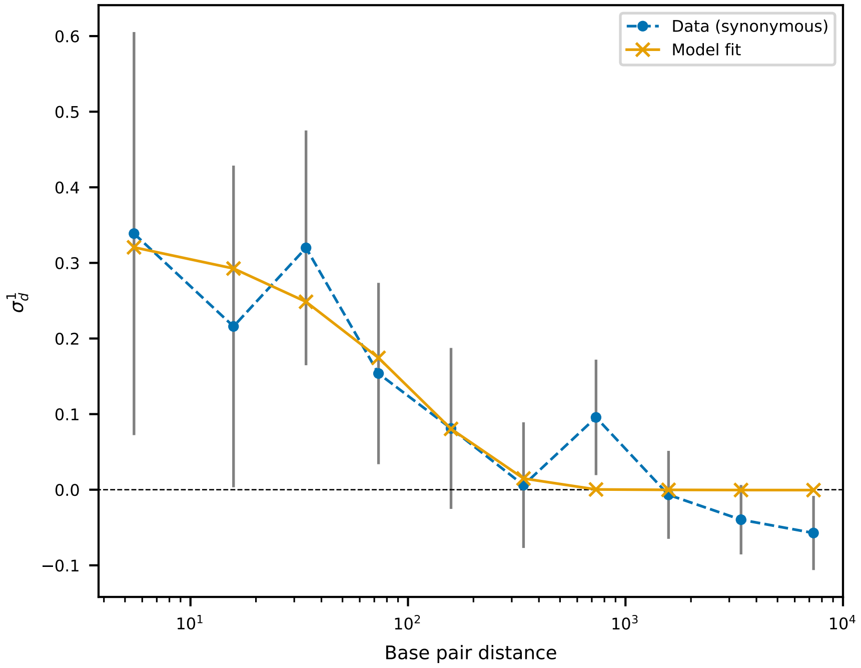

Figure S28: **Optimization of fraction of new mutations arising via multinucleotide mutations by distance.** A simple exponential function was fit to LD decay of synonymous mutations, to describe the probability that a given mutation event is a multinucleotide mutation event that spontaneously gives rise to a pair of perfectly linked mutations at count $1/2N$. Fitting $Ae^{-\lambda d}$ to mutations (see Section S2.2), the best fit parameters across all recombination rates tested were $A = 2.4 \times 10^{-5}$ and $\lambda = 0.0092$, showing that even small rates of multinucleotide mutation events can cause increased signed LD at short distances.
